# Synthetic chaperone based on Hsp90-Tau interaction inhibits pathological Tau aggregation and rescues physiological Tau-Microtubule interaction

**DOI:** 10.1101/2024.10.01.615850

**Authors:** Davide Di Lorenzo, Nicolo Bisi, Julia Kaffy, Lisa Marie Ramirez, Markus Zweckstetter, Olivier Lequin, Irene Garfagnini, Jinghui Luo, Yvonne Hannappel, Inga Ennen, Veronica Dodero, Norbert Sewald, Maria Luisa Gelmi, Nicolo Tonali, Roland Brandt, Sandrine Ongeri

## Abstract

The accumulation of intracellular aggregates of Tau protein is one main hallmark of Alzheimer’s disease (AD) and is the consequence of Tau conformational changes, increased phosphorylation, and self-association to form fibrillar aggregates. This pathological process prevents the physiological interaction of Tau with microtubules to the detriment of the structural integrity of neurons. In healthy cells, aberrant protein misfolding and aggregation are counteracted by chaperone proteins whose protective capacity decreases with age. The role of the chaperone Hsp90 and the mechanism by which it can prevent Tau aggregation toxicity are controversial. The innovative strategy of mimicking Hsp90 through the design of the β-hairpin like peptidomimetic **β-Hsp90**, inspired by two Hsp90/Tau interaction sequences, is presented here. **β-Hsp90** inhibits Tau aggregation both *in vitro* and *in cells*, restoring Tau’s physiological interaction with microtubules. **β-Hsp90**, which interacts with the P1 region of Tau, is more effective than individual peptide sequences from the chaperone HSP90 and another β-hairpin mimic based on Tau sequences. Moreover, **β-Hsp90** dramatically reduces AD-associated Aβ_1-42_ aggregation, offering the development of a dual inhibitor. This work paves the way for the design of new drugs targeting devastating untreated amyloid diseases, by mimicking physiological chaperones with small synthetic peptide drugs.

---

The accumulation of protein deposits in the brain is a common hallmark of many neurodegenerative pathologies, including Alzheimer’s (AD), Parkinson’s (PD), and Huntington’s diseases (HD), among others.^1^ The misfolding and aggregation of these proteins (amyloid-β (Aβ) and Tau in AD, α-synuclein in PD, and Huntingtin in HD)^2–4^ lead either to the formation of toxic species or to the depletion of a functional protein which plays an important physiological role in the homeostasis of neuronal cells. The incidence of these pathologies, called amyloid diseases, increases with the aging of the population and constitutes a major threat to all health systems. AD is the most prevalent devastating neurodegenerative disorder and the most prevalent form of tauopathies. Tau^5–7^ is a member of the microtubule-associated protein (MAP) family and its physiological role is to promote the assembly and dynamics of microtubules (MTs), which are essential for axonal transport and maintenance of structural integrity of neurons.^8,9^ Under pathological conditions, Tau monomers undergo conformational changes and increased phosphorylation at selected sites, which are associated with the development of abnormal soluble oligomers and filamentous aggregates, which are unanimously considered to cause degeneration of neuronal and glial cells.^5,8^ AD is also characterized by the formation of other toxic amyloid aggregates composed mainly of Aβ_1-42_. Many efforts have been made to develop a treatment for AD, mainly with single-target drugs (Tau or Aβ_1-42_), but unfortunately, most of them have failed. To date, no drug targeting Tau has been approved for the therapeutic treatment of AD. However, despite their controversial utility and high cost, three antibodies, Aducanumab, Lecanemab,^10^ and Donanemab (July 2024), targeting Aβ_1-42_, have recently arrived on the market following accelerated FDA approval, ushering in the last three decades of the amyloid hypothesis.

A hitherto unexplored strategy for combating AD, as well as other amyloidoses, consists in drawing inspiration from chaperone proteins to design effective inhibitors against pathological aggregation processes. Indeed, in healthy cells, aberrant protein misfolding and aggregation are counteracted by a group of proteins called molecular chaperones, considered to be part of the cellular quality control machinery.^11–13^ Neurons rely on molecular chaperones to cope with misfolded protein levels throughout their lifespan. However, with age, the protective capacity of molecular chaperones decreases and becomes less responsive to stress signals, allowing misfolded protein species to accumulate and eventually aggregate.^14,15^ Until now, how molecular chaperones defend neurons against the accumulation of misfolded proteins remains unclear.^16^ Tau and Aβ (as well as many other amyloid proteins) are hybrid proteins that possess both structured domains and intrinsically disordered protein (IDP) regions.^17^ Chaperones interacting with IDPs could exhibit a holdase mechanism, which involves capturing free unfolded proteins and maintaining them in soluble forms.^18^ However, full-length physiological chaperones cannot be effectively used as drugs, mainly due to pharmacokinetic and cost issues. We considered the innovative strategy to increase the proteostasis capacity by mimicking natural protein chaperones with small peptide drugs to combat tauopathies. To our knowledge, the strategy of designing small peptides derived from natural chaperone sequences has only been reported to prevent Aβ aggregation, one using a 19-residue peptide based on αA-crystallin,^19^ and the other developed by some of us using diaza-nonapeptides based on Thransthyretin.^20^ However, this strategy has not been investigated for tauopathies and is the focus of this promising new study presented here.

The heat shock protein 90 kDa (Hsp90) chaperone particularly attracted our attention because the role of Hsp90 in neurodegeneration is rather controversial. In particular, the exact activities and molecular mechanism of Hsp90-Tau complex remain enigmatic. Regarding the activity of Hsp90, its interaction with Tau appears to either promote or protect Tau against degradation and toxic aggregation, depending in particular on the associated co-chaperone.^21–25^ Hsp90 alone has been shown to inhibit the formation of Tau fibrils but to promote the formation of small Tau oligomers.^26,27^ In complex with the *cis*–*trans* peptidyl-prolyl isomerase FKBP51, Hsp90 also acts to promote Tau oligomers formation.^28^ On the contrary, the Hsp70/Hsp90 machinery counteracts the aggregation of the highly affine phosphorylated Tau.^29^ Regarding the binding sites of Tau to Hsp90, they are reported as highly polymorphic. A direct interaction of Hsp90 with Tau repeat domain (Tau-RD) has been mainly reported, while this region is also involved in the binding of Tau to MTs (Tau-RD is also called microtubule-binding region, MTBR, Fig. 1A) and in its self-aggregation.^30,31^ However, in the ternary Hsp90/FKBP51/Tau complex an interaction of the proline-rich region P1 of Tau with the Hsp90 *N-*domain was observed.^28^ Phosphorylated Tau has been also reported to interact with the Hsp70/Hsp90 complex at its proline-rich region P2 and its MTBR flanking pseudo-repeat R’.^29^ These observations suggest that Hsp90 can act by interacting through the MT-binding domain to favour aggregates that might be less toxic than oligomers and fibrils, and that this Hsp90/Tau complex might prevent deleterious interactions of Tau aggregates with cytoplasmic proteins. In multi-complex systems, Hsp90 rather interacts with the proline-rich region P1/P2 and R’ flanking Tau-RD to prevent or increase Tau oligomerization. The question of whether the Hsp90-Tau interaction allows to preserve the physiological interaction between Tau MTBR and MT has not yet been addressed.

**Fig. 1:**
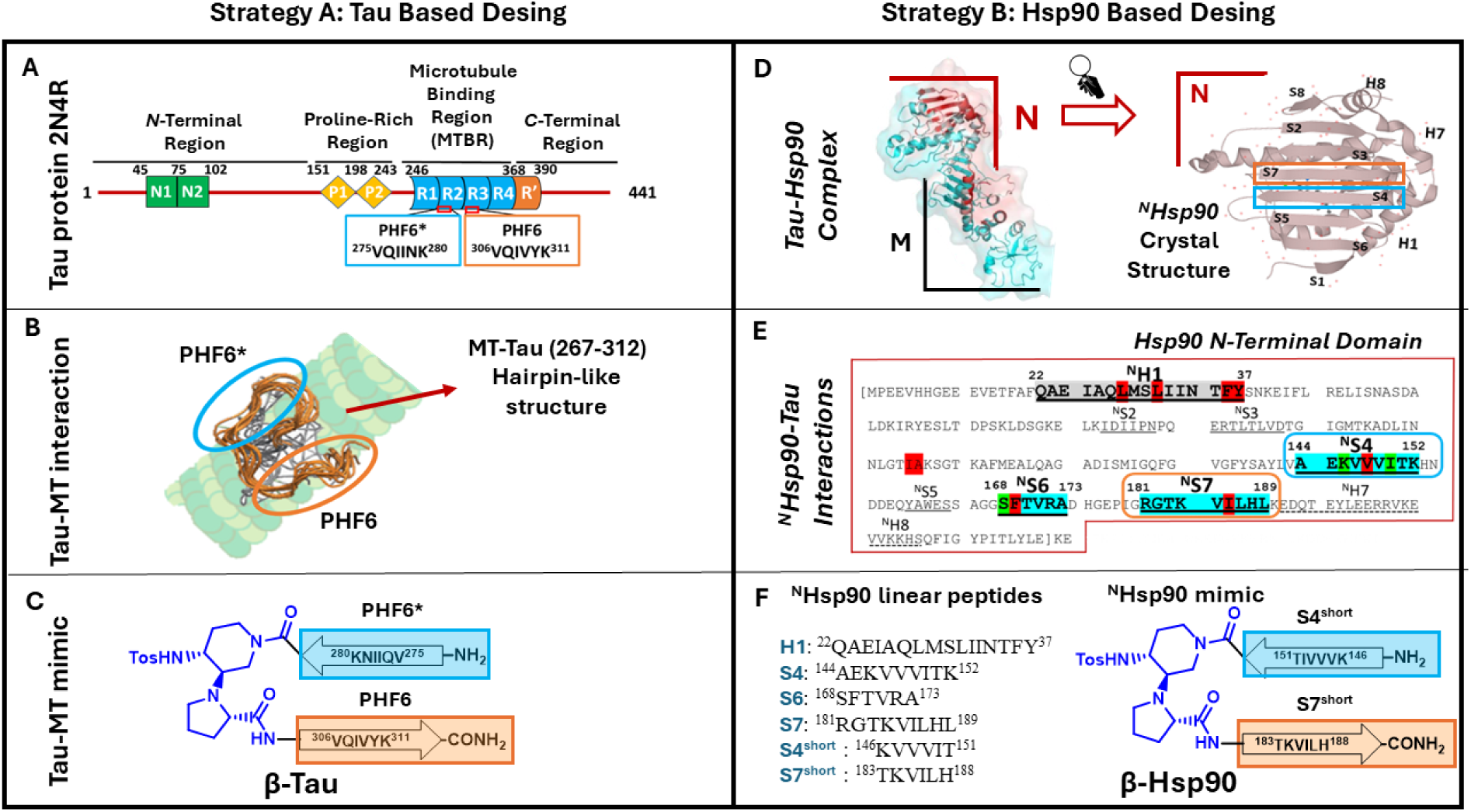
**A)** Structure of Tau: schematic representation of Tau longest CNS isoform; **B)** Schematic representation of Tau folded structure when interacting with MT^6^ ; **C)** Chemical representation of Tau-MTs β-hairpin mimic compound (**β-Tau**); **D)** Crystal structure of the Hsp90 *N*-terminal domain in complex with EC44 (PDB = 3NMQ) and schematic representation of the spatial distributions of β-strands placed at the crystal structure of the Hsp90 *N*-terminus (M = middle domain, N = N-terminal domain); **E)** Primary structure of the Hsp90 *N*-terminal region highlighting key interaction sites according to Karagöz *et al.*^30^: β-strands in blue, helical structures in grey, strong hydrophobic interactions in red, and significant mono-amino acid interactions in green. **F)** Chemical depiction of the designed peptides (**H1**, **S4**, **S6**, **S7**, **S4^short^**, **S7^short^**) and β-hairpin mimics **β-Hsp90** inspired by the Tau-^N^Hsp90 interaction.

Interestingly, for AD, although the Hsp90/Aβ complex has not been studied much, Hsp90 has also been described to be able to inhibit Aβ aggregation.^32,33^

Given these considerations, the strategy of mimicking the Hsp90 chaperone activity through small peptidomimetics inspired by Hsp90/Tau interaction sequences, is here presented. These new compounds have been employed both as exploratory tools to provide some useful insights into the mechanism and physiological role of Hsp90 on Tau, and to achieve efficient inhibitors of Tau aggregation able to restore physiological Tau-MT interaction. A flexible β-hairpin mimic, **β-Hsp90**, built on a piperidine-pyrrolidine β-turn inducer and bearing defined sequences from Hsp90, demonstrated to be a strong inhibitor of both wild-type Tau (Wt-Tau) and pro-aggregative ΔK280 Tau mutant fibrillization, as assessed by Thioflavin-T (ThT) fluorescence spectroscopy. This design was compared to the canonical self-recognition elements (SREs) approach based on specific Tau sequences (compound **β-Tau**). Transmission electron microscopy (TEM) images gave information about the morphology of Tau species in the presence of these new Tau aggregation inhibitors. Then, through live cell imaging FDAP (fluorescence decay after photoactivation), the anti-aggregative property of compound **β-Hsp90** was confirmed in a cellular context, also highlighting its ability to restore Tau-MT interaction to physiological levels. The unprecedented involvement of the P1 region of Tau in the interaction with this efficient compound was proven thanks to ^15^N-HSQC titrations of ^15^N-Tau. Furthermore, the ability of **β-Hsp90** to act as a dual inhibitor of Tau and Aβ_1-42_ aggregation was shown by ThT and TEM experiments conducted on Aβ_1-42_. These results open new prospect for future research in the field of Alzheimer’s therapy, considering that combination therapies and dual targeting molecules, such as dual inhibitors of Tau and Aβ aggregation, have already been suggested as promising therapeutic strategies, with respect to single-target approaches.^34^

## Results

### Designing β-hairpin mimics to interact with Tau and inhibit its aggregation

The effectiveness of rationally designed acyclic β-hairpin mimics, built on a piperidine-pyrrolidine β-turn inducer and bearing two small peptides based on SRE sequences involved in the misfolding of Aβ_1-42_ or hIAPP, in selectively inhibiting either Aβ_1-42_ or hIAPP aggregation processes, has already been shown by some of us.^35–37^ For Tau, the aggregative SRE sequences are essentially localized in the 306-378 region located in the MTBR,^38^ while in physiological conditions, this sequence actively participates in the binding of Tau with MTs. Thus, in a first approach, a selective peptidomimetic targeting Tau was designed taking inspiration from the secondary structure and self-interactions that Tau engages when it is bound to MTs,^6^ with the objective to stabilize this conformation and delay the misfolding towards pro-aggregative conformations. However, we were aware of the risk that mimicking this interaction zone could compete and compromise the physiological interaction of Tau with MTs. By exploring the conformation that Tau adopts when binding to MTs using NMR analysis, Kadavath *et al.* noticed that the two hexapeptide sequences, PHF6* (^275^VQIINK^280^ at the beginning of R2) and PHF6 (^306^VQIVYK^3^^11^ at the beginning of R3. Fig. 1A), form a hairpin-like structure allowing the stabilization of the Tau-MT interaction (Fig. 1B).^6^ Noteworthy, these sequences are also known to be the “hot spots” of Tau self-association, spontaneously aggregating in solution into β-sheet structures and triggering both the oligomerization and the fibril growth.^39,40^ Therefore, considering that the hairpin conformation is stabilized during the interaction with MTs and that these two sequences are possibly hidden into the globular form of the full-length protein, thereby reducing their solvent exposure and their aggregation propensity, the two PHF6 and PHF6* sequences have been inserted into the piperidine-pyrrolidine synthetic β-turn inducer to mimic this kind of β-hairpin. Our hypothesis is that this pre-organized conformation might increase the interaction with Tau and behave as a conformational pillar able to induce the full-length Tau (Tau^FL^) protein to adopt and maintain its physiological shape for its interaction with MTs (compound named **β-Tau**, **Strategy A**, Fig. 1C).

Secondly, given the intriguing role of the chaperone machinery, notably Hsp90, in regulating Tau misfolding and aggregation, our second design approach focused on mimicking the Hsp90-Tau interaction. This novel synthetic chaperone was hypothesized to reduce Tau fibrillization and it might help to clarify whether the reported promotion of oligomerization exerted by Hsp90 on Tau is detrimental or beneficial for its interaction with MTs in a cellular context. Thus, the sequences of Hsp90 suspected to be the recognition elements for its interaction with Tau were identified and inserted in a β-hairpin template, to mimic the core element of the interaction. The choice of these sequences referred to a comprehensive study employing Small-Angle X-ray Scattering (SAXS) and Nuclear Magnetic Resonance (NMR) spectroscopy to elucidate structural details of the Tau-Hsp90 complex (Fig. 1D).^30^ It is, however, essential to emphasize that, although this study disclosed the most prominent interacting regions (MTBR for Tau and *N*-terminal and central domains of Hsp90), the exact interacting residues involved in both proteins remain elusive, limiting the design of Hsp90-based inhibitors. To bridge this gap, the aforementioned study^30^ has been combined to a mapping of the crystallographic structure of the *N*-terminal domain of Hsp90 (^N^Hsp90; PDB = 3NMQ), highlighting key interacting regions (helices, β-sheets) and single amino acids hot spots (negative charges, hydrophobic sites and single interactions) within the Hsp90-Tau complex (Fig. 1E, **Strategy B**, Right side). This comprehensive analysis facilitated localizing the most relevant interacting regions and possible binding sites of ^N^Hsp90 with Tau (Fig. 1D and 1E), leading to the identification of four sequences suspected to play a pivotal role in the interaction (**^N^Helix 1** as compound **H1**, **^N^Strand 4** as compound **S4**, **^N^Strand 6** as compound **S6** and **^N^Strand 7** as compound **S7**, Fig. 1E). Notably, two of the selected sequences, **S4** and **S7**, exhibited the highest density of interacting sites and were spatially close, facing each other in an antiparallel orientation, albeit not directly connected. Consequently, the core section of these sequences (each consisting of 6 amino acids: ^151^TIVVVK^146^ and ^183^TKVILH^188^) was kept and incorporated into the piperidine-pyrrolidine β-turn inducer (compound **β-Hsp90**, Fig. 1F). Additionally, to further assess and underline the importance of the β-hairpin like structure of **β-Hsp90** for an optimal interaction with the target protein, the shorter **S4** and **S7** hexapeptides, inserted into the β-hairpin mimic **β-Hsp90**, were also synthesized (compounds **S4^short^** and **S7^short^**, Fig. 1F).

### Synthesis and conformational analysis

The preparation of the β-hairpin mimics **β-Tau** and **β-Hsp90** and of the peptides **H1**, **S4**, **S6**, **S7**, **S4^short^**, **S7^short^** was performed with satisfactory yields using classical Fmoc-based solid-phase peptide synthesis (SPPS) on the Rink amide resin (more detailed information and characterization of the compounds in supporting information). The preparation of the piperidine-pyrrolidine β-turn scaffold was done accordingly to our previous publications.^35–37^

Conformational analysis of compounds **β-Tau** and **β-Hsp90** was performed using Circular Dichroism spectroscopy (CD) in 20 mM phosphate buffer (pH 7.4) at 20°C and 37°C, showing for both compounds in aqueous medium a dynamic equilibrium between β-sheet conformation (negative Cotton effect at 218 nm) and random coil (negative band at 195 nm) structure as demonstrated by deconvolution^41^ (∼30% β-sheet/60% random coil, see SI, Fig. S1 and S2, and table S1). However, in MeOH, both compounds at 125 µM showed the typical bands of a β-sheet structure with **β-Tau** having a more pronounced tendency to aggregate and precipitate than **β-Hsp90** (see the decrease of CD signal intensity for **β-Tau** SI, Fig. S3 and Table S1 for deconvolution). These results are consistent with our previous studies using NMR, CD and molecular dynamics (MD) on this class of compounds based on the same piperidine-pyrrolidine β-turn inducer, that showed distinct degree of flexibility according to the peptide sequences.^35–37^ CD analysis of compound **H1** (^N^Helix 1) was also performed to evaluate its ability to maintain its helical structure outside the protein context. However, due to its limited solubility in aqueous medium, the CD analysis was only conducted in MeOH at 37°C. Remarkably, **H1** exhibited a typical helical structure characterized by two consecutive negative minima at 218 and 209 nm and a positive maximum at 195 nm (SI, Fig. S7). CD spectra for compounds **S4**, **S6**, **S7**, **S4^short^** and **S7^short^** were not registered, as their short peptide length was not expected to stabilize any significant secondary structure.

### Tau aggregation assays by Thioflavin-T (ThT) fluorescence spectroscopy

ThT is the standard dye used to study the aggregation kinetics of amyloidogenic proteins due to its ability to fluoresce upon binding to β-sheet rich structures.^42,43^ The aggregation curve of an amyloid protein alone classically displays a sigmoidal pattern, characterized by 3 different portions: i) an initial lag phase also called primary nucleation phase, characterized by the absence of a fluorescent signal due to the lack of β-sheet rich structures of the initial nuclei and small “on-pathway” oligomers; ii) a growth phase, also called elongation phase corresponding to the fast increment of the fluorescent signal caused by the formation of bigger β-sheet rich species (secondary nucleation); iii) a final plateau reached when the maximal level of mature fibrils is reached.^44^ The ability of the compounds to modulate the amyloid protein aggregation was assessed by ThT fluorescence spectroscopy, considering the time of the half-aggregation (*t_1/2_*), giving insight into the overall kinetics of aggregation and the intensity of the experimental fluorescence plateau (*F*), assumed to be influenced by the β-sheet content, morphology and quantity of aggregates formed.

#### β-Hairpin mimetics and ^N^Strand 7 of Hsp90 inhibit the aggregation of Ac-PHF6* hexapeptide model

Among the many short peptide models reported to mimic Tau aggregation without using Tau^FL^,^45–47^ the Ac-PHF6* hexapeptide model is one of the latest proposed, it is cheap and easy to synthesize and to handle, robust and able to aggregate rapidly.^48^ Thus, this model was used to screen the activity of all the prepared compounds (**β-Tau**, **β-Hsp90**, **H1**, **S4**, **S6**, **S7**, **S4^short^** and **S7^short^**) at 5/1, 1/1 and 0.1/1 compound/Ac-PHF6* ratios. Due to the high aggregative propensities of Ac-PHF6*, the lag phase period was not observed thus t_1/2_ could not be calculated (Extended Data Fig. 1), and only the relative maximum of fluorescence *F* with respect to the control curve was considered (Fig. 2A and Extended Data Fig.1 and Table 1).

**Fig. 2.**
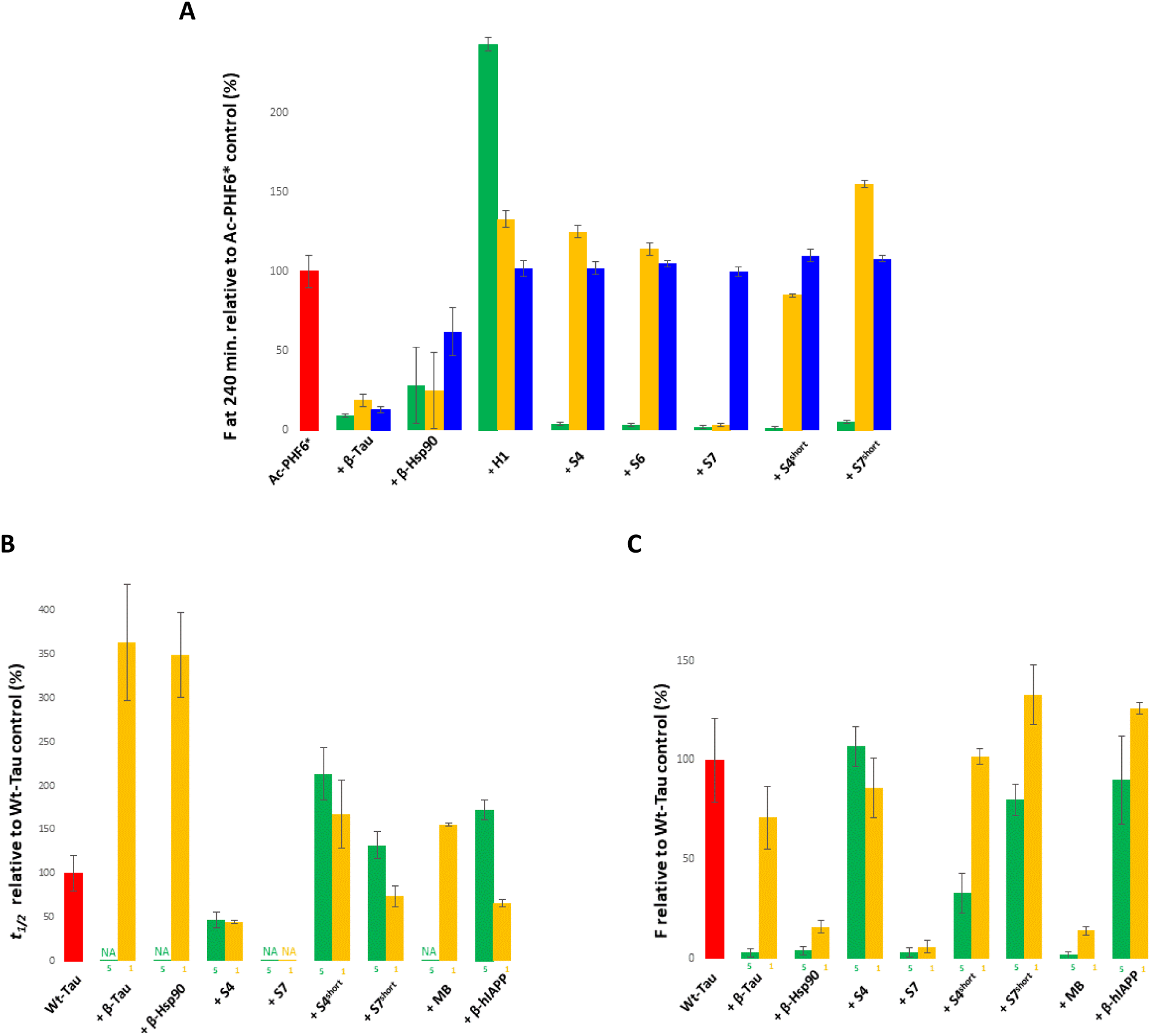
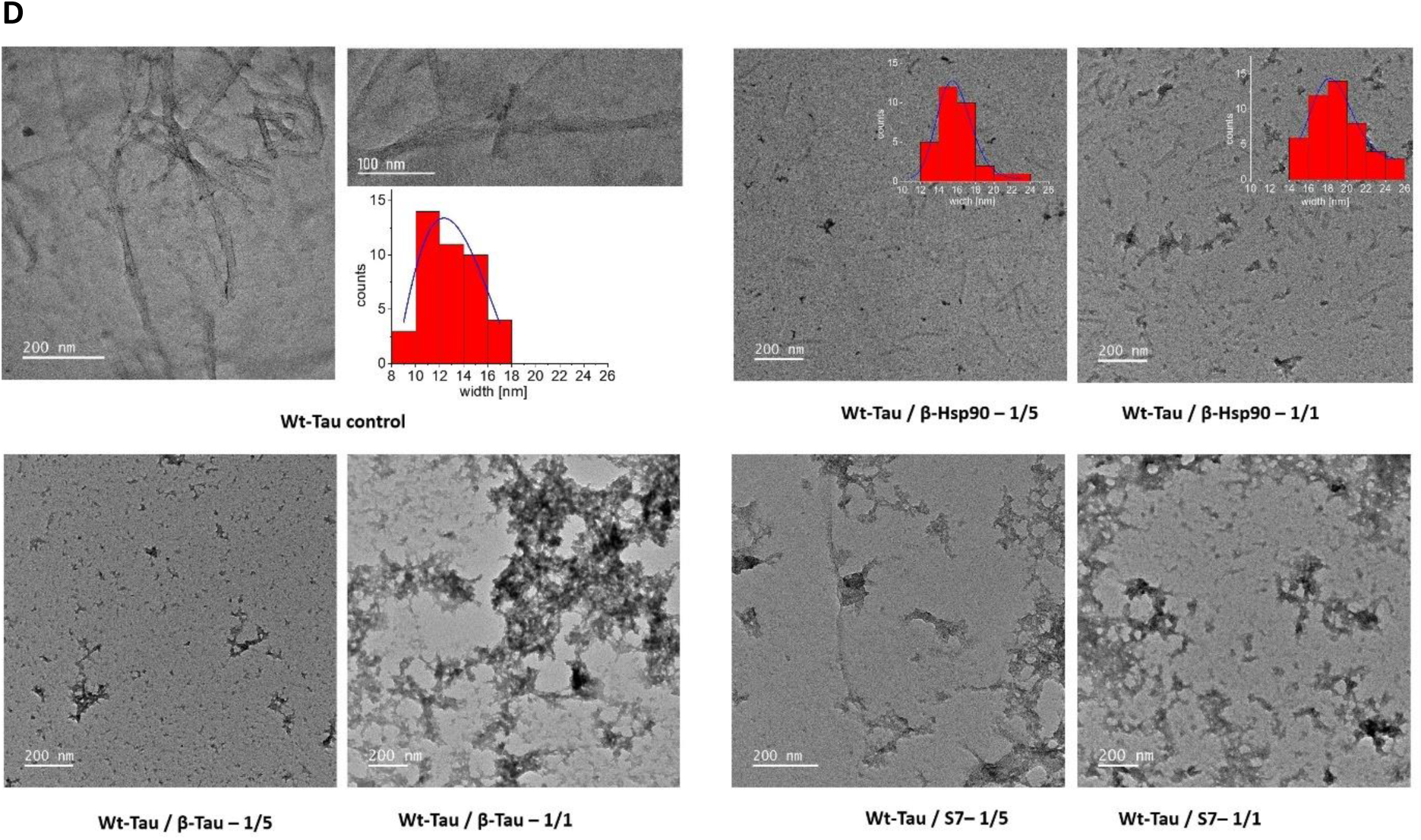
**A)**: ThT-fluorescence at 240 min of compounds **β-Tau**, **β-Hsp90**, **H1**, **S4**, **S6**, **S7**, **S4^short^** and **S7^short^** related to the maximum fluorescence obtained for the ThT-fluorescence control curve of Ac-PHF6* at 5/1 (green bars), 1/1 (yellow bars) and 0.1/1 (blue bars) compound/Ac-PHF6* ratios. Parameters are expressed as mean ± SE, n=3. Compounds were dissolved in water. The concentration of Ac-PHF6* was 25 µM. The fluorescence for Ac-PHF6* control was set at 100 % for the maximal fluorescence and the relative fluorescence was calculated related to this value. **B-C)** Effects of compounds **β-Tau**, **β-Hsp90**, **S4**, **S7**, **S4^short^**, **S7^short^**, **MB** and **β-hIAPP** on Wt-Tau:Hep fibrillization assessed by ThT-fluorescence spectroscopy at 5/1 (green bars) and 1/1 (yellow bars) ratios of compound/Wt-Tau. Parameters are expressed as mean ± SE, n=3. Compounds were dissolved in water. The concentration of Wt-Tau was 10 µM. **B)** The relative *t_1/2_* (%) was calculated related to the *t_1/2_* of Wt-Tau control (see Methods for the detailed calculations). NA = No Aggregation over 30h and **C)** The relative *F* (%) was calculated at 30 h related to the fluorescence plateau of Wt-Tau control. **D)** Transmission electron micrographs of Wt-Tau with inserted width distribution of the observed ordered structures. Fibrils of Wt-Tau:Hep (control) at low and high magnification; rod-like structures of Wt-Tau:Hep in the presence of **β-Hsp90** (5/1 and 1/1 ratios); amorphous aggregates of Wt Tau in the presence of **β-Tau** (5/1 and 1/1 ratios) and **S7** (5/1 and 1/1 ratios).

Compound **H1** (^N^Helix 1) alone showed strong ThT fluorescence at 125 µM suggesting a self-aggregating behaviour, which can be due to its hydrophobic character and its low solubility in aqueous medium (Extended Data Fig. 1). At 5/1 and 1/1 ratios (**H1**/Ac-PHF6*), the increase of *F* (+143% and +33% respectively) suggested a co-aggregation process of **H1** with Ac-PHF6* or rather simply the addition of the two individual *F* values. On the contrary, the two β-hairpin mimics **β-Tau** and **β-Hsp90** along with the Hsp90-derived peptides **S4**, **S6**, **S7**, **S4^short^** and **S7^short^**, displayed excellent activities at 5/1 ratio, almost completely suppressing the fluorescence. At stoichiometric ratio only the hairpin mimics **β-Tau**, **β-Hsp90** and peptide **S7** maintained excellent activities (respectively, F = -81%; -75%; -97%) while **S4**, **S6** and **S7^short^** displayed a pro-aggregative effect with an overall increment in the detected fluorescent level up to +155%. Remarkably, at sub-stoichiometric ratio (0.1/1), only the β-hairpin mimics **β-Tau** and **β-Hsp90** exhibited activity, with **β-Tau** in particular giving comparable results to those observed at higher ratios (F = -87%). Interestingly, the simultaneous presence of both **S4^short^** and **S7^short^** at 2.5 + 2.5/1 ratio let to maintain a strong activity which is reduced at 0.5 +0.5 /1 ratio while no major effect was observed in 0.05 + 0.05/1 ratio (Extended Data Fig. 1). These preliminary results on Ac-PHF6* hexapeptide model underly the relevance of the hairpin secondary structure and of the two rational design strategies based on Tau or Hsp90 sequences.

#### β-Hairpin mimetics inhibit the aggregation of the full-length Wt-Tau in vitro

The addition of anionic co-factors like heparin is mandatory to induce the aggregation of Tau^FL^. However, heparin-induced Tau^FL^ aggregates are different from those observed in diseases.^49,50^ Thus, Tau aggregation assay was performed in 25 mM phosphate buffer at pH 6.6, as reported by Huvent *et al.*, ^51^ but with a decreased Tau/heparin ratio (160/1), to induce the Tau assembly in an almost co-factor-free environment.^52^ The ThT-fluorescence curves were nicely reproducible and characterized by a sigmoidal shape with a lag phase of around 5 hours followed by an elongation phase and a final plateau reached after 12 hours. The two parameters *t_1/2_* and *F* were used to evaluate the activity of compounds **β-Tau**, **β-Hsp90**, **S4**, **S6**, **S7**, **S4^short^** and **S7^short^**, which were compared to the activity of Methylene Blue (**MB**, methylthioninium chloride, considered as a reference Tau inhibitor, analogue of the reduced form hydromethylthionine mesylate, LMTM or TRx0237, which has been in phase 3 clinical trials conducted by TauRx Therapeutics Ltd for mild to moderate AD (Fig. 2B and 2C and Extended Data Fig. 2 and Table 2).

Only β-hairpin mimics **β-Tau**, **β-Hsp90**, peptide **S7,** and **MB** showed a complete inhibition of the aggregation of Wt-Tau at 5/1 ratio. In contrast with **MB**, β-hairpin mimics **β-Tau** and **β-Hsp90** and peptide **S7** were still very active at 1/1 ratio to delay the aggregation (*t_1/2_* = +55% for **MB**; +266% for **β-Tau**; + 249% for **β-Hsp90** and N.A. for **S7**) and reduce *F* (*F*=-86%; -29%; -84%; -94% respectively), with **β-Hsp90** and peptide **S7** being the most active compounds on both parameters (Fig. 2B and 2C and Extended Data Fig. 2 and Table 2 ). At the very low ratio 0.1/1, **β-Tau**, **β-Hsp90**, peptide **S7** and **MB** showed no significant inhibitory effect (consistently with literature data,^53^ **MB** did not delay the kinetics but decreased *F* by 50%). The rationale of the choice of the peptide sequences was also challenged by the evaluation of a similar β-hairpin mimic (**β-hIAPP)**, recently reported by some of us,^37^ based on human islet amyloid polypeptide (hIAPP, also known as amylin) sequences (A_13_N_14_F_15_L_16_V_17_ and N_22_F_23_G_24_A_25_I_26_L_27_) and whose aggregation is involved in type 2 Diabetes (T2D). As expected, **β-hIAPP** was much less active with no activity on *F* at any ratio, only a small ability to delay the aggregation kinetics at 5/1 ratio and even slightly speeding up the aggregation process at 1/1 and 0.1/1 ratios (Fig. 2B and 2C and Extended Data Fig. 2 and Table 2).

The superior inhibitory effect of **β-Hsp90** among our lead compounds was next confirmed by transmission electron microscopy (TEM) experiments of Wt-Tau at the end of the aggregation process (96 h). At the low ratio of heparin used for the ThT assays (Tau/heparin ratio 160/1), control Wt-Tau formed long amyloid fibrils with an average diameter of 12.6 ± 0.1 nm with various morphologies as previously reported (Fig. 2D).^49^ In the presence of our lead compounds **β-Tau**, **β-Hsp90**, and **S7** at 5/1 and 1/1 ratio, the formation of the characteristic Wt-Tau fibrils was inhibited (Fig. 2D). Remarkably, in the presence of **β-Hsp90**, the mixture Tau:Hep formed short rod-like nanostructures with an average diameter of 15.7± 0.2 nm at a ratio of 5/1 and 18.5± 0.1 nm at a ratio of 1/1 (Fig. 2D) which were ThT negative (Fig. 2D). In contrast, **β-Tau** and **S7** formed amorphous aggregates. In the case of **β-Tau** at a ratio of 1/1, large amorphous aggregates were observed, probably with a high β-sheet content that gave a positive ThT signal. These aggregates were not observed at a ratio of 5/1. (Fig. 2D). The efficiency of the hairpin mimic based on the chaperone Hsp90 (**β-Hsp90**) at a higher heparin ratio (Tau/heparin 4/1) was finally investigated. As expected, Tau: Hep formed fibrils; meanwhile, in the presence of **β-Hsp90**, only amorphous structures and not detectable by ThT were observed (Extended Data Fig. 3). Altogether, all the tested compounds inhibited the formation of typical amyloid Tau fibrils. Interestingly, **β-Hsp90** promoted the formation of regular rod-like structures. Meanwhile, the others favor the amorphous pathway, although **β-Tau** at 1/1 probably had a high β-sheet content because of the positive ThT signal obtained in the initial screening.

### Stability toward proteolysis

Before the cell evaluation of the most active compounds **β-Tau**, **β-Hsp90** and **S7**, their stability towards proteolysis was assessed. The aim was to observe whether the incorporation of the unnatural piperidine-pyrrolidine β-turn inducer could limit enzymatic degradation by comparing the stability of **β-Hsp90** with **S4^short^**, **S7^short^**. Stability studies are a fundamental phase of drug development, especially with peptide-based compounds, which are easily degraded by proteolytic enzymes ubiquitously found inside and outside cells. In the Pronase medium, which has a broad specificity (mixture of neutral protease, chymotrypsin, trypsin, carboxypeptidase, and aminopeptidase), natural peptide **S7** and **S7^short^** were highly vulnerable to proteolysis, with **S7** completely degraded and only 46% of **S7^short^** remaining after 30 minutes (Extended data Table 3). Conversely, the shorter peptides **S4^short^** and **β-Tau** displayed more pronounced stability with 71% and 30% of peptides intact after 2 hours respectively. Remarkably, **β-Hsp90** emerged as the most stable candidate, showing a remarkable resistance to proteolytic degradation with complete stability up to 1 hour and 79% of the compound intact after 2 hours (detailed cleavages are represented in Extended data Fig. 4). Overall, the incorporation of the non-natural β-turn inducer notably enhanced the stability of **β-Hsp90** and limited the degradation of **β-Tau**. Thus, *in vitro* aggregation and stability studies allowed us to identify the two hairpin structures, **β-Tau** and **β-Hsp90**, as the most promising candidates for further development, while peptide **S7** exhibiting extreme instability towards proteolytic degradation was discarded for cell evaluation.

#### β-hairpin mimetics also inhibit aggregation of the pro-aggregative Tau-ΔK280 mutant in vitro

As the assessment of the compounds to inhibit Tau aggregation and to restore physiological Tau-MT interaction in cells implies model neurons expressing Tau-ΔK280 (single amino acid deletion mutant, which has been reported in two cases of tauopathies^54–56)^, an *in vitro* ThT assay on Tau-ΔK280 was carried out for **β-Tau** and **β-Hsp90**.

The ThT-fluorescence assays revealed that **β-Hsp90** was notably more efficient than **β-Tau** (this superiority being much more pronounced than towards Wt-Tau, Fig. 3A). In the presence of **β-Hsp90**, a complete inhibition of the aggregation of Tau-ΔK280 at both 5/1 and 1/1 ratios was observed. An interesting activity in decreasing *F* at the very low 0.1/1 ratio was preserved (F= -66%, Fig. 3A, 3B and 3C, and Extended Data Table 4 and Fig. 5). **β-Tau** suppressed the aggregation at 5/1 ratio, kept an interesting but lower activity at 1/1 ratio (*t_1/2_*= +179%, F= -75%), and did not show any activity at 0.1/1 ratio (Fig. 3A, 3B and Extended Data Table 4 and Fig. 5).

**Fig. 3:**
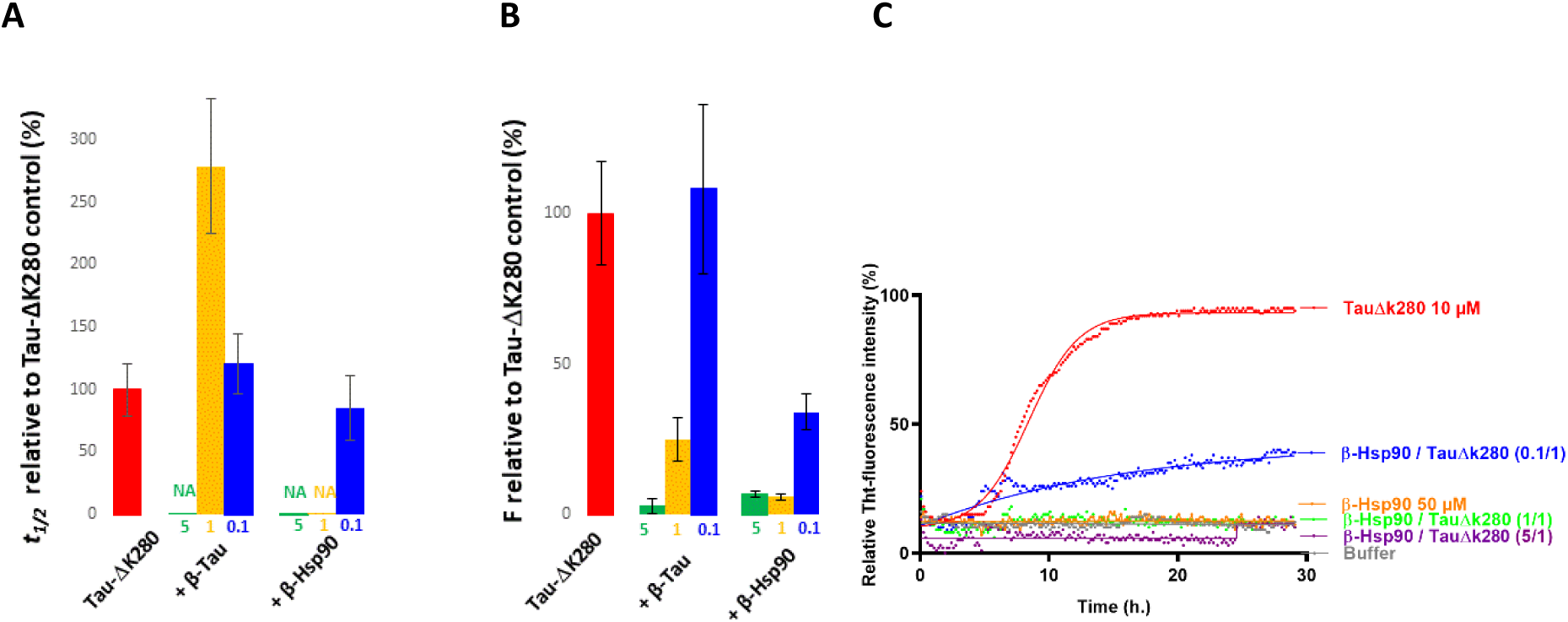
**A-B)** Effects of compounds **β-Tau** and **β-Hsp90** on Tau-ΔK280 fibrillization assessed by ThT-fluorescence spectroscopy at 5/1 (green bars) 1/1 (yellow bars) and 0.1/1 (blue bars) ratios of compound/Tau-ΔK280. Parameters are expressed as mean ± SE, n=3. Compounds were dissolved in water. The concentration of TauΔK280 was 10 µM. A) The relative *t_1/2_* (%) was calculated with respect to the *t_1/2_* of Tau-ΔK280 control (see Methods for the detailed calculations). NA = No Aggregation over 30 hours. **B)** The relative *F* (%) was calculated at 30 h related to the fluorescence plateau of Wt-Tau control. **C)** Representative curves of ThT fluorescence assays over time showing Tau-ΔK280 fibrillization (10 µM) in the absence (red curve) and in the presence of **β-Hsp90** at compound/Tau-ΔK280 ratios of 5/1 (purple curves), 1/1 (green curves) and 0.1/1 (blue curves). The control curve of the compound is represented in orange line and the buffer in grey.

**Fig. 4.**
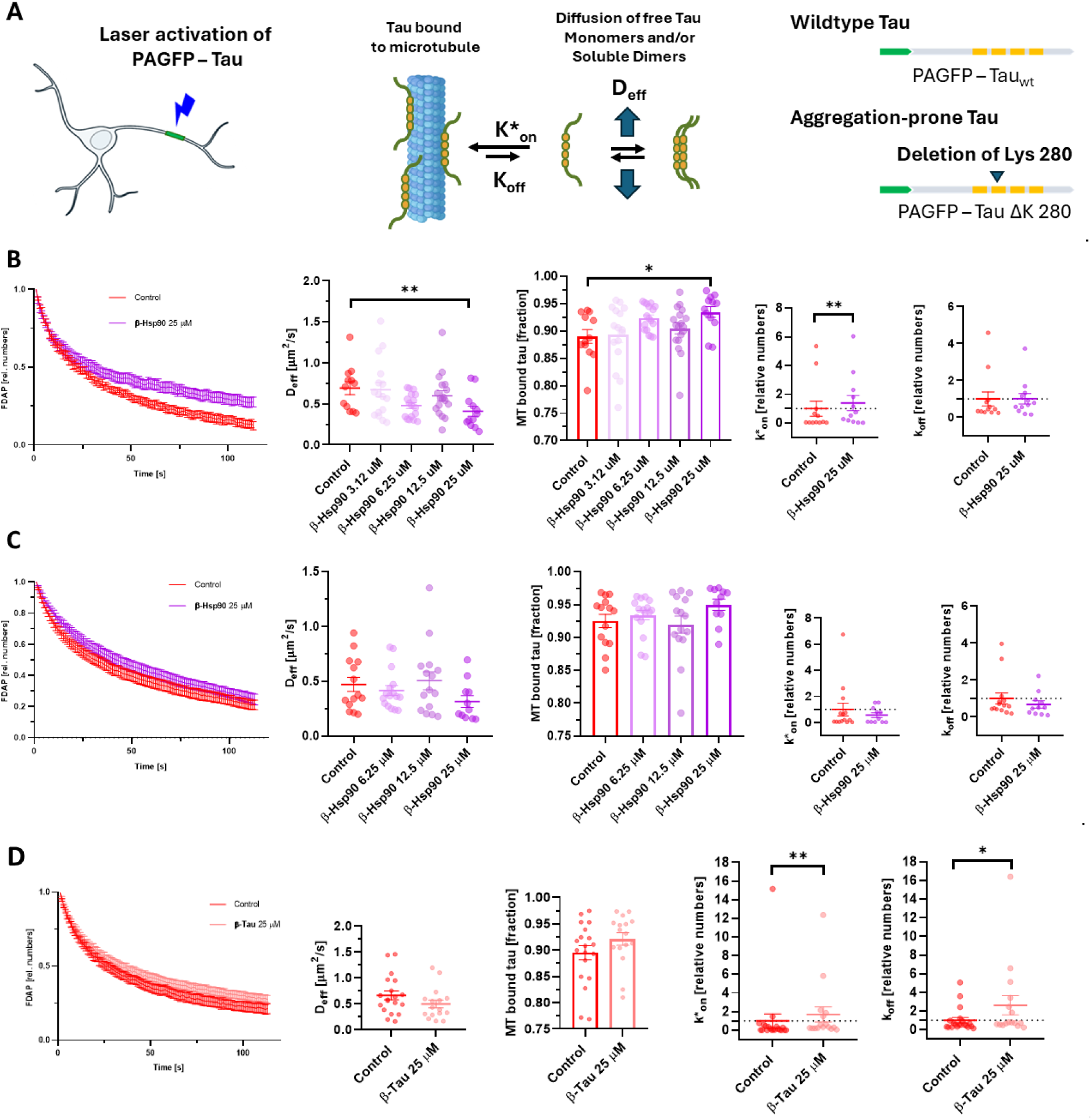
Quantitative live cell imaging. **A)** Schematic representation of the live cell imaging approach to determine Tau dynamics in axon-like processes of neuronally differentiated PC12 cells. The schematic shows that calculation of the effective diffusion constant (D_eff_) using a one-dimensional diffusion model allows determination of the fraction of microtubule-bound Tau).^59^ Schematic representations of the expressed Tau constructs are shown on the right. The MT-binding repeat regions (RR1–RR4) are indicated by yellow boxes, the N-terminal PAGFP fusion by green. The lysine deletion of the aggregation-prone construct is located in the second repeat. **B)** Effect of **β-Hsp90** on the dynamics of aggregation-prone Tau in axon-like processes. FDAP diagrams, scatterplots of effective diffusion constants (D_eff_), bar graphs of the fraction of Tau bound to microtubules, and scatterplots of k_on_ and k_off_ rate constants are shown. Mean ± SEM of 12 (control), 16 (3.12 µM β-Hsp90), 17 (6.25 µM β-Hsp90), 18 (12.5 µM β-Hsp90) and 12 (25 µM β-Hsp90) cells are shown. **C)** Effect of β-Hsp90 on wildtype Tau dynamics in axon-like processes. In contrast to aggregation-prone Tau, β-Hsp90 does not affect fluorescence decay, D_eff_, MT-bound fraction and k_on_ and k_off_ values. This indicates that increased binding of aggregation-prone Tau is caused by inhibition of the formation of soluble Tau aggregates without affecting physiological tau-microtubule interaction. Shown are mean ± SEM of n = 14 (control), n = 16 (6.25 µM β-Hsp90), n = 15 (12.5 µM β-Hsp90), n = 11 (25 µM β-Hsp90) cells. **D)** Effect of β-Tau on the dynamics of aggregation-prone tau in axon-like processes. However, β-Tau leads to a significant increase in the k_on_ and k_off_ rates, suggesting that it increases the dynamics of the Tau-microtubule interaction. Mean ± SEM of 18 (control) and 16 (25 µM β-Tau) cells are shown. Statistically significant differences from control determined by one-way ANOVA with Dunnett’s post hoc test (B, C) or unpaired two-tailed Student’s t-tests (D) are indicated. *p <0.05, ** p<0.01. In all experiments, incubation with the compound lasted 24 hrs and the final concentration of DMSO was 0.125%.

**Fig. 5:**
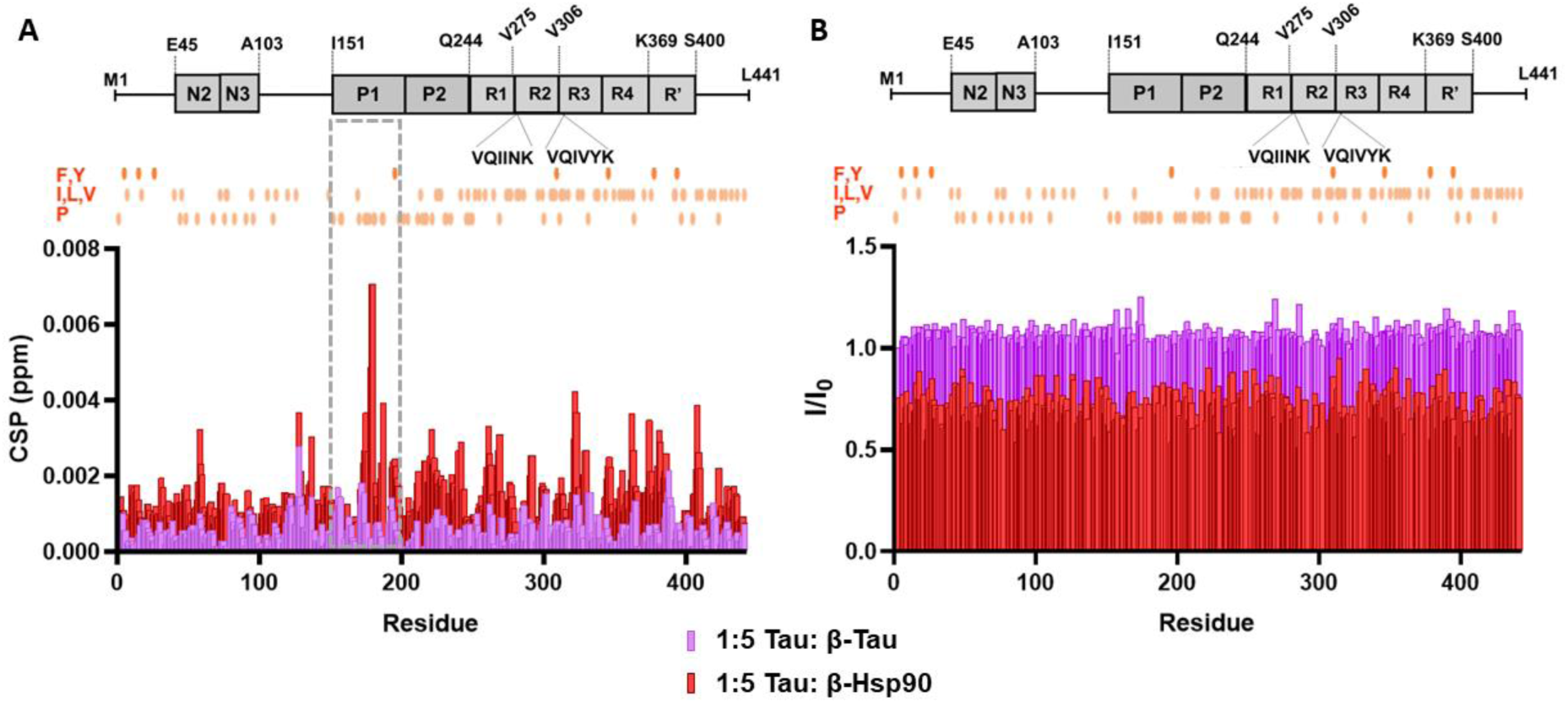
Titrations of ^15^N HSQC of 2N4R ^15^N-Tau (20 μM) with **β-Tau** and **β-Hsp90** in buffer solution (50 mM NaPi pH 6.8, 10 mM NaCl, 1 mM DTT, 10% D_2_O) at 5°C (900 MHz or 800 MHz) at a mole ratio of Wt-Tau_441_/**compounds** = 1/5. Indicated in orange circles (**A** and **B**) are the locations of selected hydrophobic amino acids (F,Y,I,L,V,P) in the sequence of Wt-Tau. **A)** Chemical shift perturbations (CSP) of Tau residues in presence of **β-Hsp90** (red) are more prominent than those of **β-Tau** (violet) particularly in the P1 region of Tau (residues ∼175-187, within the region boxed in grey). **B)** I/I_0_ values of Tau residues (I represents NMR signal intensity in the presence of peptide, and I_0_ corresponds to conditions in the absence of compounds) showed a ∼20% decrease upon addition of **β-Hsp90** at the 1/5 mole ratio, however this was not observed for the same mole ratio upon addition of **β-Tau**.

#### Quantitative live cell imaging shows that β-hairpin based on Hsp90 restores the physiological microtubule interaction of aggregation-prone Tau in model neurons

β-Hairpin mimetics **β-Tau** and **β-Hsp90** were then evaluated for their ability to inhibit Tau aggregation and to restore physiological Tau-MT interaction in axon-like processes of model neurons expressing Wt-Tau and aggregation-prone Tau mutant (Tau-ΔK280) (Fig. 4A). Assessment of neuronal cell toxicity using MTT and LDH assays^57,58^ on the model neurons revealed no cell membrane damage or metabolic impairment at concentrations up to 100 μM for **β-Tau** and **β-Hsp90** (Extended Data Fig. 6)

The quantitative live-cell imaging approach is based on the photoactivation of a defined population of Tau molecules, followed by fluorescence decay after photoactivation (FDAP) measurements to determine in the dynamics of the interaction of Tau with MTs in axon-like processes (Fig. 4A).^59,60^ After focal photoactivation, Tau-ΔK280 showed dissipation from the region of photoactivation, which was significantly reduced in the presence of **β-Hsp90** (Fig. 4B, left). Accordingly, calculating the effective diffusion constant (Deff) from the FDAP curves using a one-dimensional diffusion model resulted in lower values for **β-Hsp90** -treated cells, which became significant at 25 µM treatment (Fig. 4B, middle). This corresponds to an approximately 5% increased binding of Tau-ΔK280 to the microtubules. The application of a previously developed refined reaction-diffusion model made it possible to estimate the binding constants (k*_on_ and k_off_ rates) of the Tau-MT interaction.^59,60^ While k_off_ rates did not differ, treatment with **β-Hsp90** resulted in an increased k*_on_ rate, indicating that disruption of Tau-ΔK280 aggregation increases the availability of Tau-ΔK280 for binding to microtubules.

To confirm that **β-Hsp90** acts by reducing Tau aggregation and and does not directly affect the Tau-MT interaction, we also performed similar FDAP assays with Wt-Tau expressing cells. As expected, Wt-Tau showed increased interaction with axonal microtubules compared to Tau-ΔK280 due to the lack of oligomerization under the cellular conditions. Interestingly, in the presence of Wt-Tau, **β-Hsp90** did not induce any change in any of the parameters indicating that **β-Hsp90** does not affect the physiological interaction of Tau with MTs (Fig. 4C). Remarkably, the fraction of microtubule-bound Wt-Tau in the control sample (no compound) was very similar to the fraction of the aggregation-prone Tau-ΔK280 in the presence of 25 µM of **β-Hsp90**. This observation demonstrates that the β-hairpin mimetic based on the chaperone Hsp90 effectively inhibits Tau-ΔK280 aggregation in cells and thus fully restores the Tau-MT interaction back to physiological levels. This also suggests that the interaction of **β-Hsp90** with Tau does not prevent the binding of Tau to MTs.

Conversely, **β-Tau** exhibited a smaller impact on most parameters (in particular, Deff and MT bound Tau) suggesting that **β-Tau** has a much lower activity to reduce Tau aggregation than **β-Hsp90** in model neurons (Fig. 4D). Remarkably, treatment with **β-Tau** still resulted in a significantly increased k*_on_ indicating that the presence of **β-Tau** still increases the availability of aggregation-prone Tau-ΔK280 to axonal MTs. Interestingly, **β-Tau** also increased the k_off_ rate of the Tau-MT interaction indicating that it increases the dynamics of the Tau-MT interaction. This may indicate that, by mimicking Tau hot spots (see above the design), **β-Tau** competes with Tau to interact with MTs.

### Titration of ^15^N-Tau with β-hairpin mimics β-Tau and β-Hsp90

In order to obtain molecular information about the interaction between Tau and our β-hairpin mimics, 2D ^1^H-^15^N HSQC titrations of ^15^N-Wt-Tau (20 μM) were conducted in the presence of **β-Hsp90** or **β-Tau** at different concentrations (20 μM, 40 μM, and 100 μM). Conversely to **β-Tau**, in the presence of **β-Hsp90**, a global signal broadening of Tau was observed, suggesting that **β-Hsp90** induces a slight oligomerization of Tau. Chemical shift perturbations (CSPs) induced by **β-Hsp90** were significantly larger than those observed in the presence of **β-Tau**, suggesting that Tau has a higher affinity for **β-Hsp90**. Remarkably, in all molar ratios tested, the largest CSPs observed in the presence of β-Hsp90 were predominantly confined to the P1 region of Tau (residues ∼175-187), as depicted in Fig. 5 (due to the large extent of broadening observed in titrations with Tau and **β-Hsp90**, it was not possible to determine the binding constant (K_D_) from CSP values). The observation that **β-Hsp90** does not bind to the Tau MTBR surely explains the effectiveness of **β-Hsp90** in inhibiting Tau aggregation without preventing Tau-MTs interaction. On the other hand, the absence of CSP and signal broadening in the presence of **β-Tau** suggests that this compound does not significantly interact with Tau protein or that this interaction is too transient at the NMR level (Extended data Fig. 7). This might explain the lower efficiency of **β-Tau** compared to **β-Hsp90** in decreasing Tau toxic aggregation both *in vitro* and in cells.

### β-Hairpin mimetic β-Hsp90 inhibits the aggregation of Aβ_1-42_ *in vitro* and act as a dual and selective inhibitor for AD

As Tau and Aβ_1-42_ mutually interact during AD pathology causing a progressive worsening of symptoms,^61^ and as Hsp90 was reported to inhibit Aβ_1-42_ aggregation even at a 50-fold lower ratio,^32,33^ the potential effect of our β-hairpins on the aggregation process of Aβ_1-42_ was investigated. We followed our classical aggregation protocol,^35,36^ and the control curves were characterized by a lag phase of around 1-5 hours, followed by a growth phase that reached the plateau after 10-15 hours. β-hairpin mimics **β-Tau** and **β-Hsp90** showed an effect on Aβ_1-42_ aggregation at a 10/1 (compound/Aβ_1-42_) ratio, being able to completely suppress the formation of the fibril (Fig. 6A and 6B, Extended Data Table 5 and Fig. 8). At the lower 1/1 ratio, only compound **β-Hsp90** based kept a strong inhibitory activity (t_1/2_= +208%, F= -90%), while **β-Tau** induced a modest delay on the aggregation kinetics (t_1/2_= +27%) and no effect on *F*. **β-Hsp90** even kept a small activity at 0.1/1 ratio (t_1/2_= +94%, F= -30%). Thus, we next investigated the activity of the isolated β-strand sequences of Hsp90 **S4** and **S7**, and of the hexapeptides **S4^short^** and **S7^short^** present in **β-Hsp90**. Conversely to what was observed for Tau, only peptides **S4** and **S4^short^** slightly delayed the aggregation at the high 10/1 ratio, while **S7** and **S7^short^** were completely inefficient in decreasing Aβ_1-42_ aggregation.

**Fig. 6:**
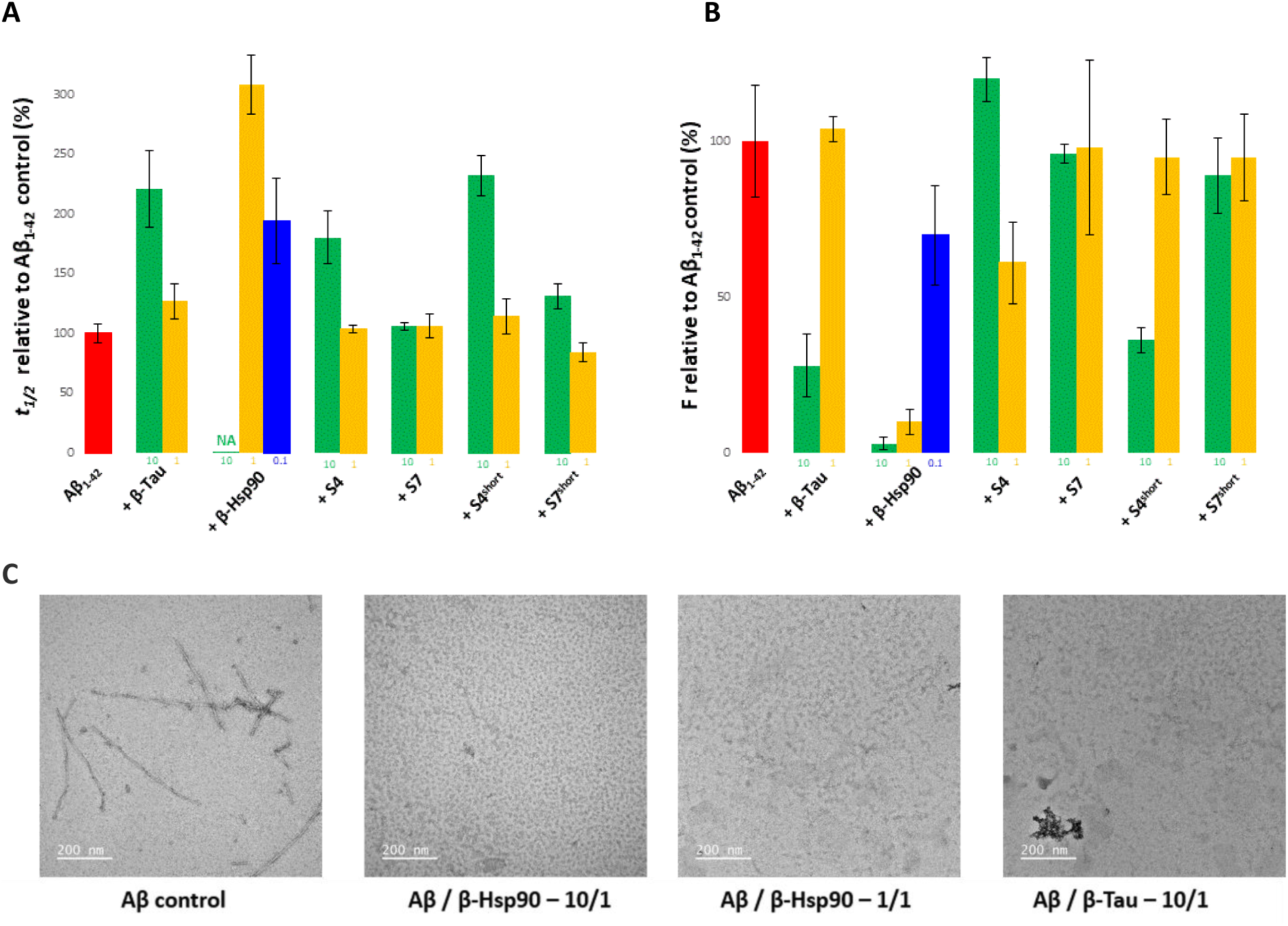
**A-B)** Effects of compounds **β-Tau**, **β-Hsp90**, **S4**, **S7**, **S4^short^** and **S7^short^** on Aβ_1-42_ fibrillization assessed by ThT-fluorescence spectroscopy at 10/1 (green bars) 1/1 (yellow bars) and 0.1/1 (blue bars, only for **β-Hsp90**) ratios of compound/Aβ_1-42_. Parameters are expressed as mean ± SE, n=2. Compounds were dissolved in water. The concentration of Aβ_1-42_ was 10 µM. **A**) The relative *t*_1/2_ (%) was calculated with respect to the *t*_1/2_ of Aβ_1-42_ control, see supporting information for the detailed calculations. NA = No Aggregation over 42 hours and **B)** The relative F (%) was calculated at 42 h related to the fluorescence plateau of Aβ_1-42_ control. **C)** Transmission electron micrographs, Aβ_1-42_ control and in the presence of **β-Hsp90** (10/1 and 1/1 ratios) and **β-Tau** (10/1 ratio).

Notably, while **β-Hsp90** and, to a lesser extent, **β-Tau** are promising dual aggregation inhibitors of the two proteins involved in AD, they did not show interesting activity on hIAPP fibrillization kinetics, an amyloid protein involved in T2D. In our ThT assays, both **β-Hsp90** and **β-Tau** even increased the fluorescence plateau at 10/1 (compound/hIAPP) ratio (with no significant activity at 1/1 ratio, Extended data Fig. 9).

The inhibitory results obtained with our β-hairpin mimics on Aβ_1-42_ aggregation were confirmed by TEM, demonstrating the absence of Aβ_1-42_ fibrils at the end of the aggregation process (24 h). In the case of **β-Hsp90** and **β-Tau** at this 10/1 ratio, non-uniform structures were detected (Fig. 6C).

2D ^1^H-^15^N HSQC titrations of ^15^N-Aβ_1-42_ (50 μM) in the presence of **β-Hsp90** (50 μM and 500 μM) were then performed. No CSP or broadening of Aβ_1-42_ ^1^H-^15^N correlations could be observed at **β-Hsp90**/Aβ_1-42_ molar ratios of 1:1 or 10:1. A 50% decrease of all Aβ_1-42_ signals was nevertheless observed in the presence of a 10 folds excess of **β-Hsp90**, suggesting a co-aggregation process (Extended data Fig. 10). The absence of CSP or signal broadening in the presence of an equimolar ratio of **β-Hsp90** suggests that this compound does not significantly interact with Aβ_1-42_ monomers or that the interaction is too transient at the NMR scale. Therefore, this compound rather interacts with larger oligomers that are not visible by NMR.

## Discussion

In this work, the possibility of inhibiting toxic Tau aggregation was investigated by designing new β-hairpin mimics **β-Tau** and **β-Hsp90** containing a piperidine-pyrrolidine β-turn inducer bearing peptide sequences based on Tau SREs and inspired by the physiological interaction of Tau with MTs interactions (PHF6 and PHF6*) for **β-Tau**, and on two peptide sequences (**^N^Strand 4** and **^N^Strand 7**) of the physiological chaperone Hsp90 for **β-Hsp90**. Four different peptides extracted from the crystal structure of Hsp90 *N*-terminal domain (PDB=3NMQ) were synthesized and found to be the most relevant interacting regions and possible binding sites of ^N^Hsp90 in cross-interaction with Tau (**^N^Helix 1** as compound **H1**, **^N^Strand 4** as compound **S4**, **^N^Strand 6** as compound **S6** and **^N^Strand 7** as compound **S7**). Their synthesis on solid phase was efficient. The conformational analysis of **β-Tau** and **β-Hsp90** by CD confirmed the presence of a flexible β-hairpin-like structure in aqueous medium (PBS) while a stabilized β-hairpin was observed in the organic protic solvent MeOH.

**β-Tau** and **β-Hsp90** showed excellent ability to inhibit Wt-Tau aggregation *in vitro* as demonstrated by ThT assays and TEM imaging, with superiority of the synthetic chaperone **β-Hsp90**. Among the four peptides **H1**, **S4**, **S6** and **S7**, also evaluated by ThT, only **S7** demonstrated activity against Wt-Tau aggregation. The need to use pre-organized β-hairpin structures to ensure inhibitory activity on Tau aggregation and ensure stability against proteolysis was demonstrated. **β-Hsp90** clearly proved to be the most stable candidate against proteolysis, showing a complete stability up to 1 hour and only 21% degradation after 2 hours. **β-Tau** showed a good stability up to 1 hour (62% of intact **β-Tau**) but was more degraded after 2 hours (64% of degradation). Despite the interesting *in vitro* activity of **S7**, its high sensitivity to proteolysis (76% and 100% degradation, after 10 and 30 minutes respectively) led us to discard it from further cellular studies and to continue with our two β-hairpin mimics **β-Tau** and **β-Hsp90**. The superiority of **β-Hsp90** was confirmed on the aggregation-prone Tau mutant ΔK280-Tau, which single mutation has been reported in two cases of tauopathies.^54–56^ In the ThT assays conducted in the presence of **β-Hsp90**, a complete inhibition of the aggregation of Tau-ΔK280 at both 5/1 and 1/1 ratios was observed, and an interesting activity at the very low 0.1/1 ratio was preserved. Conversely, **β-Tau** suppressed the aggregation only at the 5/1 ratio, but it showed a lower activity at 1/1 ratio and no activity at 0.1/1 ratio.

This difference between **β-Hsp90** and **β-Tau** was even more remarkable in the quantitative live cell imaging on model neurons where only **β-Hsp90** effectively inhibits toxic Tau-ΔK280 aggregation and fully restores the Tau-MT interaction up to physiological levels (same levels as with Wt-Tau). This highly interesting result also suggests that the interaction of **β-Hsp90** with Tau does not prevent the binding of Tau to MTs. Indeed, our NMR studies showed that **β-Hsp90** was predominantly interacting with the P1 region of Tau (in particular, residues ∼175-187), rather than with the region involved in the binding of Tau to MTs (MTBR). The reported observations in literature that Hsp90 alone could interact with Tau MTBR to increase oligomerization,^30,31^ or in multi-complex systems with the proline-rich region P1 (in Hsp90/FKBP51/Tau complexe^28^), or P2 and R’ flanking Tau-RD (in Hsp70/Hsp90 complex^29^), to prevent Tau aggregation, might indicate that our small synthetic chaperone **β-Hsp90** is capable of acting as multi-protein systems. Furthermore, the P1 region of Tau targeted by **β-Hsp90** includes two major phosphorylation site, Thr175 and Thr181 (in particular, the concentration of Tau phosphorylated at site 181 is a well-established biomarkers of AD in the cerebrospinal fluid (CSF)^62^ and a studied plasma biomarker^63^). However, the slight oligomerization observed in our NMR and TEM analyses of Tau in the presence of **β-Hsp90** is in accordance with some literature data reporting that Hsp90 inhibits the formation of Tau fibrils but could promotes the formation of small Tau oligomers.^26,27^ Furthermore, while the question of whether the interaction Hsp90-Tau allows to preserve the physiological interaction between Tau MTBR and MT has not yet been addressed, our cellular FDAP studies indicate that **β-Hsp90** might induce non-toxic small off-pathway oligomers that can prevent deleterious interactions of Tau aggregates with cytoplasmic proteins (such as kinases) or that can be in equilibrium with monomers able to interact with MTs.

Remarkably, **β-Tau** based on PHF6 and PHF6* was also able to inhibit *in vitro* Wt-Tau and Tau-ΔK280 aggregation, albeit more weakly than **β-Hsp90**, and with reduced inhibitory activity in cells. This smaller activity could be due to its lower proteolysis stability or to its interaction with Tau MTBR that decreases the subsequent Tau-MT interaction. However, its lack of visible interaction with Tau in the NMR studies rather suggests that by mimicking Tau-MT hot spots, **β-Tau** might compete with Tau and interact with MTs, which is consistent with the increased k_off_ observed in the FDAP assay in the presence of **β-Tau**. Additional highly interesting results obtained in this work concern the selectivity of our designed compounds for AD. Indeed, a similar β-hairpin mimic (**β-hIAPP)**, recently reported by us,^37^ based on hIAPP sequences inhibiting hIAPP aggregation involved in T2D was inactive towards Tau aggregation. Conversely, **β-Hsp90** and **β-Tau** did not inhibit hIAPP aggregation and even significantly increased the aggregation in the ThT assays, at the high ratio 10/1. A link between Tau and hIAPP was highlighted by the demonstration that these two proteins act in a coordinated manner to impair beta cell function and glucose homeostasis, linking AD and T2D.^64^ The increased aggregation induced by **β-Tau** is in accordance with this report and indicates that PHF6 and PHF6* might be part of the hot-spots of Tau-hIAPP cross-interaction. Even if the ubiquitin–proteasomal system, which includes Hsp90, is important for hIAPP clearance,^65^ to our knowledge, a direct link between Hsp90 and hIAPP has not been yet evidenced. Our results might suggest a potential but not beneficial Hsp90-hIAPP interaction.

Importantly, **β-Hsp90** was also found to be highly efficient to inhibit the aggregation of Aβ_1-42,_ which is the second amyloid protein involved in AD, thus confirming the preliminary data found in the literature reporting the inhibition of Aβ_1-42_ aggregation by the physiological Hsp90 chaperone.^32,33^ Our results with the isolated peptides indicate that, while **^N^Strand 7** of Hsp90 (peptide **S7**) seems to be the key sequence of Hsp90 interacting with Tau, **^N^Strand 4** (peptide **S4**) appeared to rather be important for inhibiting Aβ_1-42_ aggregation. In contrast, **β-Tau** was much less efficient in inhibiting Aβ_1-42_ aggregation at low ratios **β-Tau**/Aβ_1-42_. Tau and Aβ_1-42_ have been suspected to interact mutually during the AD pathology, through the cross-seeding of the amyloid core of Aβ_1-42_ and PHF6 and PHF6* of Tau.^61^ Our data suggest that PHF6 and PHF6* are involved in this cross-interaction but may not be involved in the seeding of Aβ_1-42_.

To conclude, we demonstrate that mimicking natural protein chaperones with small peptide drugs is an innovative and highly promising strategy to combat tauopathies. Furthermore, our lead β-hairpin mimic **β-Hsp90**-based on the physiological chaperone Hsp90, showing proteolytic stability and potent inhibitory activity on both Tau and Aβ_1-42_ aggregation mutually implicated in AD, offers the possibility to develop a very efficient dual-inhibitor. In complex diseases, where single-target drugs do not achieve the desired results, drug combinations or multitarget drugs treatment often result in higher effectiveness.^66,67^ Dual-targeting compounds such as dual inhibitors of Tau and Aβ have been proposed as new strategies to achieve better therapeutic benefits for AD.^34^

To the best of our knowledge there are no other Tau inhibitors (small molecules or peptides) that interact with the P1 region of Tau. This work paves the way to design new inhibitors of aggregation that do not affect the Tau-MTs physiological interaction. As proline rich region appears to be a critical region both in regulating Tau interaction with MTs as well as Tau aggregation, it is also important to study the behaviour of full-length Tau proteins rather than small peptide models that only contain the MT binding sequence.

Importantly, our β-hairpin mimics **β-Hsp90** and **β-Tau** provide valuable explorative tools for further and deeper studies of Hsp90/amyloid proteins and amyloid/amyloid cross-interactions (such as Hsp90/Tau, Tau/Aβ_1-42,_ Hsp90/Aβ_1-42,_ Tau/hIAPP and Hsp90/hIAPP). We are confident that these new results in the amyloid field will open new ways for designing peptide drug candidates not only to combat AD, but also, by extending the design from other chaperones acting on different amyloid proteins, to target other untreated amyloid diseases such as PD, HD, T2D or Amyotrophic lateral sclerosis (ALS). We consider that this study may help to develop urgently needed therapies for neurodegeneration and other age-dependent pathologies.

## Supporting information

Synthesis of compounds and CD spectra

## Methods

### General procedure for the Synthesis on Solid phase Peptide Synthesis Strategy (SPSS) for compounds β-Tau and β-Hsp90

All the reactions involved were agitated in plastic syringe tubes equipped with filters on an automated shaker on Rink Amide resin (0.2 mmol scale, loading 0.327 mmol/g, 600 mg). The coupling yields were monitored with the Fmoc-test procedure (reported below).

Removal of Fmoc group was performed in 20% (v/v) piperidine/DMF for 20 min twice. Capping steps were performed by treating the resin with the mixture of acetic anhydride (0.25 M) and NMM (0.25 M) in DMF solution for 20 min. After each reaction, the resin was washed with DMF (3 × 10mL), MeOH (3 × 10 mL) and DCM (3 × 10 mL) successively.

Rink Amide resin (600 mg, 0.2 mmol/g) was swelled in DMF for 1h before using. Natural amino acids and scaffold were coupled using different coupling reactive accordingly to their position:

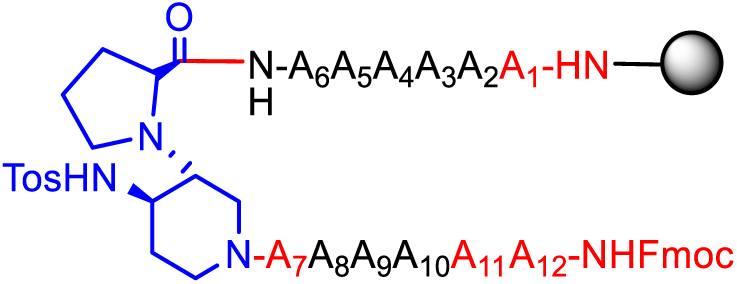

**Red positions**: The resin was suspended in DMF (4 mL) and collidine (17 eq). Fmoc-AA-OH (5 eq) DIC (5 eq) and Oxyma pure (5 eq) were solubilized in DMF/DCM 33% (v/v) (3 mL) and let under magnetic stirring for 7 minutes. Afterward, the solution containing the amino acid was poured into the reactor containing the resin and the mixture shaked for 16h at room temperature. (Positions 1, 7, 11 and 12)

**Black position**: Fmoc-AA-OH (5 eq) and HCTU (5 eq) were solubilized in 5 mL of DMF/NMM 20% (V/V). The prepared solution is added to the resin and shaked for 20 minutes. Afterward, the solvent was evaporated and the resin washed with DMF (1 x 5 mL) and the coupling procedure repeated. (Positions from 2 to 6 and from 8 to 10)

**Blue position**: The resin was suspended in DMF (4 mL) and collidine (17 eq). The piperidine-pyrrolidine β-turn scaffold^35^ (1.5 eq) DIC (1.5 eq) and OXYMA (1.5 eq) were solubilized in DMF/DCM 33% (3 mL) and let under magnetic stirring for 7 minutes. Afterward, this solution was poured into the reactor containing the resin and the mixture was shaked for 16h at room temperature.

At the end of the synthesis the peptides were cleaved from the resin shaking for 2 hours in the presence of 5 mL of an acidic solution containing trifluoracetic acid/H_2_O/TIPS/Phenol/Thioanisole; 87.5%/5%/2.5%/2.5%/2.5%. The liquid was poured in dry cold Et_2_O (40 mL) in ice bath by filtering it over a cotton pad. The peptides precipitated and isolated by centrifugation at 6000 rounds/min for 7 min. The pad was resuspended with ether and centrifuged again to remove the remaining TFA. The hairpin mimics were lyophilized and purified over reverse phase RP-HPLC.

### Synthesis of natural peptides H1, S4, S6 and S7

Peptides were synthesized using microwave assisted solid phase peptide synthesis performed on a Liberty Blue Microwave Automated Peptide Synthesizer (CEM Corporation, Matthews, NC, USA), following the standard protocols for Fmoc/tBu strategy (0.1 mmol scale, 300 mg, 0.327 mmol/g). Fmoc-deprotection cycles were respectively of 15 s (75 °C, 155 W) and 30 s (90 °C, 30 W). Couplings of Arg residues were performed in 1500 s (25 °C, 0 W) and 300 s (75 °C, 30 W). Couplings of His were performed in 240 s (50 °C, 35 W). Couplings of other residues are performed in 15 s (90 °C, 170 W) and 110 s (90 °C, 30 W). Peptide cleavage (3 h at room temperature) from the resin and deprotection of the amino acid side chains were performed by using the reagent K^14^ (trifluoracetic acid/phenol/water/thioanisole/1.2-ethanedithiol; 82.5/5/5/5/2.5) for 180 min. The liquid was poured in dry cold Et_2_O (40 mL) in ice bath by filtering it over a cotton pad. The peptides precipitated and isolated by centrifugation at 6000 rounds/min for 7 min. The pad was resuspended with ether and centrifuged again to remove the remaining TFA. The peptides were lyophilized and purified over RP-HPLC if necessary.

### Synthesis of natural peptides S4^short^ and S7^short^ via manual solid phase peptide synthesis

All the reaction involved were agitated in plastic syringe tubes equipped with filters on an automated shaker, on Rink Amide resin (0.2 mmol scale, loading 0.327 mmol/g, 600 mg).

Removal of Fmoc group was performed in 20% (v/v) piperidine/DMF for 20 min twice. Capping steps were performed by treating the resin with the mixture of acetic anhydride (0.25 M) and NMM (0.25 M) in DMF solution for 20 min. After each reaction, the resin was washed with DMF (3 × 10mL), MeOH (3 × 10 mL) and DCM (3 × 10 mL) successively.

Rink Amide resin (600 mg, 0.2 mmol/g) was swelled by DMF for 1h before using. The loading was performed by suspending the resin with DMF (4 mL) and collidine (17 eq). Fmoc-AA-OH (5 eq) DIC (5 eq) and Oxyma pure (5 eq) were solubilized in DMF/DCM 33% (v/v) (3 mL) and let under magnetic stirring for 7 minutes. Afterward, the solution containing the amino acid was poured into the reactor containing the resin and the mixture shaked for 16h at room temperature. Every following coupling using Fmoc-protected natural amino acid was carried out twice to get satisfactory yields using Fmoc-AA-OH/HCTU (2.5/2.5 eq) in NMM/DMF (20%V/V, 4 ml).

Peptides were cleaved from the resin by shaking for 2 hours with an acidic mixture (TFA/water/TIPS/Thioanisole; 95%/2.5%/1.25%/1.25%) and repeated the same procedure explained in General procedure **A**.

### Fmoc loading Test phase (Fmoc-test)

From 5 to 10 mg of dry resin was shaken in DMF/Piperidine 20% solution (1 mL) for 30 minutes. The resin was sedimented by centrifugation and 1 mL of the supernatant was added to 9 mL of DMF and mixed well. This final solution was used for the UV analysis measuring the absorbance at 301 nm versus a DMF blank (triplicate analysis).

For the quantification of resin loading, we used the general formula: L = (A_301_× V × d)/(E_c_ × w × M). Where: L = Resin loading; A_301_ = UV Absorbance at 301 nm; V = Volume of the cleavage solution = 1 mL; d = Dilution = 10; E_c_ = Extinction coefficient = 7800 mL/mmol*cm; w = Width of the cuvette = 1 cm; M = Weight of the resin sample in g.^68^ Substituting the extinction coefficient, volume, dilution and cell width into the general formula results in this formula which can be used to calculate the resin loading.

L = (100 × A_301_)/(7.8 × M(mg)) in units of mmols/gram.

### Circular Dichroism Spectroscopy

Peptidomimetics were dissolved in MQ water to a concentration of 500 μM as stock solutions. Before measurement, each compound was diluted to 125 µM concentration with 20 mM PB (pH 7.2) buffer or MeOH into a cuvette with a pathlength of 1 mm. The CD spectra were recorded on a J-815 spectropolarimeter (JASCO, Tokyo, Japan) from 190 to 260 nm at 20 and 37°C and a scan rate of 50 nm/min (accumulation n=3). Each CD spectrum was corrected by subtracting the corresponding baseline (PB buffer 20 mM or MeOH). Data processing was performed using Solver in Excel software (Microsoft). Deconvolution process was executed according to literature.^41^

### Fluorescence-Detected ThT Binding Assay (Ac-PHF6*-NH_2_)

Thioflavin-T and Heparin sodium salt (H-3149, average MW 18 kDa) were obtained from Sigma-Aldrich. Ac-PHF6*-NH_2_ was self-produced according to the general SPPS protocol of **S4^short^ and S7^short^** and purified by HPLC. The purity was confirmed by LC-MS and HRMS. Ac-PHF6*-NH_2_ was dissolved in pure hexafluoro-isopropanol (HFIP) at a concentration of 1 mM and incubated for 10 minutes at room temperature to dissolve any preformed aggregates. Next, HFIP was evaporated under a stream of dry nitrogen gas followed by vacuum desiccation for at least 3 hours. The resulting thin film was dissolved in 20 mM MOPS buffer (pH 7.4), sonicated for 1 min and vortexed for 2 min to get fully disperse clean solution of AcPHF6*-NH_2_ (500 µM). Stock solutions of compounds to test were dissolved in water (10, 2 and 0.2 mM). Thioflavin-T binding assays was used to measure the formation of fibrils in solution using a plate reader (Fluostar Optima, BmgLabtech) and standard 96-wells flat-bottom black microtiter plates (final volume 200 µL) in combination with a 440 nm excitation and 480 nm emission filters.ThT assay was started by adding 2 µL of a 100 µM heparin solution (Heparin sodium salt H-3149, average MW 18 kDa) to a mixture containing 25 µM of Ac-PHF6*-NH_2_, 20 μM ThT in 20 mM MOPS pH 7.4 buffer. The concentration of Ac-PHF6*-NH_2_ was held constant at 25 μM for all experiments and inhibitors were added to yield compound/ Ac-PHF6*-NH_2_ ratios of 5/1, 1/1 and 0.1/1. The ThT assays were performed in triplicate.

### Production and purification of recombinant Wt-Tau and TauΔK280

Prokaryotic expression plasmids were based on human adult Tau (Tau441wt) in a pET-3d vector.^69,70^ pET-3d-tau plasmids expressing human Tau protein (Wt-Tau) and the ΔK280 deletion variant (TauΔK280), were transformed into Escherichia coli BL21(DE3)pLysS cells for expression. Cells were grown, induced and harvested as previously described.^71^ For TEM analysis, native Tau was purified from the cell extract by sequential anion exchange and phosphocellulose chromatography as previously described.^69^ The Tau protein eluate was dialyzed against water and concentrated with Vivaspin® (15R, 2,000 MWCO, Sartorius, UK). Protein concentrations were determined by a bicinchoninic acid (BCA) assay (Thermo-Fisher Scientific, USA) using bovine serum albumin as a standard. For ThT assays, Tau441wt protein was also purified using heat denaturation following established protocols^70^ and dialyzed against 50 mM ammonium bicarbonate buffer before being lyophilized.

### Fluorescence-Detected ThT Binding Assay (Wt-Tau441 and Tau441-ΔK280)

Thioflavin-T and Heparin sodium salt (H-3149, average MW 18 kDa) were obtained from Sigma-Aldrich. Lyophilized full length Wt-Tau441 and Tau441 ΔK280 were diluted to 40 μM in NaPi 25 mM, NaCl 25 mM, and EDTA 2.5 mM pH 6.8. Stock solutions of compounds to test were dissolved in water. ThT fluorescence was measured to evaluate the development of Tau fibrils over time using a fluorescence plate reader (Fluostar Optima, BMG labtech) with 384-wells flat-bottom black plates (final volume in the wells of 40 μL). Experiments were started by adding Heparin (0.0625 μM) to each well containing the resulting Tau solution (final Tau concentration equal to 10 μM), thioflavin-T (25 μM) with and without the compounds to test at different concentrations (50, 10 and 1 μM, ratios compounds/Wt-Tau equal to 5/1, 1/1 and 0.1/1) in NaPi 25 mM, NaCl 25 mM, EDTA 2.5 mM pH 6.8. The ThT fluorescence intensity of each sample (performed in triplicate) was recorded over time with 440/480 nm excitation/emission filters set for 30 h under continuous agitation (orbital shaking) at 37 °C on plates sealed with a transparent film. The ability of compounds to inhibit Wt-Tau aggregation was assessed considering the time of the half-aggregation (*t1/2*) and the intensity of the experimental fluorescence plateau (F), both values were obtained by fitting the obtained kinetic data to a Boltzmann sigmoidal curve using GraphPad Prism 5. The relative *t_1/2_* is defined as the experimental *t_1/2_* in the presence of the tested compound relative to the one obtained without the compound and is evaluated as the following percentage: [*t_1/2_* (Tau + compound)] / *t_1/2_* (Tau) × 100. The relative F is defined as the intensity of experimental fluorescence plateau observed with the tested compound relative to the value obtained without the compound and is evaluated as the following percentage: (F(Tau + compound) / F(Tau) × 100. Curves of the tested compounds are fitted to a Boltzmann sigmoidal model and represented relative to the maximal fluorescence of the control experiment.

### Fluorescence-Detected ThT Binding Assay (Aβ_1−42_)

Aβ_1−42_ was purchased from Bachem and Thioflavin-T was obtained from Sigma. The peptide was dissolved in an 1% aqueous ammonia solution to a concentration of 1 mM and then, just prior to use, was diluted to 0.2 mM with 10 mM Tris-HCl and 100 mM NaCl buffer (pH 7.4). Stock solutions of compounds to test were dissolved in water. ThT fluorescence was measured to evaluate the development of Aβ_1−42_ fibrils over time using a fluorescence plate reader (Fluostar Optima, BMG labtech) with standard 96-well black microtiter plates (final volume in the wells of 200 μL). Experiments were started by adding the peptide (final Aβ_1−42_ concentration equal to 10 μM) into a mixture containing 40 μM ThT in 10 mM Tris-HCl and 100 mM NaCl buffer (pH 7.4) with and without the compounds at different concentrations (100 and 10 μM, compound/ Aβ_1−42_ ratios of 10/1 and 1/1) at room temperature. The ThT fluorescence intensity of each sample (performed in duplicate) was recorded with 440/480 nm excitation/emission filters set for 42 h performing a double orbital shaking of 10 s before the first cycle. The ability of compounds to inhibit Aβ_1−42_ aggregation was assessed considering the time of the half aggregation (*t_1/2_*) and the intensity of the experimental fluorescence plateau (F), both values were obtained by fitting the obtained kinetic data to a Boltzmann sigmoidal curve using GraphPad Prism 5. The relative *t_1/2_*is defined as the experimental *t_1/2_* in the presence of the tested compound relative to the one obtained without the compound and is evaluated as the following percentage: [*t_1/2_* (Aβ + compound)] / *t_1/2_* (Aβ) × 100. The relative F is defined as the intensity of experimental fluorescence plateau observed with the tested compound relative to the value obtained without the compound and is evaluated as the following percentage: F(Aβ + compound) / F(Aβ) × 100. Curves of the tested compounds are fitted to a Boltzmann sigmoidal model and normalized to the maximal fluorescence of the control experiment.

### Fluorescence-Detected ThT Binding Assay (hIAPP)

Thioflavin-T was obtained from Sigma. hIAPP, purchased from Bachem, was dissolved in pure hexafluoro-isopropanol (HFIP) at a concentration of 1 mM and incubated for 1 hour at room temperature to dissolve any preformed aggregates. Next, HFIP was evaporated under a stream of dry nitrogen gas followed by vacuum desiccation for at least 3 hours. The resulting peptide film was then dissolved in DMSO to obtain stock solutions of hIAPP (0.2 mM) and stock solutions of compounds to test were dissolved in water. Thioflavin-T binding assays were used to measure the formation of fibrils in solution or in the presence of membranes over time. A plate reader (Fluostar Optima, BmgLabtech) and standard 96-wells flat-bottom black microtiter plates in combination with a 440 nm excitation and 480 nm emission filters were used. The ThT assay was started by adding 5 μL of a 0.2 mM hIAPP stock solution to a mixture of 10 μM ThT and 10 mM Tris/HCl, 100 mM NaCl at pH 7.4, containing the compounds to test (50 and 5 μM, compound/ hIAPP ratios of 10/1 and 1/1) at room temperature. The ThT assays were performed once. The curves of the tested compounds are fitted to a Boltzmann sigmoidal model and normalized to the maximal fluorescence of the control experiment.

### Transmission Electron Microscopy (TEM) for Wt-Tau_441_

Wt-Tau_441_ solubilization immediately before use: Lyophilized Tau was dissolved in buffer NaPi (Na_2_HPO_4_ and NaH_2_PO_4_ 25 mM, NaCl 25 mM, EDTA 2.5 mM, pH 6.6) to a final concentration of 40 µM (stock solution).

Sample preparation: Experiments were started by adding 10 µL of Buffer, 10 µL of Tau solution (Final concentration of 10 µM), 10 µL of desired active compound (either 200µM or 40µM to evaluate 5:1 and 1:1 ratios), and 10 µL Heparin solution 0.25 µM solubilized in NaPi buffer (final concentration 0.0625 µM). In blank analysis, 10 µL of active compound was replaced by 10 µL MQ H_2_O. These solutions were maintained at 37°C for 96h at 1400 rpm.

The samples were prepared on carbon-coated copper grids (ECF200-Cu, 200 mesh, Science Services, Munich, Germany). The grids were pre-treated with argon in a plasma cleaner (Diener Electronics, Ebhausen, Germany). The sample (1.2 µL) was applied onto the grids, and after a sedimentation time of 5 minutes, the excess suspension was removed with filter paper. Grids were then stained with 1% acetate uranyl solution (1.2 µL) for 5 minutes, excess stain was removed with filter paper and washed thoroughly with 3 x 10 µL of Milli-Q water. The prepared grids were analyzed with a JEOL JEM-2200FS electron microscope (JEOL, Freising, Germany), a cold field emission electron gun, and an applied acceleration voltage of 200 kV. A bottom-mounted Gatan OneView camera (Gatan, Pleasanton, CA, USA) was used for digital recording. The images were processed using the image-processing system Digital Micrograph GMS3 (Gatan, Pleasanton, CA, USA) and the image editing software ImageJ.

### Transmission Electron Microscopy (TEM) for Aβ_1-42_

Aβ_1-42_ Pre-treatment for storage: Aβ_1-42_ was dissolved in 1mg/mL in NaOH 60 mM and sonicated for 5 minutes. Afterward the solution was centrifugated at 5000 rpm for 5 minutes and lyophilized.^72^

Aβ_1-42_ Pre-treatment immediately before use: pre-treated Aβ_1-42_ was dissolved in an HCl 60 mM obtaining a final concentration of 200 µM.

Sample preparation: Experiments were started by adding the peptide (final Aβ_1−42_ concentration equal to 10 μM) into a solution containing the active compound in the appropriate ratio (100 or 10 μM) into 20 mM phosphate buffer pH 7.4, containing 1 mM NaN_3_ and 200 µM EDTA. The mixture was then shacked for 24h at 37°C at 700 rpm.

Transmission electron microscopy. The samples were prepared on carbon-coated copper grids (ECF200-Cu, 200 mesh, Science Services, Munich, Germany). The grids were pre-treated with argon in a plasma cleaner (Diener Electronics, Ebhausen, Germany). The sample (2 µL) was applied onto the grids, and after a sedimentation time of 5 minutes, the excess suspension was removed with filter paper. Grids were then stained with 1% acetate uranyl solution (2 µL) for 5 minutes, excess stain was removed with filter paper and washed thoroughly with 3 x 10 µL of Milli-Q water. The prepared grids were analyzed with a JEOL JEM-2200FS electron microscope (JEOL, Freising, Germany), a cold field emission electron gun, and an applied acceleration voltage of 200 kV. A bottom-mounted Gatan OneView camera (Gatan, Pleasanton, CA, USA) was used for digital recording. The images were processed with the image-processing system Digital Micrograph GMS3 (Gatan, Pleasanton, CA, USA) and the image editing software ImageJ.

**Cell lines.** PC12 cells (originally obtained from J.A. Wagner, Harvard Medical School) were cultured in serum-DMEM (DMEM supplemented with 10% fetal bovine serum and antibiotics (100 U/ml penicillin and 100 µg/ml streptomycin)) at 37°C with 10% CO_2_ in a humidified incubator. For live cell imaging, cells were transfected with the respective pRc/CMV expression vector using Lipofectamine 2000 (Thermo-Fisher Scientific, USA) as previously described.^73^

### Metabolic activity and cytotoxicity profiling using a combined LDH and MTT Assay

PC12 cells were cultured in 96-well plates at 1×10^4^ cells/well in 50 µL of serum-reduced medium supplemented with 100 ng/mL 7S mouse NGF to induce neuronal differentiation. Cells were incubated for 48, test compounds were added in an additional volume of 50 µL, and incubation was continued for 20 hours. For the LDH assay, 50 µL medium from each well of the assay plate was transferred to a separate 96-well plate. To quantify LDH release, 50µL of LDH reagent (4 mM iodonitrotetrazolium chloride (INT), 6.4 mM beta-nicotinamide adenine dinucleotide sodium salt (NAD), 320 mM lithium lactate, 150 mM of 1-methoxyphenazine methosulfate (MPMS) in 0.2 M Tris-HCl buffer, pH 8.2) was added. Absorbance was measured at 490 nm using a Thermomax Microplate Reader operated with SoftMaxPro Version 1.1 (Molecular Devices Corp., Sunnyvale, CA, U.S.A.). For the MTT assay, MTT reagent (3,(4,5.dimethylthiazol-2-yl)2,5-diphenyltetrazolium bromide) was added to the wells of the remaining assay plate at a final concentration of 1 mg/ml MTT. Cells were incubated for 2 hours before the reaction was stopped by adding 50 μl of lysis buffer (20% (wt/vol) sodium dodecyl sulfate in 1:1 (vol/vol) *N,N*-dimethylformamide/water, pH 4.7). After overnight incubation at 37 °C, optical densities of the formazan product were determined at 570 nm. MTT conversion measurements were normalized to the optical densities of negative control wells. All experiments were performed in triplicates on two independent plates.

### Live-cell imaging and Fluorescence Decay After Photoactivation (FDAP)

FDAP experiments were performed essentially as previously described.^60^ Briefly, cells expressing wild-type tau or TauΔK280 were plated on 35-mm glass-bottom culture dishes (MatTek, USA), transfected, and neuronally differentiated by medium exchange to serum-reduced DMEM containing 100 ng/mL 7S mouse NGF. After 3 days, the medium was exchanged to serum-reduced DMEM without phenol red with NGF, and the respective compound (or DMSO for carrier control) were added at the desired concentration. After 20 hours, live cell imaging was performed using a laser scanning microscope (Nikon Eclipse Ti2-E (Nikon, Japan)) equipped with a LU-N4 laser unit with 488-nm and 405-nm lasers and a Fluor 60× ultraviolet-corrected objective lens (NA 1.4) enclosed in an incubation chamber at 37°C and 5% CO_2_. Photoactivation was performed with a 405-nm laser using the microscope software (NIS-Elements version AR 5.02.03 (Nikon, Japan)). A series of consecutive images was acquired at a frequency of 1 frame/s, and 112 images were collected per activated cell at a resolution of 256×256 pixels. Effective diffusion constants were determined by fitting the fluorescence decay data from the photoactivation experiments using a one-dimensional diffusion model function for FDAP. A reaction-diffusion model was used to estimate the association rate k*_on_ and the dissociation rate k_off_ constant of tau binding.

### Pronase stability assays

Prepare the assay buffer by combining 50 mM Tris, 10 mM CaCl_2_, 150 mM NaCl, and 0.05% (w/v) Brij 35, adjusting the pH to 7.5. Dissolve the peptide in 200 µL of assay buffer to achieve a concentration of 400 µM. Place the solution in a thermomixer set at 37°C with stirring. Dilute pronase to a concentration of 5 µg/mL in 200 µL of assay buffer. Place the solution in a thermomixer set at 37°C with stirring. Combine the peptide solution (200 µL) with the pronase solution (200 µL) in an Eppendorf tube. The final concentration of pronase should be 2.5 µg/mL, and the final concentration of the peptide should be 200 µM. Leave the reaction mixture in the thermomixer at 37°C. Monitor the reaction progress by LC-MS at various time points: 0 min, 10 min, 30 min, 1 h, and 2 h. At each time point, transfer 40 µL of the reaction solution to an HPLC vial containing 6 µL of 10% TFA (to deactivate the enzyme). Repeat this procedure for each time point.

### Preparation of Wt (2N4R) ^15^N-Tau

The full-length isoform of Tau (2N4R, Uniprot ID P10636) enriched with the 15-Nitrogen stable isotope was prepared based on a previously published method.^74^ Cloning was accomplished with a pNG2 vector (Novagen). The expression plasmid containing the gene encoding for Tau was subsequently used to transform BL21(DE3) *Escherichia coli* cells, which were grown to an optical density (at 600 nm) of 0.8 in Luria Broth at 37 °C. Cells were pelleted by centrifugation, washed, and resuspended in M9 minimal medium (6 g/L Na_2_HPO_4_, 3 g/L KH_2_PO_4_, 0.5 g/L NaCl, 4 g/L glucose, pH 7.0) containing 0.5 g ISOGRO-^15^N (Sigma Aldrich) and 2 g/L ^15^N-ammonium chloride as the nitrogen source. Induction of protein expression was accomplished by the addition of IPTG to 0.5 mM followed by overnight incubation at 37 °C.

The cell pellet was collected and subjected to lysis using a French cell press (lysis buffer contains 20 mM MES, 1 mM EGTA, 0.2 mM MgCl_2_, 5 mM DTT, 1 mM PMSF, 1 mg/mL lysozyme, 10 µg/mL DNAse, pH 6.8). Salt, 500 mM NaCl, was added to the lysate, which was then boiled to denature proteins. The lysate was cleared of cell debris, DNA, and precipitated proteins by high-speed centrifugation (12700 × g, 40 min, 4 °C). Residual nucleic acids were precipitated by incubating the supernatant with 20 mg/mL streptomycin. The precipitated nucleic acids were separated by centrifugation. The resulting supernatant was incubated with 0.36 g/mL ammonium sulfate then subjected to centrifugation to collect a protein pellet mostly containing Tau protein. The protein pellet was resuspended and dialyzed into a dialysis buffer (20 mM MES, 1 mM EDTA, 2 mM DTT, 50 mM NaCl, pH 6.8). Tau was purified from the mixture using cation exchange chromatography (MonoS 10/100, 2 mL/min with a buffer gradient of 50 mM to 1 M NaCl in 20 mM MES, 1 mM EGTA, 2 mM DTT, pH 6.8), and size exclusion chromatography (Superdex 75 26/600, 2 mL/min in phosphate buffered saline with additional 500 mM NaCl). The purified Tau protein was dialyzed overnight against 50 mM NaPi pH 6.8, concentrated and flash-frozen in aliquots, and stored at −80 °C until they were used in NMR experiments. The protein concentration was determined by bicinchoninic acid assay following the manufacturer’s instructions (Pierce).

### NMR Spectroscopy: 2D ^1^H-^15^N HSQC titrations of ^15^N-Tau

The lyophilized compounds **β-Tau** and **β-Hsp90** were resuspended in water to a concentration of 1000 µM, followed by dilution to a final buffer composition of 50 mM NaPi pH 6.8, 10 mM NaCl, 1 mM DTT and a working concentration of 400 µM compound. For each titration series, a 200 µL solution of 2N4R ^15^N-Tau (20 µM) was prepared in 50 mM NaP pH 6.8, 10 mM NaCl, 1 mM DTT, 10% D_2_O and incremented with 20 µM, 40 µM, and 100 µM of **β-Tau** and **β-Hsp90**.

Heteronuclear single quantum coherence (^1^H-^15^N HSQC) spectra of Tau with and without compounds were collected at 5°C using 900 MHz and 800 MHz spectrometers equipped with cryoprobes. HSQC spectra were processed using TopSpin version 3.6.2 (Bruker) and analyzed using Sparky.^75^ The Tau binding profiles were represented by peak intensity ratios (I/I_0_) and chemical shift perturbations (CSP). The I/I_0_ ratio was calculated by dividing the peak amplitude (I) of Tau amide peaks in the presence of the compound by the corresponding peak intensity (I_0_) in the absence of the compound. CSPs were calculated using the equation 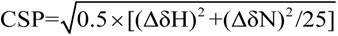, where ΔδH and ΔδN correspond to the chemical shift differences of H and N. CSPs were smoothed with a 3-residue averaging window.

### Preparation of ^15^N-Aβ_1−42_

Recombinant ^15^N-Aβ1−42 (rPeptide) was resuspended in 1% NH_4_OH at 1 mg/mL concentration, lyophilized overnight and stored at -20°C. Aliquots containing 70 µg ^15^N-Aβ1−42 were resuspended in 300 µL of 20 mM NaPi pH 7.2, 10% D_2_O buffer, to obtain a concentration of 50 µM. ^15^N-Aβ1−42 samples containing 50 µM or 500 µM **β-Hsp90** were prepared by adding the appropriate volume of a 1 mM **β-Hsp90** stock solution in the same buffer.

### 2D ^1^H-^15^N HSQC titrations of ^15^N-Aβ_1-42_

5 mm Shigemi tubes were used for NMR experiments. 2D ^1^H-^15^N HSQC spectra of ^15^N-Aβ_1−42_ in the absence and in the presence of **β-Hsp90** at 1:1 and 10:1 ratios were collected at 5°C using a 500 MHz spectrometer equipped with a cryoprobe. NMR spectra were processed with nmrPipe and analyzed using Sparky.

## Acknowledgements

This research has received funding from the European Union’s Horizon 2020 research and innovation program H2020-MSCAITN-2019-EJD: Marie Skłodowska-Curie Innovative Training Networks (European Joint Doctorate)—Grant Agreement No: 860070—TubInTrain (DDL, NB, RB, SO) M.Z. and L.R. thank Maria-Sol Cima-Omori for the expression and purification of 2N4R ^15^N-Tau. M.Z. was supported by the European Research Council (ERC) under the EU Horizon 2020 research and innovation programme (grant agreement numbers 787679 and 101069214).

I.G. has been supported by Erasmus+ program (Pisa University).

Robert Thai (CEA Paris Saclay) is thanked for his contribution to the LC-MS analysis of pronase stability assays.

## Author contributions

D.D.L. performed the synthesis, NMR and CD analyses, pronase stability and participate to ThT assays on Tau under the co-supervision of N.T. and S.O. and the preparation of TEM specimens and their analysis under the supervision of V.D.

N.B. performed and analyzed MTT assays and quantitative live-cell imaging under the supervision of R.B.

J.K. performed and analyzed the ThT assays (Wt-Tau, TauΔK280, Aβ_1-42_, hIAPP)

L.R. performed and analyzed the NMR titrations of WT (2N4R) ^15^N-Tau with **β-Hsp90** and **β-Tau**, under the supervision of M.Z.

O.L. performed and analyzed the NMR titrations of Aβ_1-42_ ^15^N-Tau with **β-Hsp90**

I.G. synthesized AcPHF6* and performed the ThT assays of AcPHF6* under the supervision of D.D.L. and N.T.

J.L. produced Wt-Tau for ThT assays and TEM

Y.H. and I.E. performed and analyzed TEM imaging of Wt-Tau and Aβ_1-42_

V.D. designed, supervised and analyzed TEM experiments

N.S. paid and supervised D.D.L. for TEM during his post-doctoral period at Bielefeld University

M.L.G. co-supervised D.D.L. during his period at Università degli Studi di Milano

N.T. designed the compounds and co-supervised D.D.L.

R.B. supervised N.B. for MTT assays and quantitative live-cell imaging

S.O. supervised D.D.L. and N.B. and managed the global project and the preparation of the manuscript

## Competing interests

The authors declare no competing interests.

Supplementary information. The online version contains supplementary material available at

**Extended data Fig. 1:**
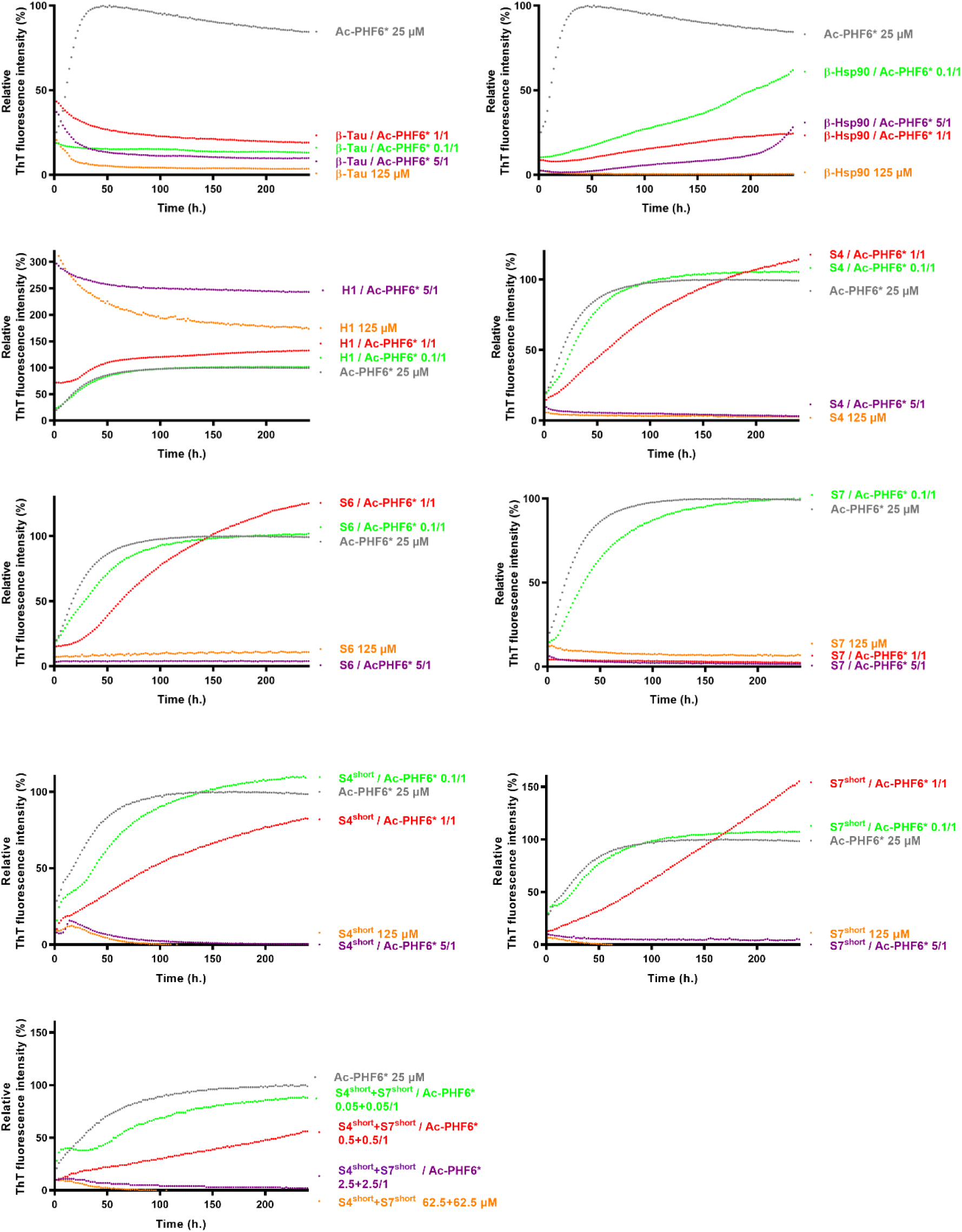
Mean curves of ThT fluorescence assays over time showing Ac-PHF6* fibrillization (25 µM) in the absence (grey curve) and in the presence of compounds **β-Tau**, **β-Hsp90**, **H1**, **S4**, **S6**, **S7**, **S4^short^, S7^short^**, **S4^short^** plus **S7^short^**at compound/Ac-PHF6* ratios of 5/1 (purple curves), 1/1 (red curves) and 0.1/1 (green curves). The control curves of compounds alone are represented in orange lines. (mean curves of triplicates).

**Extended data Fig. 2:**
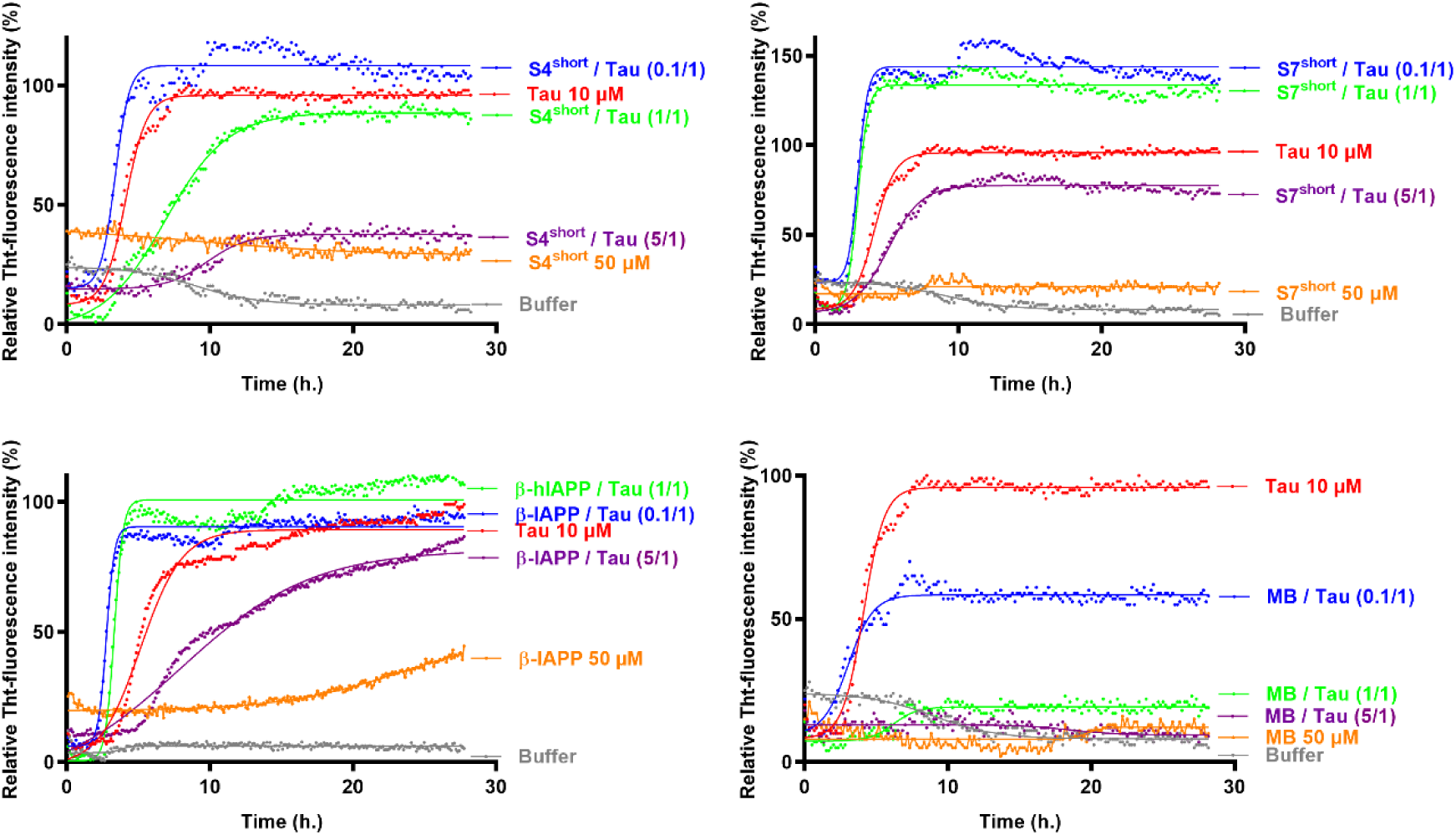
Representative curves of ThT fluorescence assays over time showing Wt-Tau fibrillization (10 µM) induced by heparin (Tau/heparin ratio of 160/1) in the absence (red curve) and in the presence of **β-Tau**, **β-Hsp90**, **S4**, **S7**, **S4^short^**, **S7^short^**, **MB and β-hIAPP** at compound/Wt-Tau ratios of 5/1 (purple curves), 1/1 (green curves) and 0.1/1 (blue curves). The control curves are represented in orange lines and buffer in grey.

**Extended data Fig. 3:**
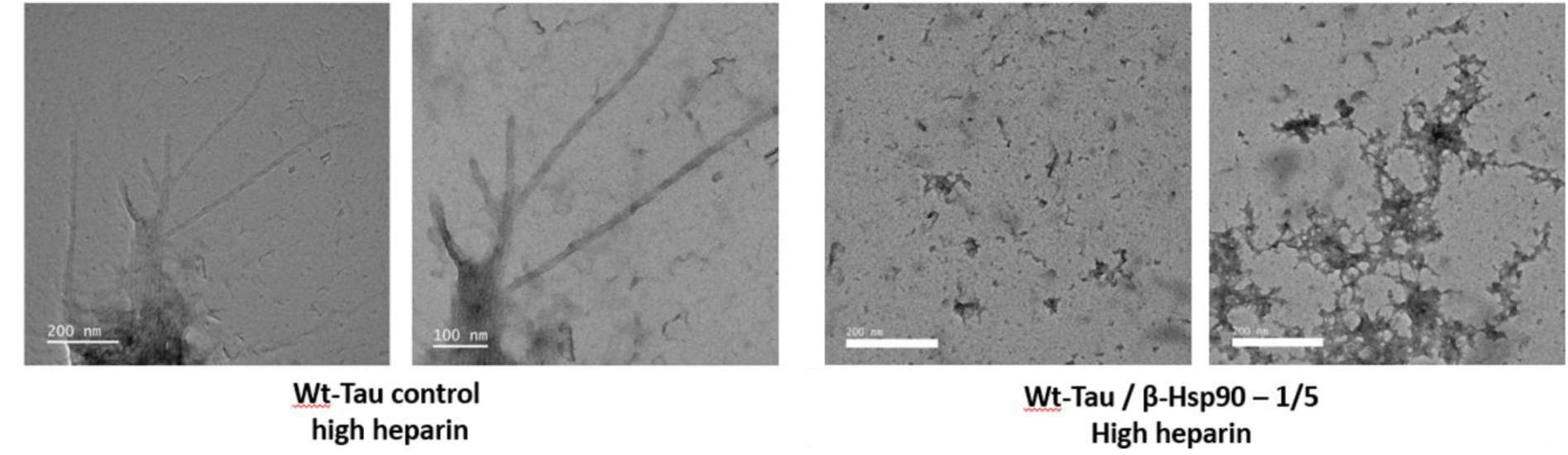
Transmission electron micrographs of Wt-Tau fibrils formed at a higher Heparin concentration (4:1 ratio) and the amorphous aggregates of Wt-Tau in the presence of 5 equiv. of **β-Hsp90**.

**Extended data Fig. 4:**
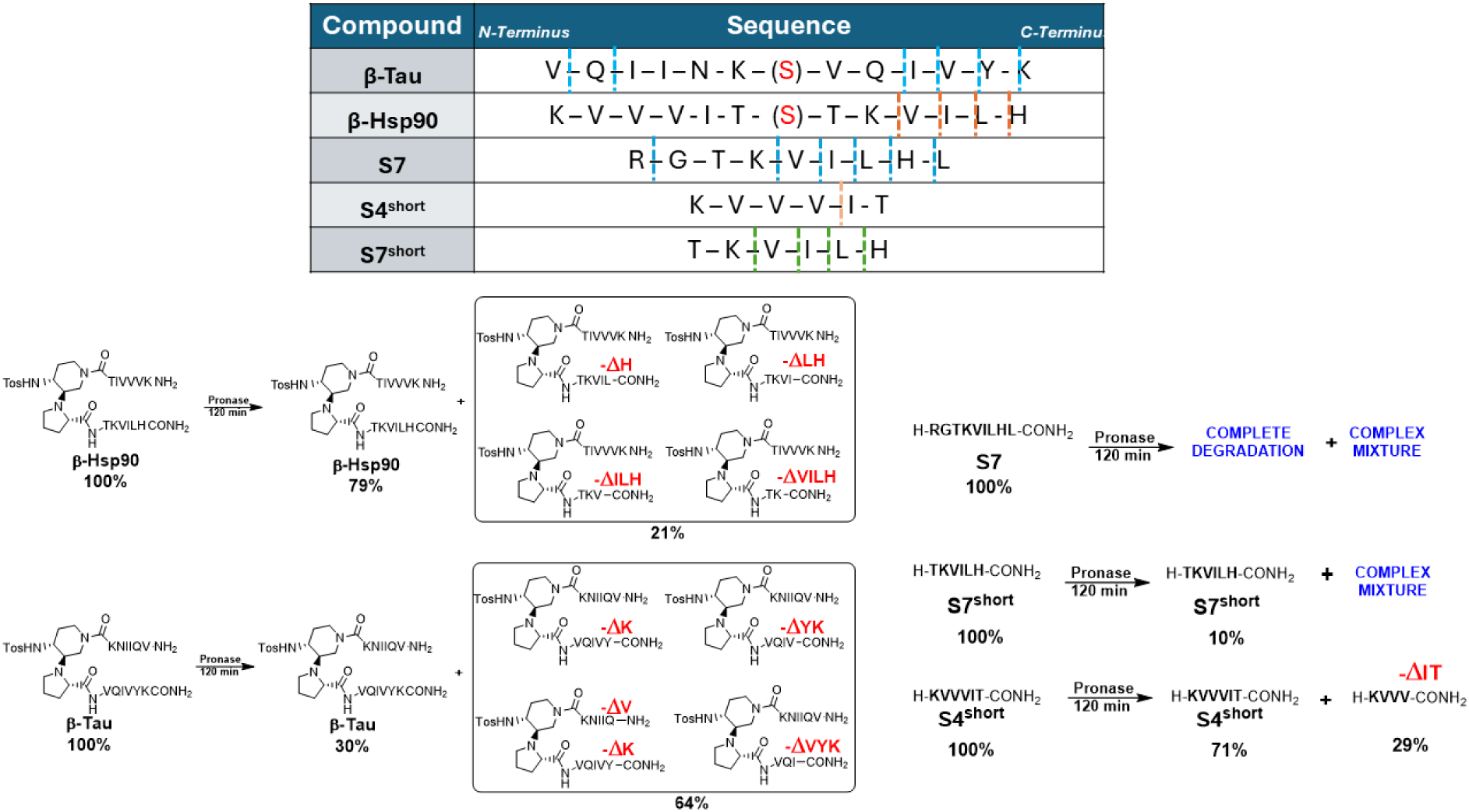
Representation of the principal cleavages induced by the proteases present in the pronase mixture on **β-Tau**, **β-Hsp90**, **S7**, **S4^short^** and **S7^sho^**^rt^. Quantification of subproducts derived from compounds **S7** and **S7^sho^**^rt^ was not possible due to overlap of the picks.

**Extended data Fig. 5:**
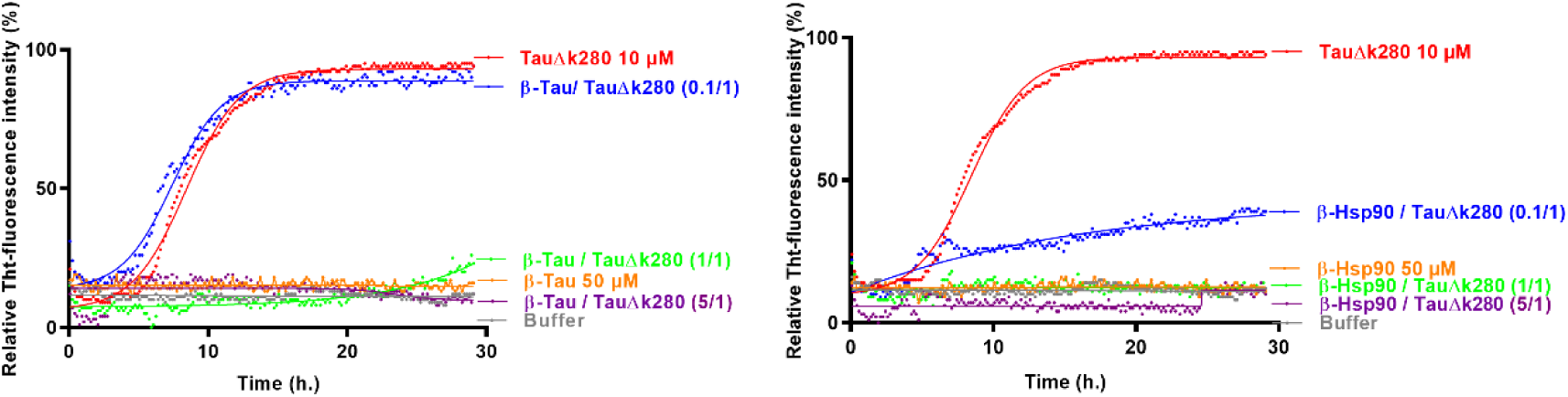
Representative curves of ThT fluorescence assays over time showing Tau-ΔK280 fibrillization (10 µM) in the absence (red curve) and in the presence of compounds **β-Tau** and **β-Hsp90** at compound/Tau-ΔK280 ratios of 5/1 (purple curves), 1/1 (green curves) and 0.1/1 (blue curves). The control curves are represented in orange lines and buffer in grey.

**Extended data Fig. 6:**
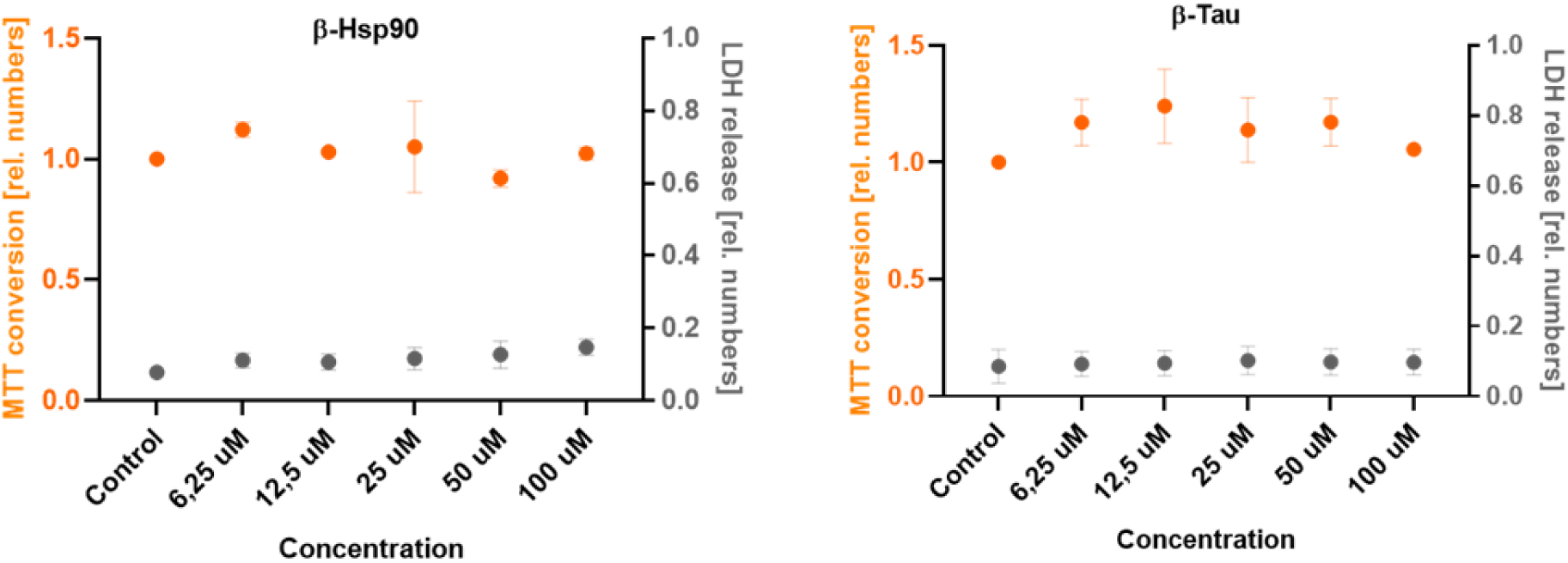
Toxicity and metabolic activity profiles of **β-Hsp90** and **β-Tau** using a combined LDH- and MTT assays. Neuronally differentiated PC12 cells were exposed to the respective compounds. Each compound and concentration were measured in triplicates on two independent assay plates while mean ± SEM of the two plates is shown. Statistical assessment of metabolic activity and cytotoxicity compared to carrier control was performed by One-way ANOVA using a Dunnet’s post-hoc test. Neither membrane damage nor metabolic impairment was observed with compounds’ concentrations up to 100 µM.

**Extended data Fig. 7:**
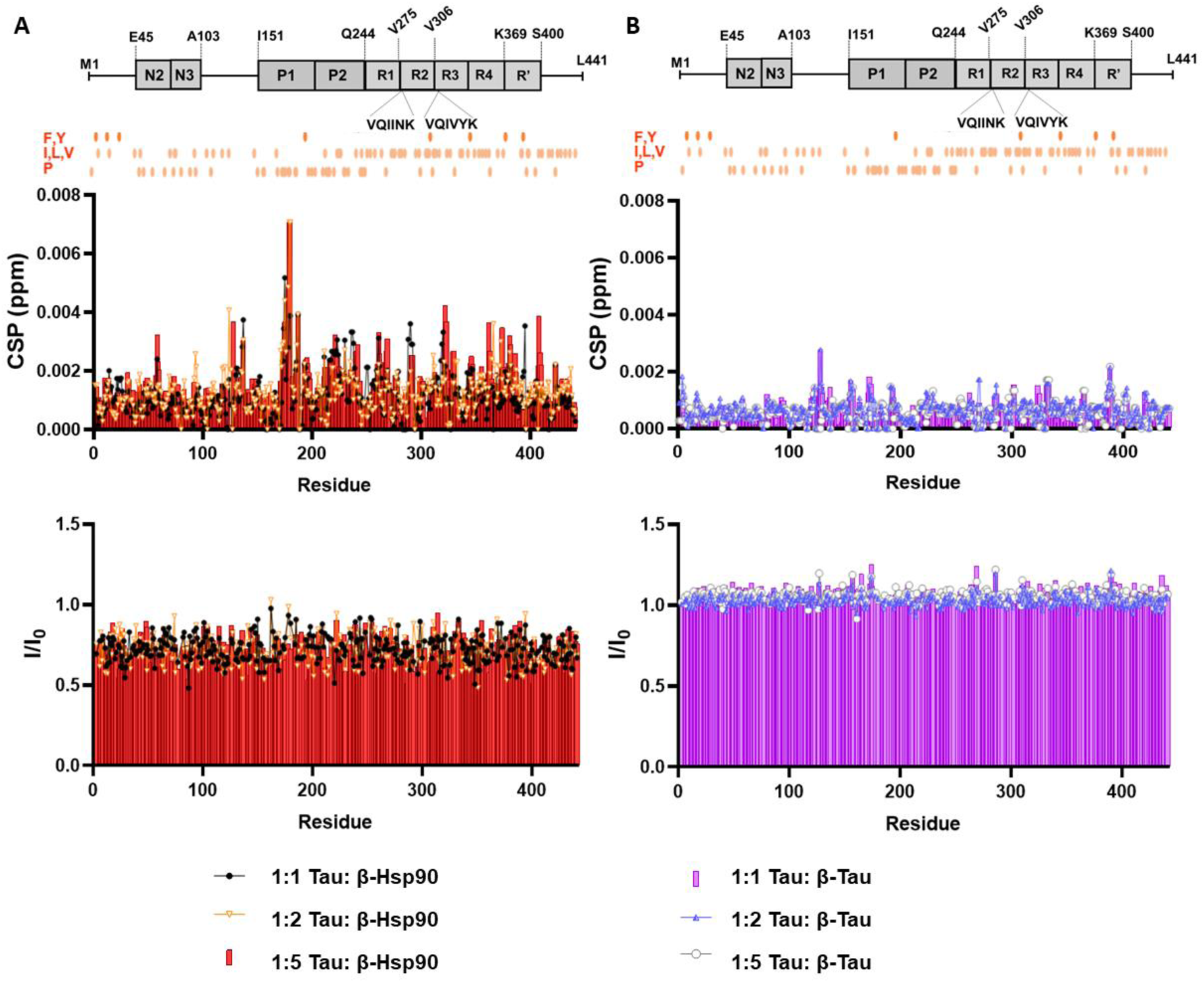
Titrations of ^15^N HSQC of 2N4R ^15^N-Tau (20 μM) with β-Tau and β-Hsp90 in buffer solution (50 mM NaPi pH 6.8, 10 mM NaCl, 1 mM DTT, 10% D_2_O) at 5°C (900 MHz or 800 MHz). Different ratios Wt-Tau_441_/compounds = 1/5, 1/2 and 1/1 were used. I/I_0_ values (I represents NMR signal intensity in the presence of peptide, and I_0_ corresponds to conditions in the absence of compounds) and CSPs due to β-Hsp90/ β-Tau are provided. A) CSP and I/I_0_ values of Tau residues in titrations with β-Tau (1/5, 1/2, and 1/1 ratios). B) CSP and I/I_0_ values of Tau residues in titrations with β-Hsp90 (1/5, 1/2, and 1/1 ratios). Indicated in orange circles (A and B) are the locations of selected hydrophobic amino acids (F,Y,I,L,V,P) in the sequence of 2N4R Tau. CSP and I/I_0_ values at the mole ratio 1/5 are identical to those in Figure 5.

**Extended data Fig. 8:**
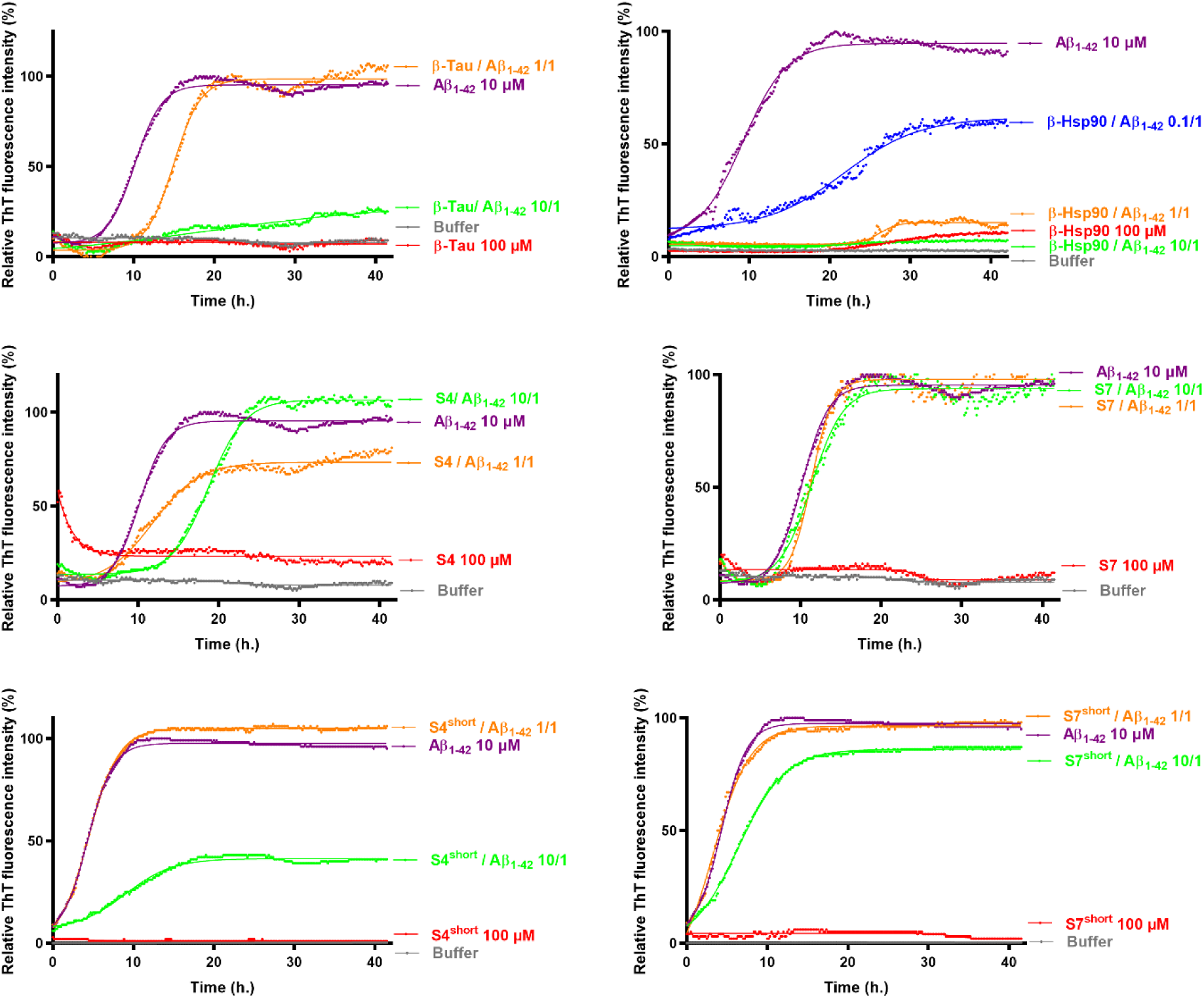
Representative curves of ThT fluorescence assays over time showing Aβ_1-42_ fibrillization (10 µM) in the absence (purple curve) and in the presence of **β-Tau**, **β-Hsp90**, **S4**, **S7**, **S4^short^** and **S7^short^** at compound/Aβ_1-42_ ratios of 10/1 (green curves), 1/1 (orange curves) and 0.1/1 (blue curve, only for **β-Hsp90**). The control curves are represented in red lines and buffer in grey.

**Extended data Fig. 9:**
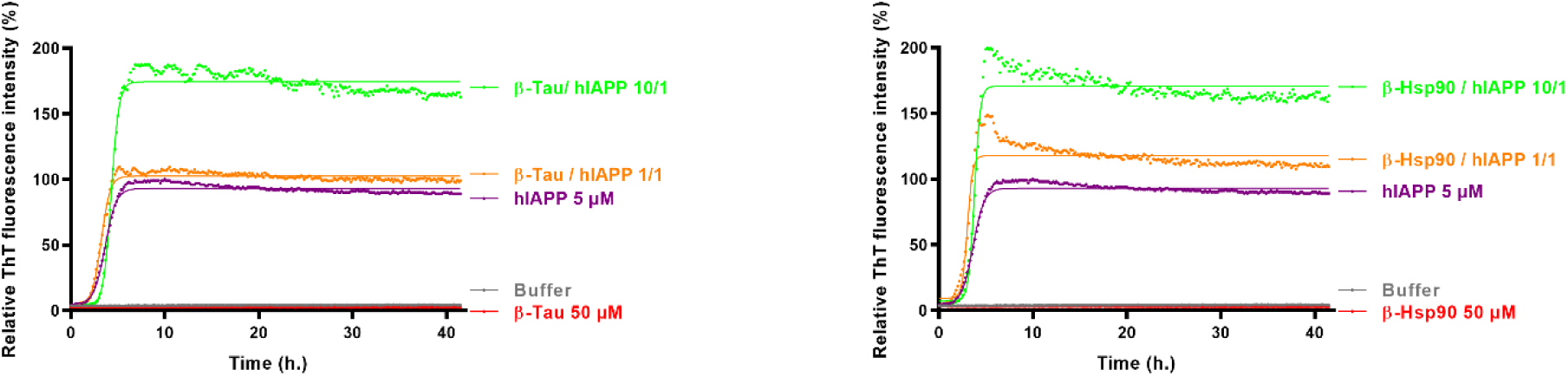
Representative curves of ThT fluorescence assays over time showing hIAPP fibrillization (5 µM) in the absence (purple curve) and in the presence of **β-Tau** and **β-Hsp90** at compound/Aβ_1-42_ ratios of 10/1 (green curves) and 1/1 (orange curves). The control curves are represented in red lines and buffer in grey.

**Extended data Fig. 10:**
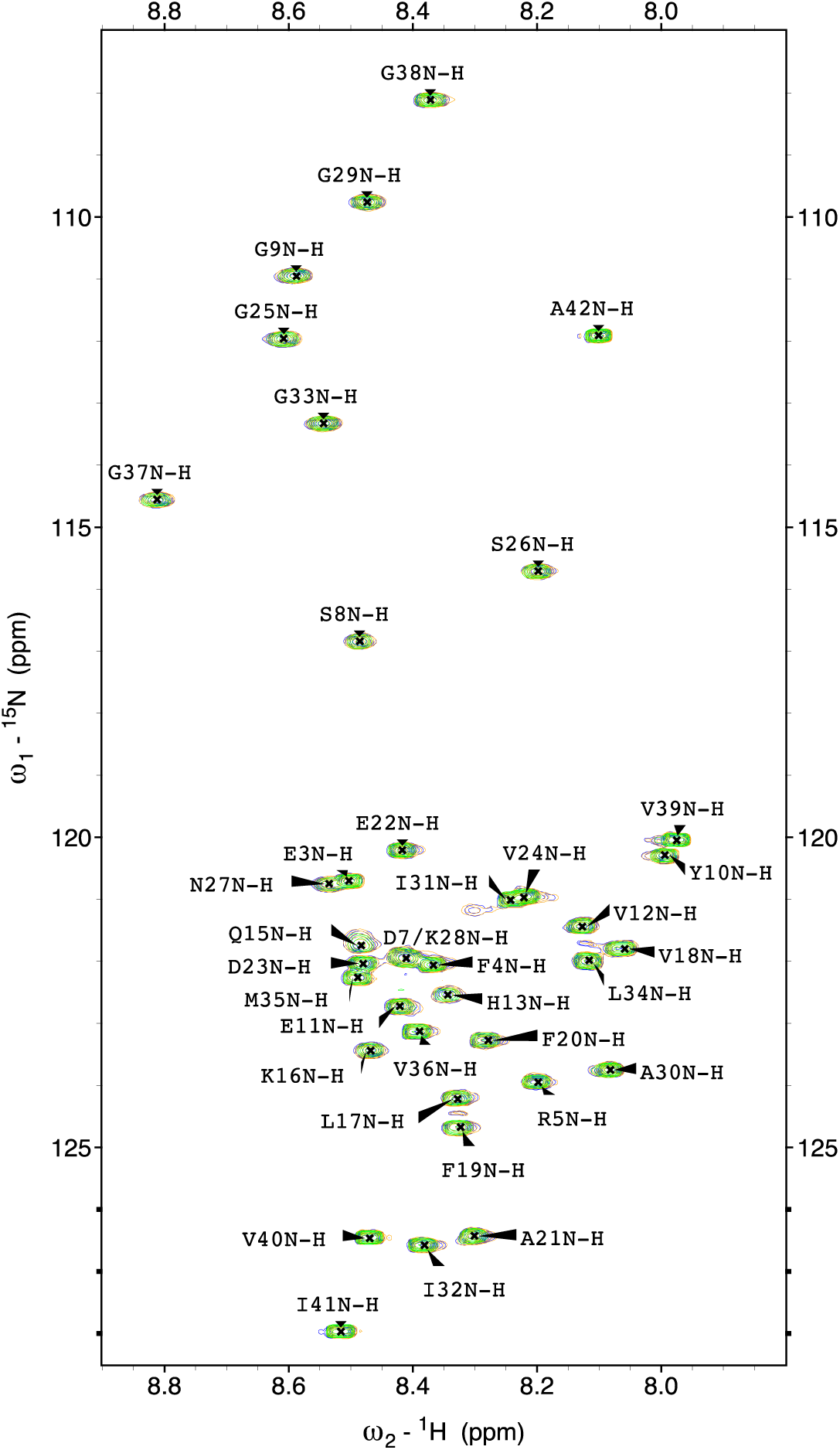
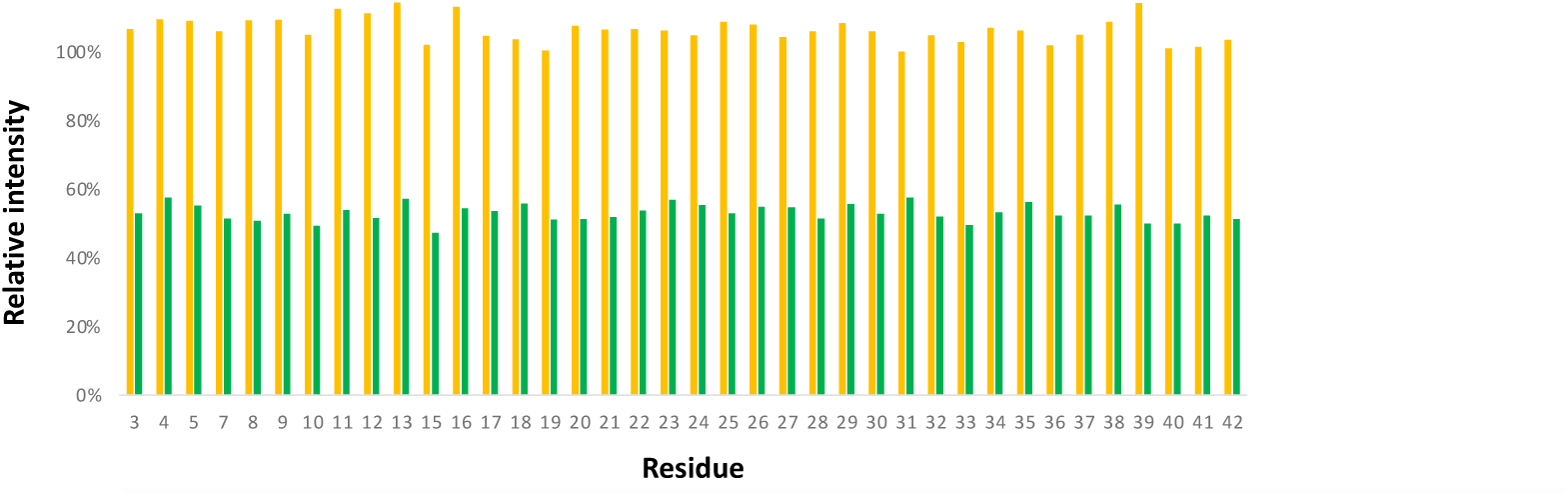
2D ^1^H-^15^N HSQC spectra of ^15^N-Aβ_1-42_ (50 μM) in the absence (blue contours) and in the presence of β-Hsp90 (50 μM, orange contours and 500 μM, green contours). No significant ^1^H, ^15^N chemical shifts perturbation could be detected. The HSQC relative peak intensity at β-Hsp90/Aβ_1-42_ ratios of 1/1 (orange) and 10/1 (green) is calculated for each residue with respect to the reference spectrum in the absence of β-Hsp90.

**Extended data Table 1:** Relative fluorescence of compounds **β-Tau**, **β-Hsp90**, **H1**, **S4**, **S6**, **S7**, **S4^short^**, **S7^short^**, **S4^short^** plus **S7^short^** related to the maximum fluorescence obtained for the ThT-fluorescence control curve of Ac-PHF6* at 5/1, 1/1 and 0.1/1 compound/Ac-PHF6* ratios.

| Compound | Compound/ Ac-PHF6* Ratio<sup>[a]</sup> | Relative Fluorescence (%)<sup>[b]</sup> |
| --- | --- | --- |
| β-Tau | (5/1) | 9±1% |
| | (1/1) | 19±4% |
| | (0.1/1) | 13±2% |
| β-Hsp90 | (5/1) | 28±24% |
| | (1/1) | 25±24% |
| | (0.1/1) | 62±15% |
| H1<sup>[c]</sup> | (5/1) | 243±4% |
| | (1/1) | 133±5% |
| | (0.1/1) | 102±5% |
| S4 | (5/1) | 4±1% |
| | (1/1) | 125±4% |
| | (0.1/1) | 102±4% |
| S6 | (5/1) | 3±1% |
| | (1/1) | 114±4% |
| | (0.1/1) | 105±2% |
| S7 | (5/1) | 2±1% |
| | (1/1) | 3±1% |
| | (0.1/1) | 100±3% |
| S4<sup>short</sup> | (5/1) | 1±1% |
| | (1/1) | 85±1% |
| | (0.1/1) | 110±4% |
| S7<sup>short</sup> | (5/1) | 5±1% |
| | (1/1) | 155±2% |
| | (0.1/1) | 108±2% |
| S4<sup>short</sup> + S7<sup>short</sup> | (2.5+2.5/1) | 3±1% |
| | (0.5+0.5/1) | 56±5% |
| | (0.05+0.05/1) | 87±3% |

Parameters are expressed as mean ± SE, n=3. <sup>[a]</sup> Compounds were dissolved in water. The concentration of Ac-PHF6\* was 25 μM. <sup>[b]</sup> The fluorescence for Ac-PHF6\* control was set at 100 % for the maximal fluorescence value and the relative fluorescence was calculated compared to this value. <sup>[c]</sup> Compound H1 has a strong tendency to self-aggregate at 125 μM.

**Extended data Table 2:**
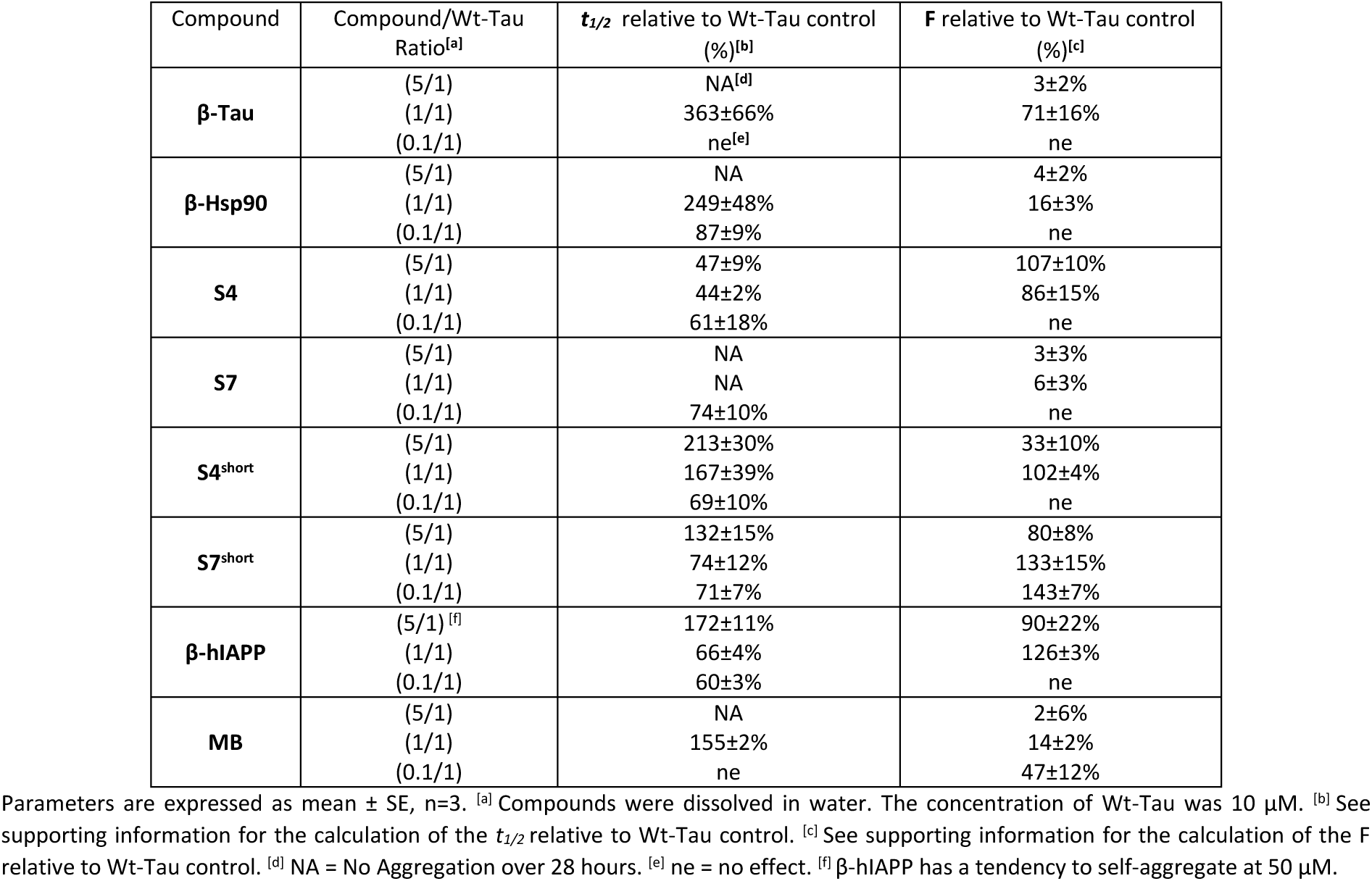
Effects of compounds **β-Tau**, **β-Hsp90**, **S4**, **S7**, **S4^short^**, **S7^short^**, **MB and β-hIAPP** on WT-Tau fibrillization assessed by ThT-fluorescence spectroscopy at 5/1, 1/1 and 0.1/1 ratios of compound/Wt-Tau.

**Extended data Table 3:** *In vitro* stability of peptides **β-Tau**, **β-Hsp90**, **S7**, **S4^short^** and **S7^sho^**^rt^ in Pronase mixture at 37 °C. Percentage of intact peptides at 0, 10, 30, 60, 120 minutes. NP = no pronase at t=0

| Compound | NP | Time (min) |  |  |  |  |
| --- | --- | --- | --- | --- | --- | --- |
|  |  | 0 | 10 | 30 | 60 | 120 |
| β-Tau | 100 | 100 | 87% | 84% | 62% | 30% |
| β-Hsp90 | 100 | 100 | 100% | 100% | 100% | 79% |
| S7 | 100 | 100 | 24% | 0% | 0% | 0% |
| S7<sup>short</sup> | 100 | 100 | 83% | 46% | 21% | 10% |
| S4<sup>short</sup> | 100 | 100 | 91% | 89% | 83% | 71% |

**Extended data Table 4:** Effects of compounds **β-Tau** and **β-Hsp90** on Tau-Δk280 fibrillization assessed by ThT-fluorescence spectroscopy at 5/1, 1/1 and 0.1/1 ratios of compound/Tau-Δk280.

| Compound | Compound/Wt-Tau Ratio<sup>[a]</sup> | $t_{1/2}$ relative to Tau-Δk280 control (%)<sup>[b]</sup> | F relative to Tau-Δk280 control (%)<sup>[c]</sup> |
| --- | --- | --- | --- |
| β-Tau | (5/1) | NA<sup>[d]</sup> | 3±7% |
| | (1/1) | 279±54% | 25±7% |
| | (0.1/1) | 121±24% | 108±38% |
| β-Hsp90 | (5/1) | NA | 7±1% |
| | (1/1) | NA | 4±1% |
| | (0.1/1) | 85±26% | 34±6% |

Parameters are expressed as mean ± SE, n=3. <sup>[a]</sup> Compounds were dissolved in water. The concentration of Tau-Δk280 was 10 μM. <sup>[b]</sup> See supporting information for the calculation of the $t_{1/2}$ relative to Tau-Δk280 control. <sup>[c]</sup> See supporting information for the calculation of the F relative to Tau-Δk280 control. <sup>[d]</sup> NA = No Aggregation.

**Extended data Table 5:** Effects of compounds **β-Tau**, **β-Hsp90**, **S4**, **S7**, **S4^short^** and **S7^short^** on Aβ_1-42_ fibrillization assessed by ThT-fluorescence spectroscopy at 10/1 and 1/1 ratios of compound/Aβ_1-42_.

| Compound | Compound/ A$\beta_{1-42}$ Ratio<sup>[a]</sup> | $t_{1/2}$ relative to A$\beta_{1-42}$ control (%)<sup>[b]</sup> | F relative to A$\beta_{1-42}$ control (%)<sup>[c]</sup> |
| --- | --- | --- | --- |
| β-Tau | (10/1) | 221±32% | 28±10% |
| | (1/1) | 127±15% | 104±4% |
| β-Hsp90 | (10/1) | NA<sup>[d]</sup> | 3±2% |
| | (1/1) | 308±25% | 10±4% |
| | (0.1/1) <sup>[e]</sup> | 194±36% | 70±16% |
| S4 | (10/1) | 180±22% | 120±7% |
| | (1/1) | 104±3% | 61±13% |
| S7 | (10/1) | 106±3% | 96±3% |
| | (1/1) | 106±10% | 98±28% |
| S4<sup>short</sup> | (10/1) | 232±17% | 36±4% |
| | (1/1) | 114±15% | 95±12% |
| S7<sup>short</sup> | (10/1) | 131±11% | 89±12% |
| | (1/1) | 84±8% | 95±14% |

Parameters are expressed as mean ± SE, n=2. <sup>[a]</sup> The concentration of A $\beta_{1-42}$ was 10 μM. <sup>[b]</sup> See supporting information for the calculation of the $t_{1/2}$ relative to A $\beta_{1-42}$ control. <sup>[c]</sup> See supporting information for the calculation of the F relative to A $\beta_{1-42}$ control. <sup>[d]</sup> NA = No Aggregation over 40 hours. <sup>[e]</sup> Due to the strong activity of β-Hsp90 at 1/1 ratio, it was tested at a lower 0.1/1 ratio.

