## Supplementary material for "Synthetic chaperone based on Hsp90-Tau interaction inhibits pathological Tau aggregation and rescues physiological Tau-Microtubule interaction": Synthesis of compounds and CD spectra

#### Table of contents


**General procedures for the syntheses** **3**

**Compounds and purities** **7**

**Circular Dichroism spectra** **25**

### General procedures for the synthesis

Usual solvents were purchased from commercial sources. Thin-layer chromatography (TLC) analyses were performed on silica gel 60 F250 (0.26 mm thickness) plates. The plates were visualized with UV light (λ = 254 nm) alternatively stained with a 4 % solution of phosphomolybdic acid or ninhydrin in EtOH.

NMR spectra were recorded on an ultra-field Bruker 300 ($^{1}$H, 300 MHz, $^{13}$C, 75 MHz) or on a Bruker AVANCE 400 ($^{1}$H, 400 MHz, $^{13}$C, 101 MHz, $^{19}$F 376 MHz). Chemical shifts δ are in ppm with the solvent resonance as the internal standard ($^{1}$H NMR, $CDCl_3$: δ = 7.26 ppm, $CD_3OD$: δ = 3.31 ppm, $CD_3CN$: δ = 1.93 ppm; $^{13}$C NMR, $CDCl_3$: δ = 77.16 ppm, $CD_3OD$: δ = 49.00 ppm; $CD_3CN$: δ = 1.3 ppm), and the following abbreviations are used: singlet (s), doublet (d), doublet of doublet (dd), triplet (t), quintuplet (qt), multiplet (m), broad multiplet (brm), and broad singlet (brs), broad doublet (brd). Mass spectra were obtained using a Bruker Esquire electrospray ionization apparatus. HRMS were obtained using a TOF LCT Premier apparatus (Waters) with an electrospray ionization source. The purity of compounds was determined by HPLC-MS on Agilent 1260 Infinity.

Column: ATLANTIS T3 column (C18, 2.1 x 150mm-3µm), mobile phase: ACN/$H_2O$ + 0.1% TFA (gradient 1-30% in 15 or 20 min).

Preparative HPLC were performed on Agilent 1260 Infinity II. Column: Pursuit (C18 10 x 250µm-5µm), mobile phase: ACN/$H_2O$ + 0.1% formic acid (FA) and on Waters XBridge BEH300 (C18, 2.1 x 150mm-5µM).

UPLC-MS analyses were performed on a Waters Acquity UPLC apparatus equipped with a Luna Omega PSC18 Column (1.5 µM, 2.1 x 50 mm) coupled to a single quadrupole EDI-MS (Mictomass ZQ). ESI mass spectra were recorded on an LCQESI MS on a LCQ Advantage spectrometer from Thermo Finningan and a LCQ Fleet spectrometer from Thermo Scientific.

The preparation of the piperidine-pyrrolidine β-turn scaffold was done accordingly to our previous publications[1,2]

#### A. General procedure for the Synthesis on Solid phase peptide Synthesis Strategy (SPSS) for compounds β-Tau and β-Hsp90 .

All the reactions involved were agitated in plastic syringe tubes equipped with filters on an automated shaker on Rink Amide resin (0.2 mmol scale, loading 0.327 mmol/g, 600 mg). The coupling yields were monitored with the Fmoc-test procedure (reported here below).

Removal of Fmoc group was performed in 20% (v/v) piperidine/DMF for 20 min twice. Capping steps were performed by treating the resin with the mixture of acetic anhydride (0.25 M) and NMM (0.25 M) in DMF solution for 20 min. After each reaction, the resin was washed with DMF (3 × 10mL), MeOH (3 × 10mL) and DCM (3 × 10mL) successively.

Rink Amide resin (600 mg, 0.2 mmol/g) was swelled in DMF for 1h before using. Natural amino acids and scaffold were coupled using different coupling reactive accordingly to their position:

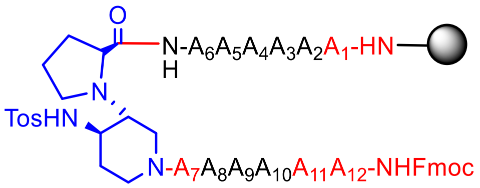


- **Red positions**: The resin was suspended in DMF (4 mL) and collidine (17 eq). Fmoc-AA-OH (5 eq) DIC (5 eq) and Oxyma pure (5 eq) were solubilized in DMF/DCM 33% (v/v) (3 mL) and let under magnetic stirring for 7 minutes. Afterward, the solution containing the amino acid was poured into the reactor containing the resin and the mixture shaked for 16h at room temperature. (Positions 1, 7, 11 and 12)

- **Black position**: Fmoc-AA-OH ( 5 eq) and HCTU (5 eq) were solubilized in 5 ml of DMF/NMM 20% (V/V). The prepared solution is added to the resin and shaked for 20 minutes. Afterward, the solvent was removed and the resin washed with DMF (1 x 5 mL) and the coupling procedure repeated. (Positions from 2 to 6 and from 8 to 10)

- **Blue position**: The resin was suspended in DMF (4 mL) and collidine (17 eq). **8** (1.5 eq) DIC (1.5 eq) and OXYMA (1.5 eq) were solubilized in DMF/DCM 33% (3 mL) and let under magnetic stirring for 7 minutes. Afterward, the solution containing the peptidomimetic was poured into the reactor containing the resin and the mixture shaked for 16h at room temperature.

At the end of the synthesis the peptides were cleaved from the resin shaking for 2 hours in the presence of 5 ml of an acidic solution containing trifluoracetic acid/$H_2O$/TIPS/Phenol/Thioanisole; 87.5%/5%/2.5%/2.5%/2.5%. The liquid was poured in dry cold $Et_2O$ (40 mL) in ice bath by filtering it over a cotton pad. The peptides precipitated and isolated by centrifugation at 6000 rounds/min for 7 min. The pad was resuspended with ether and centrifuged again to remove the remaining TFA. The hairpin mimics were lyophilized and purified over reverse phase RP-HPLC.

#### Fmoc loading Test phase (Fmoc-test)

From 5 to 10 mg of dry resin was shaken in DMF/Piperidine 20% solution (1 mL) for 30 minutes. The resin was sedimented by centrifugation and 1 mL of the supernatant was added to 9 mL of DMF and mixed well. This final solution was used for the UV analysis measuring the absorbance at 301 nm versus a DMF blank (triplicate analysis).

For the quantification of resin loading, we used the general formula: $L = (A_{301} x\ V\ x\ d)/(E_c\ x\ w\ x\ M)$. Where: L = Resin loading; $A_{301}$ = UV Absorbance at 301 nm; V = Volume of the cleavage solution = 1 mL; d = Dilution = 10; $E_c$ = Extinction coefficient = 7800 mL/mmol*cm; w = Width of the cuvette = 1 cm; M = Weight of the resin sample in g. [Ref: aaptec https://www.peptide.com/custdocs/1198.pdf] Substituting the extinction coefficient, volume, dilution and cell width into the general formula results in this formula which can be used to calculate the resin loading.

$L = (100\ x\ A_{301})/(7.8\ x\ M(mg))$ in units of mmols/gram.

#### B. Synthesis of peptides H1, S4, S6 and S7

Peptides were synthesized using microwave assisted solid phase peptide synthesis performed on a Liberty Blue Microwave Automated Peptide Synthesizer (CEM Corporation, Matthews, NC, USA), following the standard protocols for Fmoc/tBu strategy (0.1 mmol scale, 300 mg, 0.327 mmol/g). Fmoc-deprotection cycles were respectively of 15 s (75 °C, 155 W) and 30 s (90 °C, 30 W). Couplings of Arg residues were performed in 1500 s (25 °C, 0 W) and 300 s (75 °C, 30 W). Couplings of His were performed in 240 s (50 °C, 35 W). Couplings of other residues are performed in 15 s (90 °C, 170 W) and 110 s (90 °C, 30 W). Peptide cleavage (3 h at room temperature) from the resin and deprotection of the amino acid side chains were performed by using the reagent K[14] (trifluoracetic acid/phenol/water/thioanisol/1.2-ethanedithiol; 82.5/5/5/5/2.5) for 180 min. The liquid was poured in dry cold $Et_2O$ (40 mL) in ice bath by filtering it over a cotton pad. The peptides precipitated and isolated by centrifugation at 6000 rounds/min for 7 min. The pad was resuspended with ether and centrifuged again to remove the remaining TFA. The peptides were lyophilized and purified over RP-HPLC if necessary.

#### C. Synthesis of peptides S4$^{short}$, S7$^{short}$ and Ac-PHF6*-$NH_2$ via manual solid phase peptide synthesis

All the reaction involved were agitated in plastic syringe tubes equipped with filters on an automated shaker, on Rink Amide resin (0.2 mmol scale, loading 0.327 mmol/g, 600 mg).

Removal of Fmoc group was performed in 20% (v/v) piperidine/DMF for 20 min twice. Capping steps were performed by treating the resin with the mixture of acetic anhydride (0.25 M) and NMM (0.25 M) in DMF solution for 20 min. After each reaction, the resin was washed with DMF (3 × 10mL), MeOH (3 × 10mL) and DCM (3 × 10mL) successively.

Rink Amide resin (600 mg, 0.2 mmol/g) was swelled by DMF for 1h before using. The loading was performed by suspending the resin with DMF (4 mL) and collidine (17 eq). Fmoc-AA-OH (5 eq) DIC (5 eq) and Oxyma pure (5 eq) were solubilized in DMF/DCM 33% (v/v) (3 mL) and let under magnetic stirring for 7 minutes. Afterward, the solution containing the amino acid was poured into the reactor containing the resin and the mixture shaked for 16h at room temperature. Every following coupling using Fmoc-protected natural amino acid was carried out twice to get satisfactory yields using Fmoc-AA-OH/HCTU (2.5/2.5 eq) in NMM/DMF (20%V/V, 4 ml).

Peptides were cleaved from the resin by shaking for 2 hours with an acidic mixture containing TFA/water/TIPS/Thioanisole; 95%/2.5%/1.25%/1.25%) and repeated the same procedure explained in General procedure **A**.

**β-Tau**

*(S)-N[1]-((2S,3S)-1-(((2S,3S)-1-(((S)-4-amino-1-(((S)-6-amino-1-((3R,4R)-3-((S)-2-(((3S,6S,9S,12S,15S,18S)-22-amino-6-(3-amino-3-oxopropyl)-9-((S)-sec-butyl)-18-carbamoyl-15-(4-hydroxybenzyl)-12-isopropyl-2-methyl-4,7,10,13,16-pentaoxo-5,8,11,14,17-pentaazadocosan-3-yl)carbamoyl)pyrrolidin-1-yl)-4-((4-methylphenyl)sulfonamido)piperidin-1-yl)-1-oxohexan-2-yl)amino)-1,4-dioxobutan-2-yl)amino)-3-methyl-1-oxopentan-2-yl)amino)-3-methyl-1-oxopentan-2-yl)-2-((S)-2-amino-3-methylbutanamido)pentanediamide*

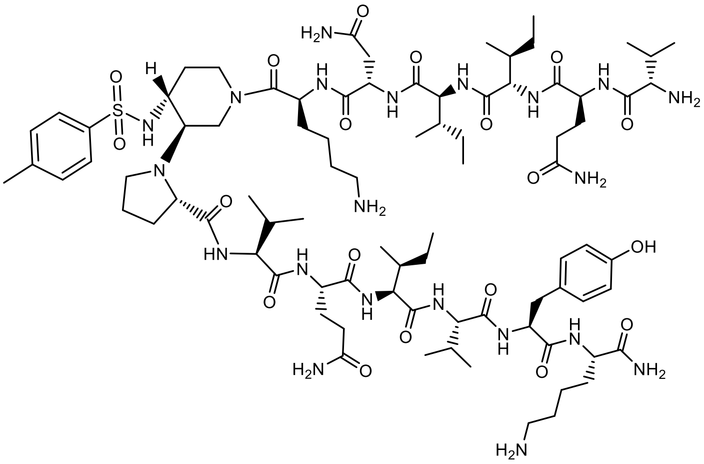

Peptidomimetic **β-Tau** was synthesized accordingly to **General procedure A**.

**β-Tau** was purified by semi-preparative RP-HPLC (Gradients of 5-40 % ACN in $H_2O$ containing 0.1% TFA in 20 min, retention time = 10.86 min, yield isolation: 45%).

**Molecular weight**: 1792.0417 g/mol

**HRMS**: Calcd. for $[C_{85}H_{141}N_{21}O_{19}S + H]^+$: m/z 1793.0506 found 1793.0497 $[M + H]^+$ and Calcd. for $[C_{85}H_{141}N_{21}O_{19}S + Na]^+$: m/z 1815.0326 found 1815.0303 $[M + Na]^+$

**UPLC purity**: POROSHELL 120 column (C18, 2.1 x 50 mm-1.9 µm); (Gradients of 5-100% ACN in $H_2O$ containing 0.1% FA in 10 min); Rt = 2.50 min, 100 %.

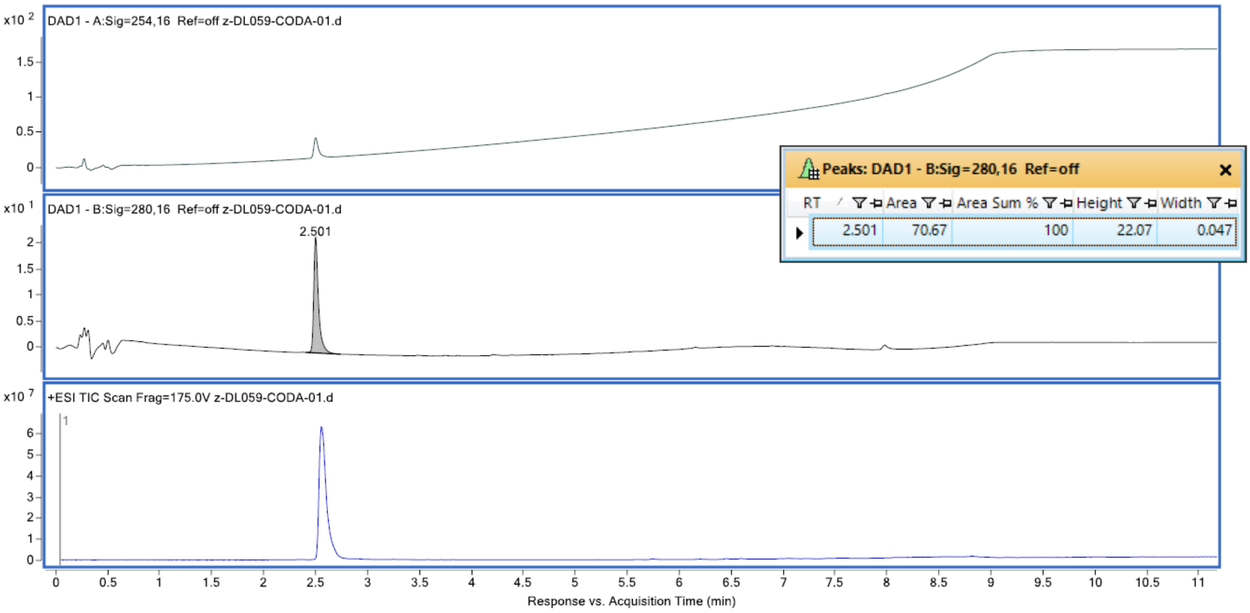

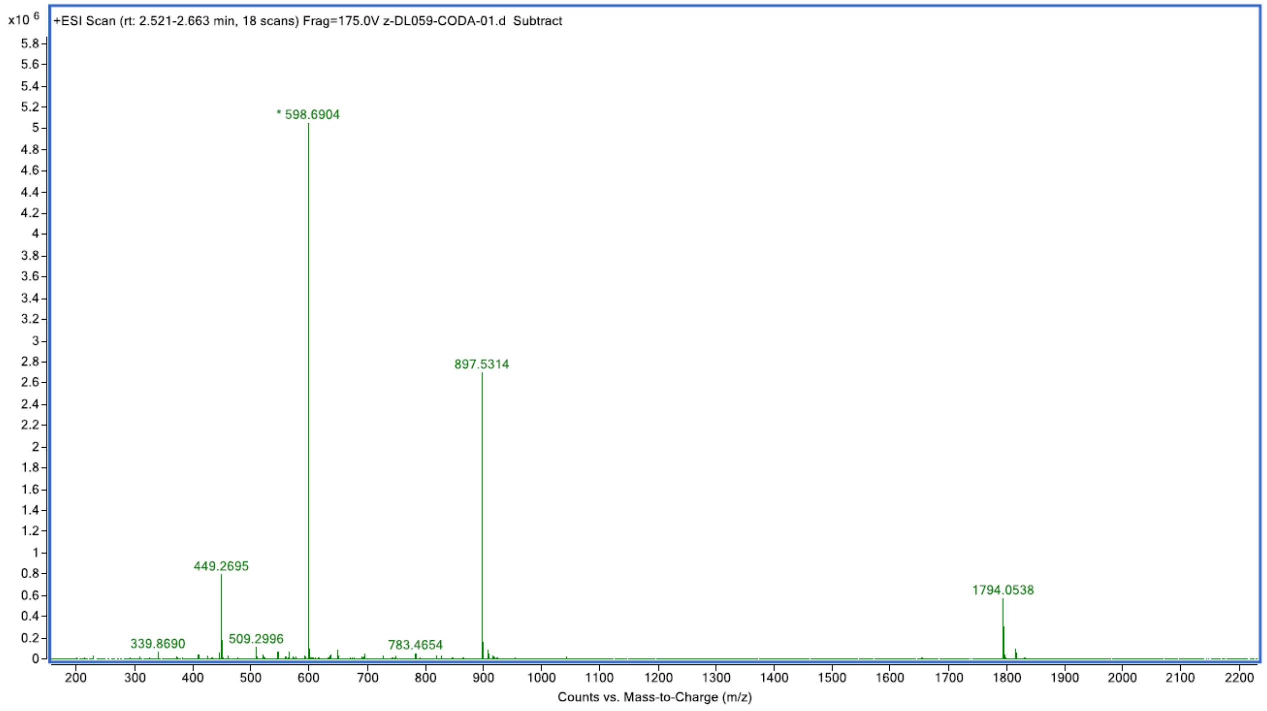


| Fragmentor Voltage | Collision Energy | Ionization Mode |
|---|---|---|
| 175 | 0 | ESI |

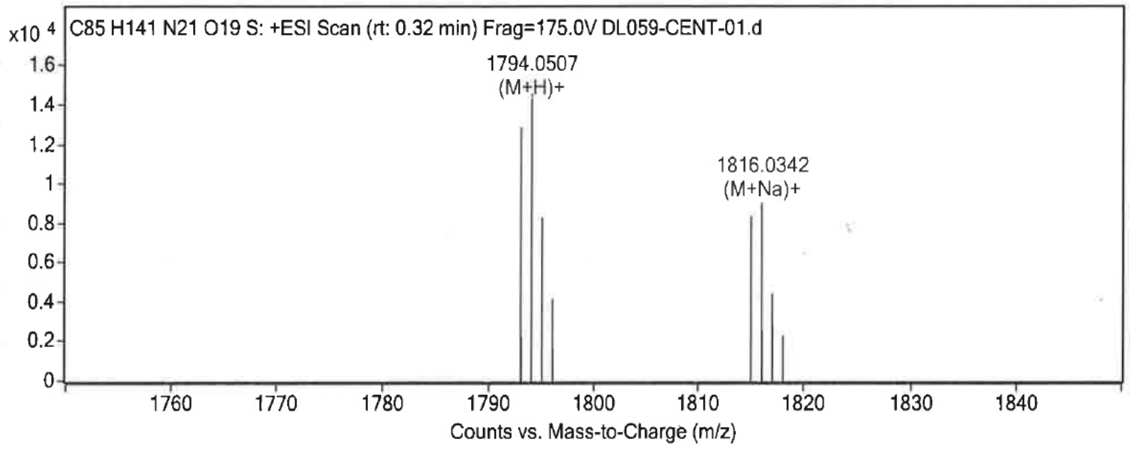


**Formula Calculator Results**

| Best | Generated Molecular Formula | Ion m/z | Generated Ion Formula | Calculated Molec Mass | Theorical Molec Mass | Rel Diff (ppm) | Diff (mDa) | Mass Match Probability |
|---|---|---|---|---|---|---|---|---|
| VRAI | C85 H141 N21 O19 S | 1793.0497 | C85 H142 N21 O19 S | 1792.0417 | 1792.0433 | -0.93 | -1.66 | 99.47 |
| VRAI | C85 H141 N21 O19 S | 1815.0303 | C85 H141 N21 Na O19 S | 1792.0402 | 1792.0433 | -1.75 | -3.14 | 98.17 |

**Formula Calculator Results**

| Generated Molec Formula | Ion Species | Ion Formula | Best | Ion m/z | Calculated Molec Mass | Theorical Molec Mass | Rel Diff (ppm) | Diff (mDa) | Mass Match Probability | Match Score |
|---|---|---|---|---|---|---|---|---|---|---|
| C88 H137 N21 O19 S | (M+H)+ | C88 H138 N21 O19 S | VRAI | 1825.0165 | 1824.0091 | 1824.0120 | -1.63 | -2.97 | 95.68 | 94.28 |
| C88 H137 N21 O19 S | (M+Na)+ | C88 H137 N21 Na O19 S | VRAI | 1846.9985 | 1824.0093 | 1824.0120 | -1.49 | -2.73 | 96.40 | 98.06 |

**β-Hsp90**

*(S)-1-((3R,4R)-1-(L-lysyl-L-valyl-L-valyl-L-valyl-L-isoleucyl-L-allothreonyl)-4-((4-methylphenyl)sulfonamido)piperidin-3-yl)-N-((2S,3S)-1-(((S)-6-amino-1-(((S)-1-(((2S,3S)-1-(((S)-1-(((S)-1-amino-3-(1H-imidazol-4-yl)-1-oxopropan-2-yl)amino)-4-methyl-1-oxopentan-2-yl)amino)-3-methyl-1-oxopentan-2-yl)amino)-3-methyl-1-oxobutan-2-yl)amino)-1-oxohexan-2-yl)amino)-3-hydroxy-1-oxobutan-2-yl)pyrrolidine-2-carboxamide*

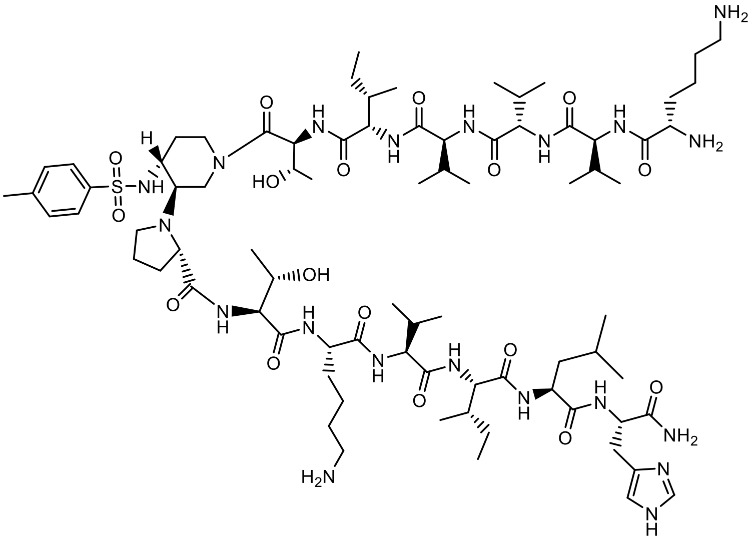

Peptidomimetic **β-Hsp90** was synthesized accordingly to **General procedure A**.

**β-Hsp90** was purified by semi-preparative RP-HPLC (Gradients of 5-47.2 % ACN in $H_2O$ containing 0.1% TFA in 13 min, retention time = 10.34 min, yield isolation: 53%).

**Molecular weight**: 1697.0419 g/mol

**HRMS**: Calcd. for $[C_{81}H_{140}N_{20}O_{17}S + H]^+$: m/z 1698.0499 found 1698.0491 $[M + H]^+$ and Calcd. for $[C_{81}H_{140}N_{20}O_{17}S + Na]^+$: m/z 1710.0318 found 1720.0310 $[M + Na]^+$

**HPLC purity**: XSELECT column (C18, 2.1 x 75mm-2.5µm); (Gradients of 5-100% ACN in $H_2O$ containing 0.1% FA in 15 min); Rt = 5.018 min, 100 %.

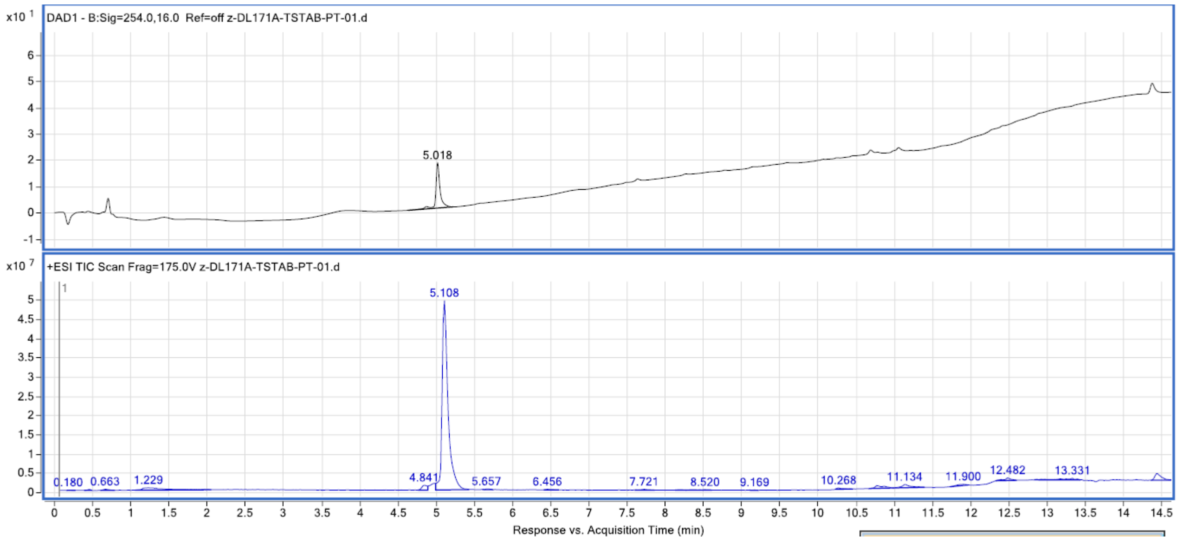

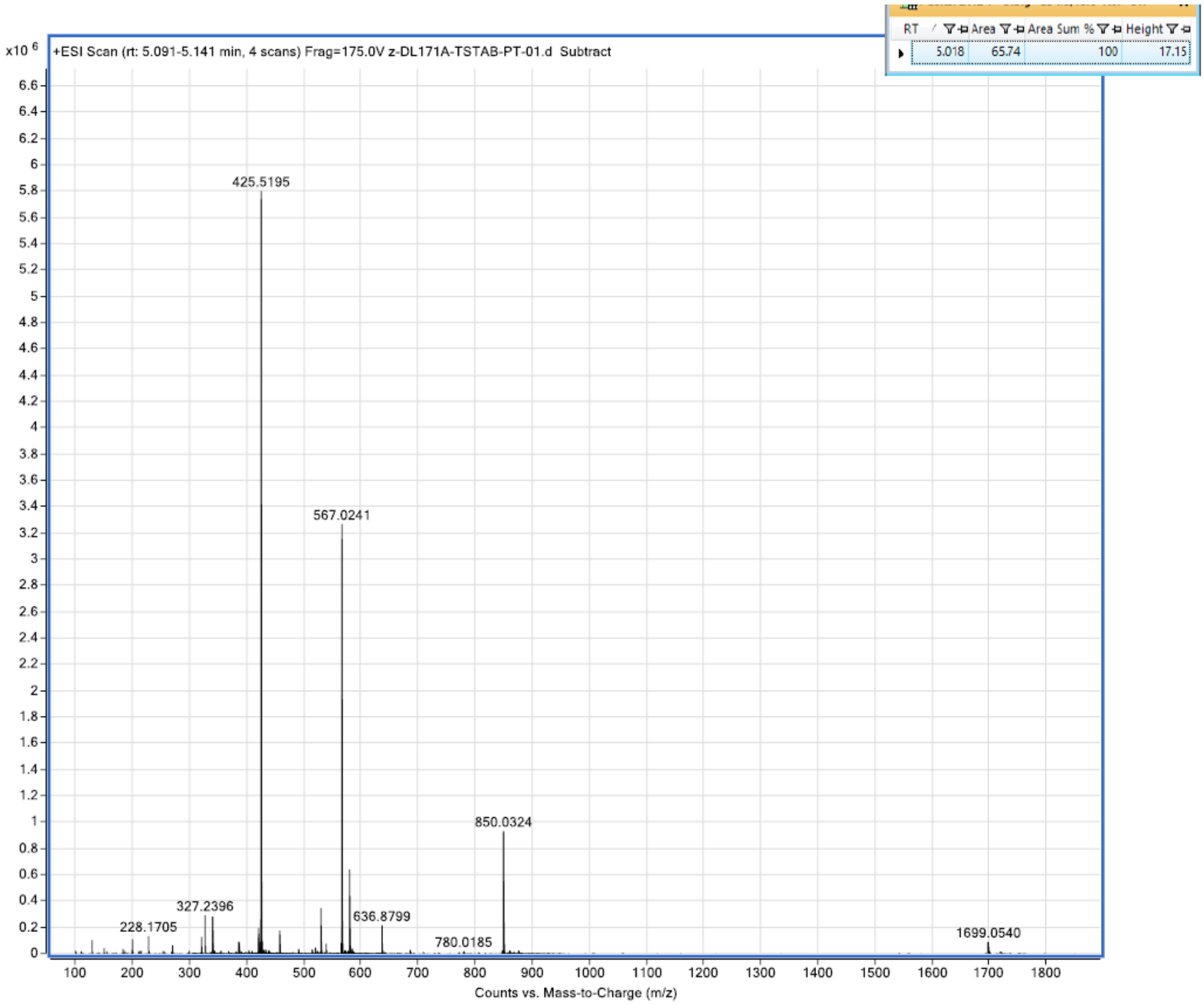


| Spectrum Source | Fragmentor Voltage | Collision Energy | Ionization Mode |
|---|---|---|---|
| Peak (1) in "+ TIC Scan" | 175 | 0 | ESI |

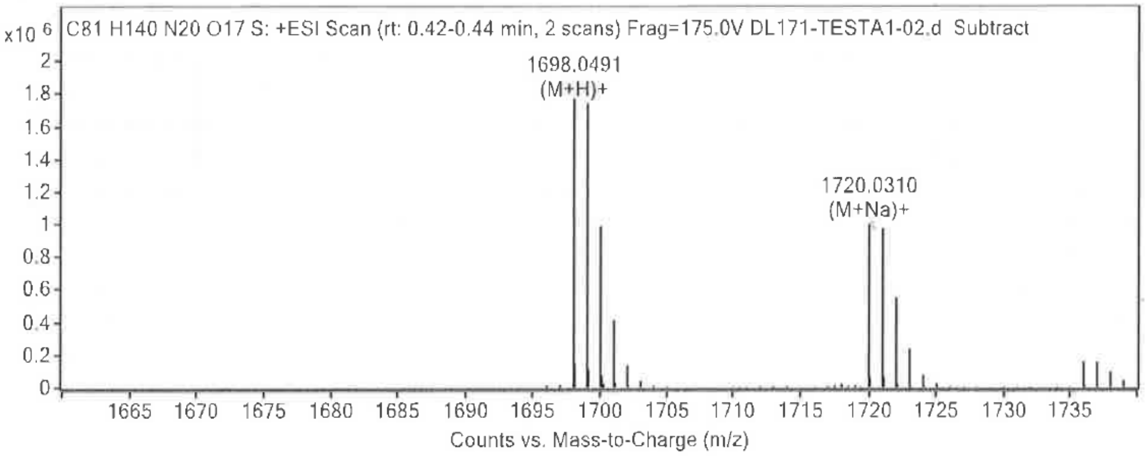


**Formula Calculator Results**

| Best | Generated Molecular Formula | Ion m/z | Generated Ion Formula | Calculated Molec Mass | Theorical Molec Mass | Rel Diff (ppm) | Diff (mDa) | Mass Match Probability |
|---|---|---|---|---|---|---|---|---|
| VRAI | C81 H140 N20 O17 S | 1698.0491 | C81 H141 N20 O17 S | 1697.0419 | 1697.0426 | -0.44 | -0.75 | 99.90 |
| VRAI | C81 H140 N20 O17 S | 1720.0310 | C81 H140 N20 Na O17 S | 1697.0419 | 1697.0426 | -0.39 | -0.67 | 99.92 |

**Compound H1**

*(2S,5S,8S,11S,14S,17S,20S,23S,26S,29S,32S,35S,38S,41S)-1-amino-11-(2-amino-2-oxoethyl)-32-(3-amino-3-oxopropyl)-5-benzyl-14,17,29,38-tetra((S)-sec-butyl)-41-((S)-2-((S)-2,5-diamino-5-oxopentanamido)propanamido)-2-(4-hydroxybenzyl)-8-((S)-1-hydroxyethyl)-23-(hydroxymethyl)-20-isobutyl-35-methyl-26-(2-(methylthio)ethyl)-1,4,7,10,13,16,19,22,25,28,31,34,37,40-tetradecaoxo-*3,6,9,12,15,18,21,24,27,30,33,36,39-tridecaazatetratetracontan-44-oic acid

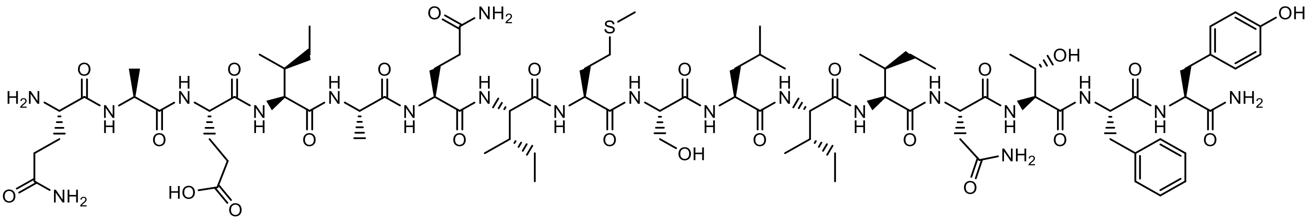

Peptide **H1** was synthesized accordingly to **General procedure B**.

The peptide was purified by semi-preparative RP-HPLC (Gradients of 5-100 % ACN in $H_2O$ containing 0.1% TFA in 30 min, retention time = 14.97 min, yield isolation: 4%).

**Molecular weight**: 1852.9757 g/mol

**HRMS**: Calcd. for $[C_{85}H_{136}N_{20}O_{24}S + Na]^+$: m/z 1875.9645 found 1875.9670 $[M + Na]^+$

**HPLC purity**: SYNERGI POLAR (C18, 2 x 100mm-2.5µm); (Gradients of 5-100% ACN in $H_2O$ containing 0.1% FA in 20 min); Rt = 9.37 min, 62 %.

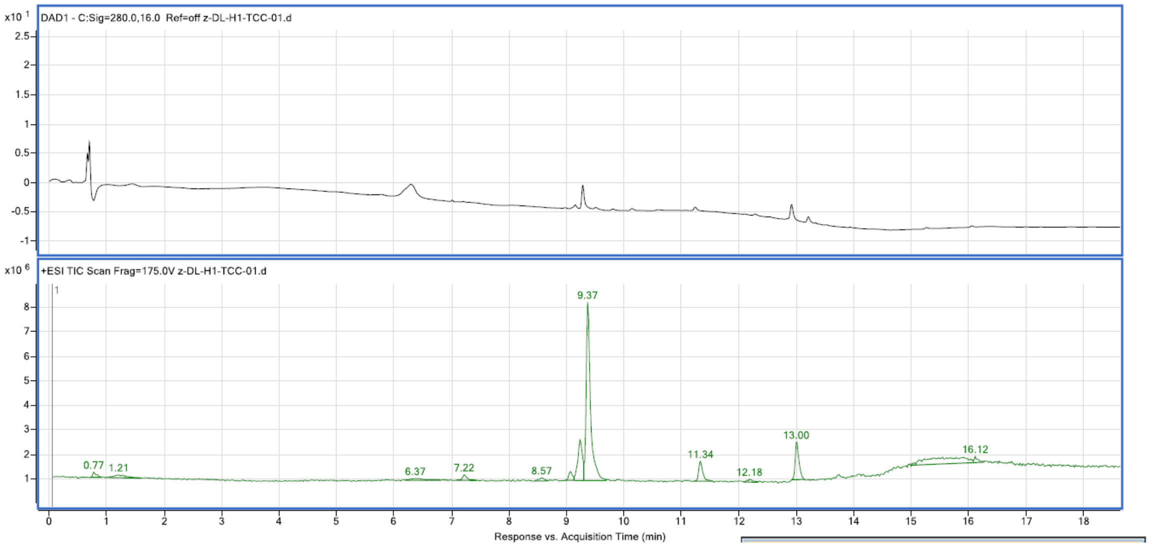

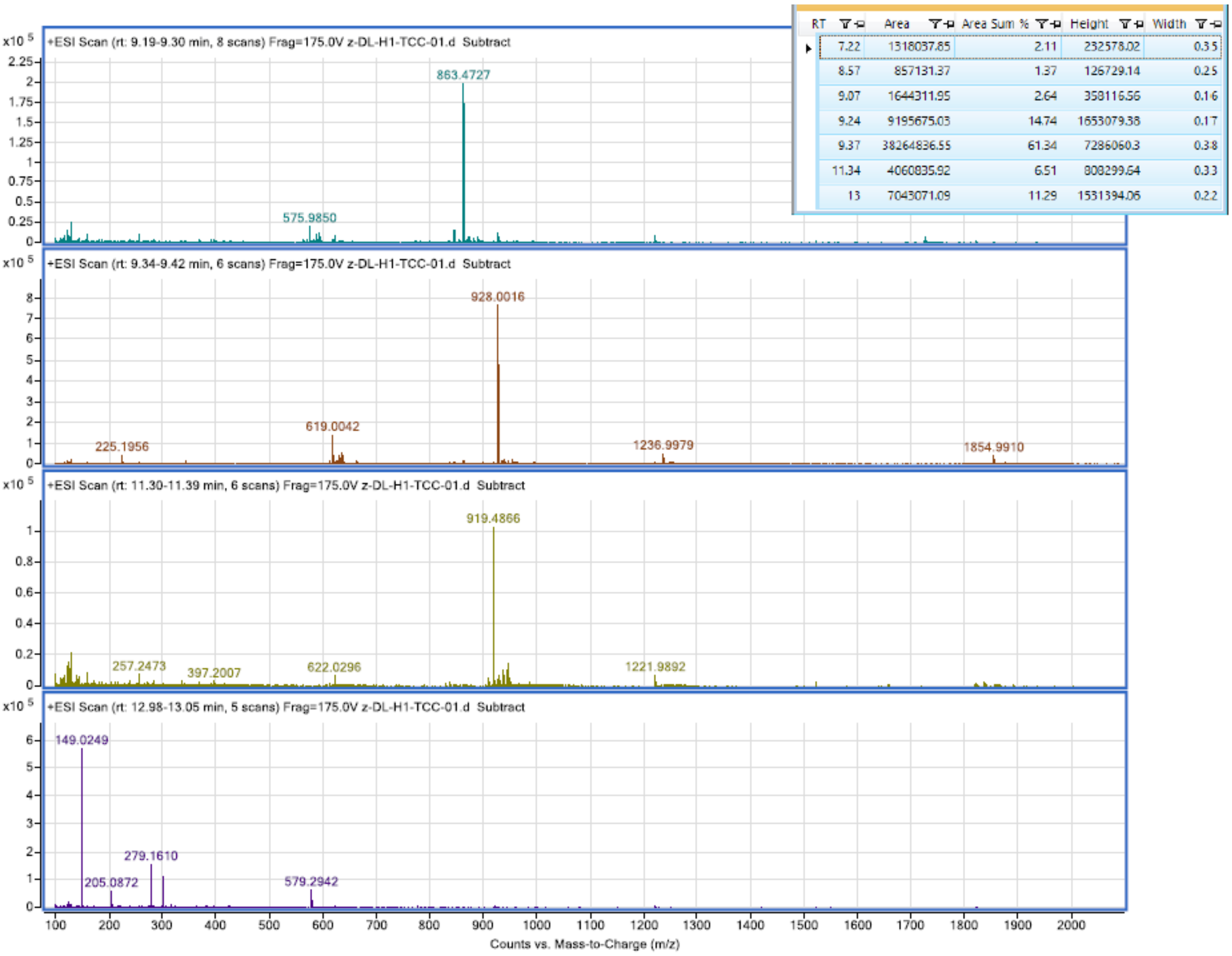


| RT | Area | Area Sum % | Height | Width |
|---|---|---|---|---|
| 7.22 | 1318037.85 | 2.11 | 232578.02 | 0.35 |
| 8.57 | 857131.37 | 1.37 | 126729.14 | 0.25 |
| 9.07 | 1644311.95 | 2.64 | 358116.56 | 0.16 |
| 9.24 | 9195675.03 | 14.74 | 1653079.38 | 0.17 |
| 9.37 | 38264836.55 | 61.34 | 7286060.3 | 0.38 |
| 11.34 | 4060835.92 | 6.51 | 808299.64 | 0.33 |
| 13 | 7043071.09 | 11.29 | 1531394.06 | 0.22 |

**Formula Calculator Results**

| Best | Generated Molecular Formula | Ion m/z | Generated Ion Formula | Calculated Molec Mass | Theorical Molec Mass | Rel Diff (ppm) | Diff (mDa) | Mass Match Probability |
|---|---|---|---|---|---|---|---|---|
| VRAI | C85 H136 N20 O24 S | 927.4964 | C85 H138 N20 O24 S | 1852.9775 | 1852.9757 | 0.98 | 1.82 | 99.69 |
| VRAI | C85 H136 N20 O24 S | 949.4771 | C85 H136 N20 Na2 O24 S | 1852.9759 | 1852.9757 | 0.10 | 0.18 | 100.00 |
| VRAI | C85 H136 N20 O24 S | 1875.9670 | C85 H136 N20 Na O24 S | 1852.9775 | 1852.9757 | 0.97 | 1.79 | 99.52 |

**Compound S4**

*(4S,7S,10S,13S,16S,19S,22S,25S)-29-amino-7-(4-aminobutyl)-4-((S)-2-aminopropanamido)-19-((S)-sec-butyl)-25-carbamoyl-22-((S)-1-hydroxyethyl)-10,13,16-triisopropyl-5,8,11,14,17,20,23-heptaoxo-6,9,12,15,18,21,24-heptaazanonacosanoic acid*

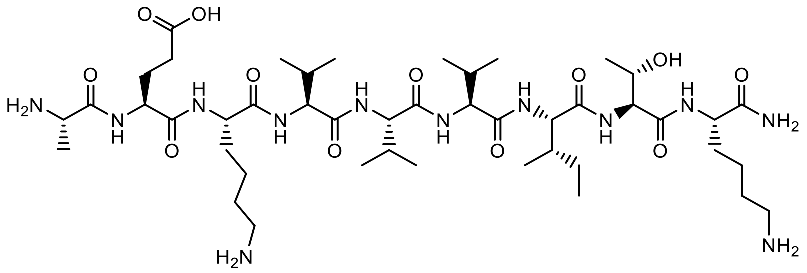

Peptide **S4** was synthesized accordingly to **General procedure B**.

The peptide was purified by semi-preparative RP-HPLC (Gradients of 5-47.2 % ACN in $H_2O$ containing 0.1% TFA in 13 min, retention time = 8.53 min, yield isolation: 10%).

**Molecular weight**: 984.6332 g/mol

**HRMS**: Calcd. for $[C_{45}H_{84}N_{12}O_{12} + H]^+$: m/z 985.6404 found 985.6431 $[M + H]^+$

**HPLC purity**: XSELECT column (C18, 2.1 x 75mm-2.5µm); (Gradients of 5-100% ACN in $H_2O$ containing 0.1% FA in 20 min); Rt = 9.472 min, 100%.

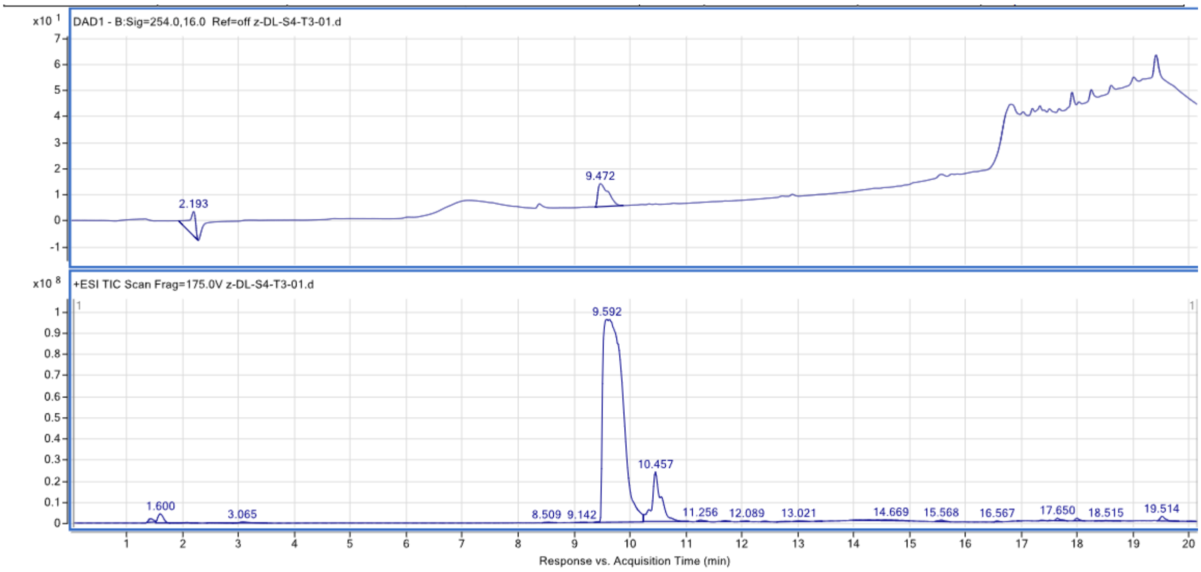


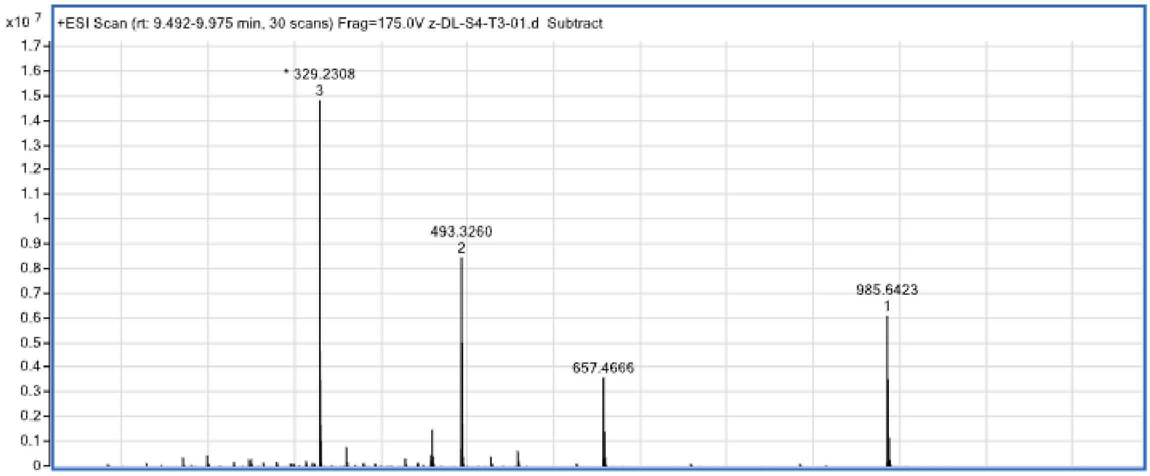

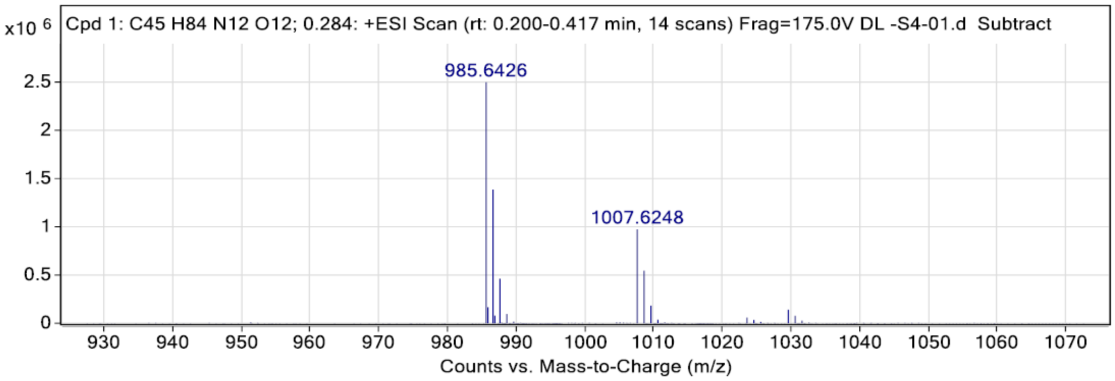


**Peak List**

| *m/z* | z | Abund | Target Molecular Formula | Target Ion Species | Calc. Mz | Rel. Diff. (ppm) |
|---|---|---|---|---|---|---|
| 985.6426 | 1 | 2491879.25 | C45H84N12O12 | (M+H)+ | 985.6404 | 2.2 |
| 986.6458 | 1 | 1382626.13 | C45H84N12O12 | (M+H)+ | 986.6433 | 2.5 |
| 987.6490 | 1 | 460691.44 | C45H84N12O12 | (M+H)+ | 987.6460 | 3.0 |
| 1007.6248 | 1 | 969610.88 | C45H84N12O12 | (M+Na)+ | 1007.6224 | 2.4 |
| 1008.6283 | 1 | 543614.56 | C45H84N12O12 | (M+Na)+ | 1008.6253 | 3.0 |

**Compound S6**

*(S)-N-((S)-1-amino-1-oxopropan-2-yl)-2-((S)-2-((2S,3S)-2-((S)-2-((S)-2-amino-3-hydroxypropanamido)-3-phenylpropanamido)-3-hydroxybutanamido)-3-methylbutanamido)-5-guanidinopentanamide*

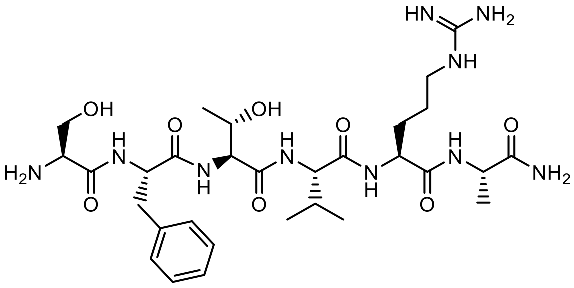

Peptide **S6** was synthesized accordingly to **General procedure B**.

was pure (no need to be purified)

Yield 100%

**Molecular weight**: 678.3826 g/mol

**HRMS**: Calcd. for [$C_{30}H_{50}N_{10}O_8$ + H]$^+$: m/z 679.3886 found 679.3895 [M + H]$^+$

**UPLC purity**: Waters Acquity UPLC apparatus equipped with a Luna Omega PSC18 Column (1.5 µM, 2.1 x 50 mm) coupled to a single quadrupole EDI-MS (Mictomass ZQ); (Gradients of 5-100% ACN in $H_2O$ containing 0.1% TFA in 5 min); Rt = 1.31 min, 100 %.

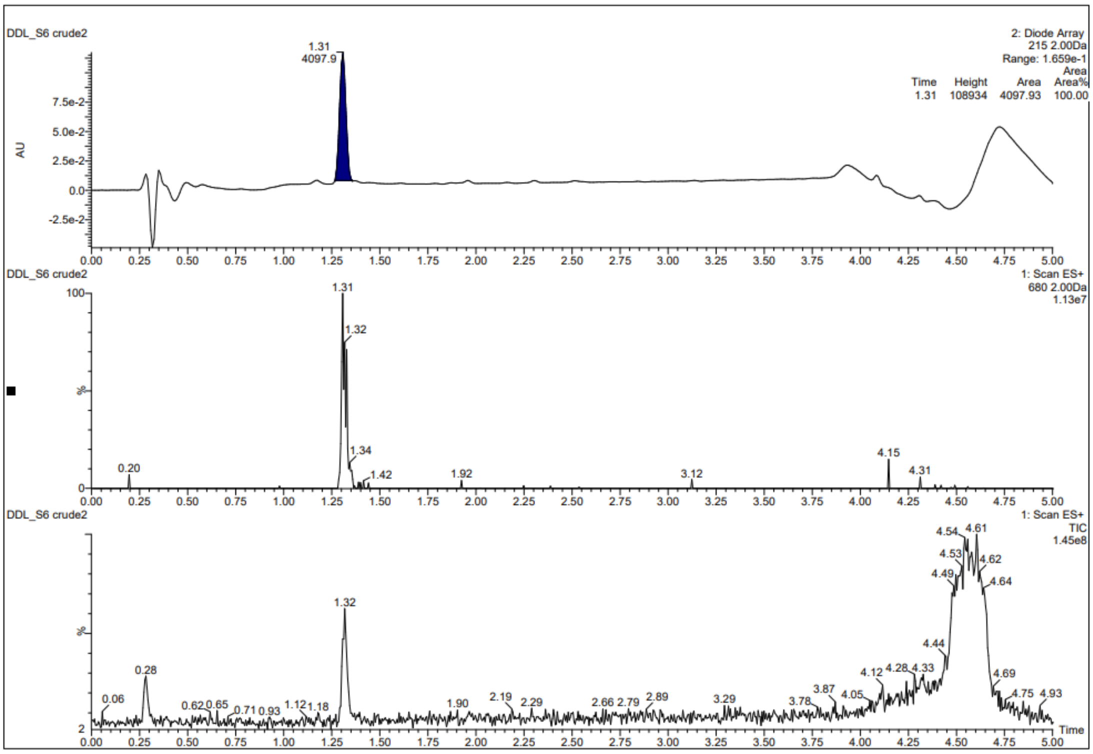
DDL_S6 crude2
2: Diode Array
215 2.00Da
Range: 1.659e-1
Time Height Area Area%
1.31 108934 4097.93 100.00
1.31
4097.9
AU
DDL_S6 crude2
1: Scan ES+
680 2.00Da
1.13e7
1.31
1.32
1.34
1.42
0.20
1.92
3.12
4.15
4.31
DDL_S6 crude2
1: Scan ES+
TIC
1.45e8
4.54
4.61
4.53
4.62
4.49
4.64
1.32
4.44
0.28
4.12
4.28
4.33
4.69
3.87
0.06
0.62
0.65
0.71
0.93
1.12
1.18
1.90
2.19
2.29
2.66
2.79
2.89
3.29
3.78
4.05
4.75
4.93
Time

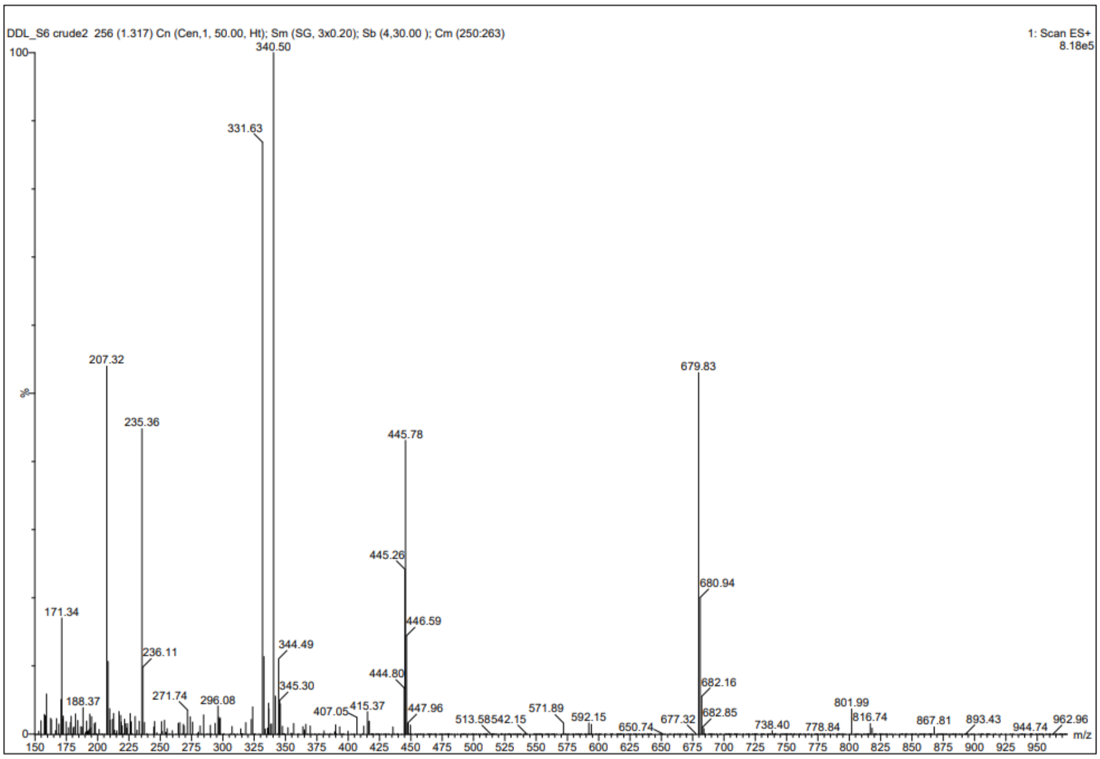
DDL_S6 crude2 256 (1.317) Cn (Cen,1, 50.00, Ht); Sm (SG, 3x0.20); Sb (4,30.00 ); Cm (250:263)
1: Scan ES+
8.18e5
340.50
331.63
207.32
679.83
235.36
445.78
445.26
680.94
171.34
446.59
236.11
344.49
444.80
682.16
345.30
188.37
271.74
296.08
407.05
415.37
447.96
513.58
542.15
571.89
592.15
650.74
677.32
682.85
738.40
778.84
801.99
816.74
867.81
893.43
944.74
962.96
m/z

| Spectrum Source | Fragmentor Voltage | Collision Energy | Ionization Mode |
|---|---|---|---|
| Peak (1) in "+ TIC Scan" | 175 | 0 | ESI |

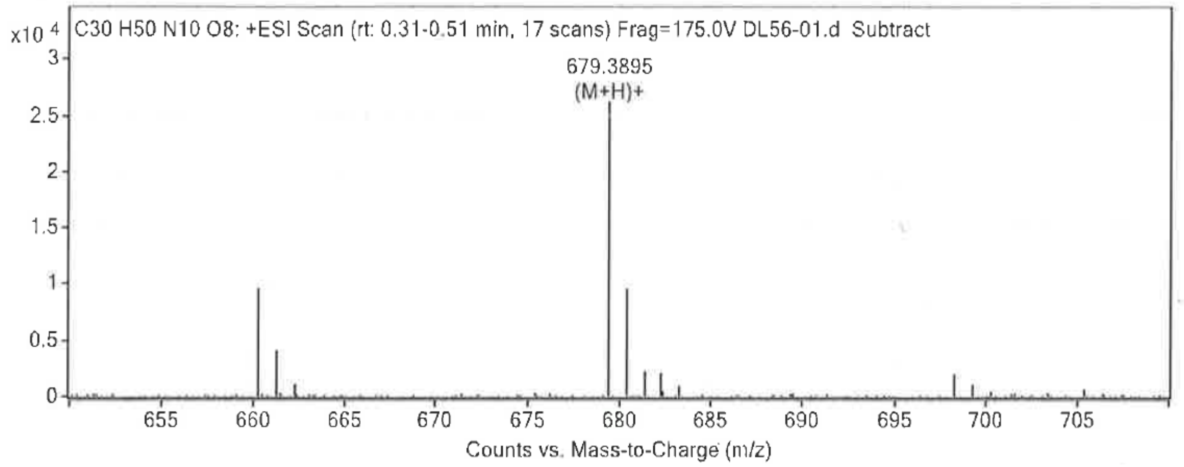


**Formula Calculator Results**

| Best | Generated Molecular Formula | Ion m/z | Generated Ion Formula | Calculated Molec Mass | Theorical Molec Mass | Rel Diff (ppm) | Diff (mDa) | Mass Match Probability |
|---|---|---|---|---|---|---|---|---|
| VRAI | C30 H50 N10 O8 | 679.3895 | C30 H51 N10 O8 | 678.3826 | 678.3813 | 1.86 | 1.26 | 99.16 |

**Compound S7**

*(S)-N-((3S,6S,9S,12S,15S)-12-((1H-imidazol-5-yl)methyl)-6-((S)-sec-butyl)-15-carbamoyl-9-isobutyl-2,17-dimethyl-4,7,10,13-tetraoxo-5,8,11,14-tetraazaoctadecan-3-yl)-6-amino-2-((2S,3S)-2-(2-((S)-2-amino-5-guanidinopentanamido)acetamido)-3-hydroxybutanamido)hexanamide*

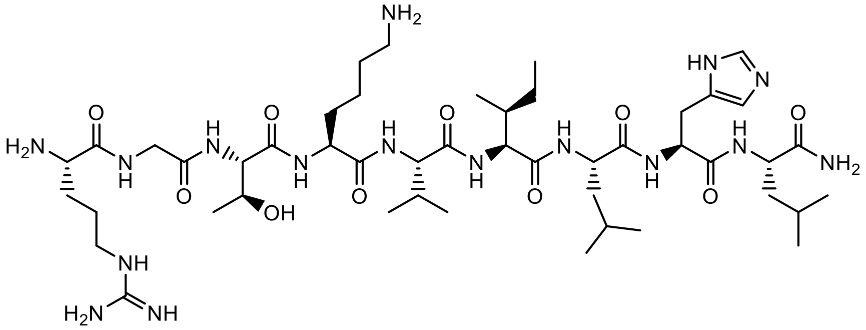

Peptide **S7** was synthesized accordingly to **General procedure B**.

The peptide was pure (no need to be purified)

Yield 100%

**Molecular weight**: 1034.6715 g/mol

**HRMS**: Calcd. for $[C_{47}H_{86}N_{16}O_{10} + H]^+$: m/z 1035.6715 found 1035.6792 $[M + H]^+$

**UPLC purity**: Waters Acquity UPLC apparatus equipped with a Luna Omega PSC18 Column (1.5 µM, 2.1 x 50 mm) coupled to a single quadrupole EDI-MS (Mictomass ZQ); (Gradients of 5-100% ACN in $H_2O$ containing 0.1% TFA in 5 min); Rt = 1.40 min, 100 %.

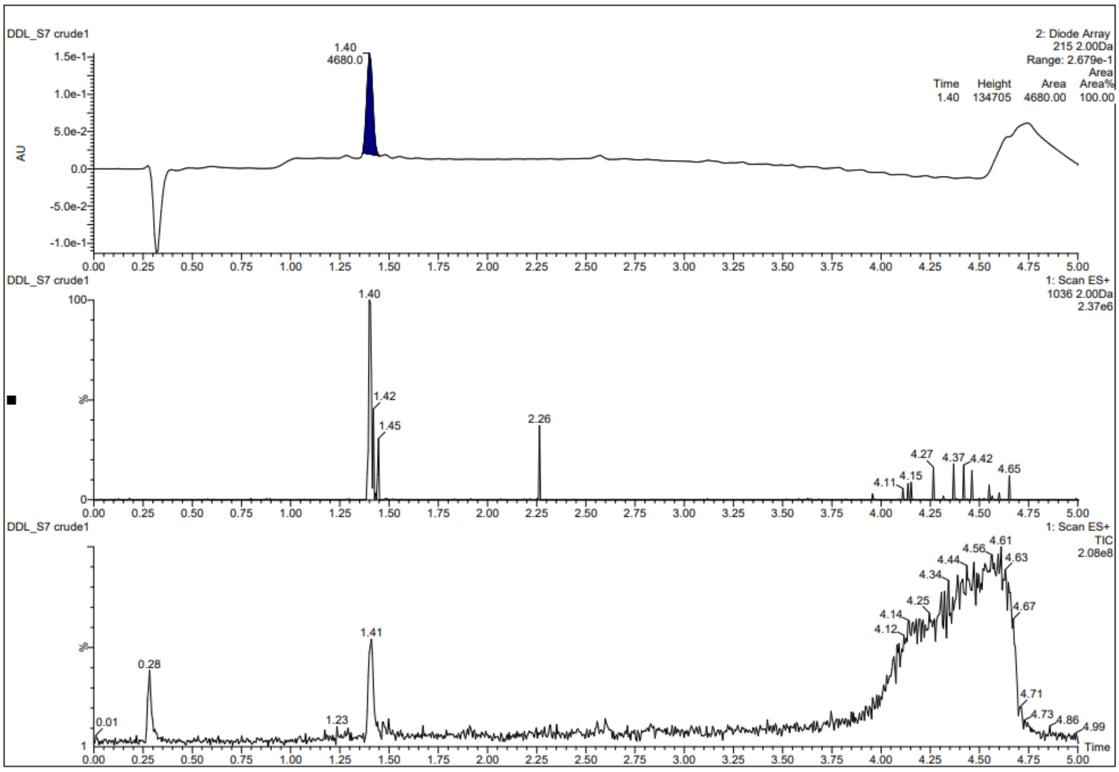
DDL_S7 crude1
2: Diode Array
215 2.00Da
Range: 2.679e-1
Time Height Area Area%
1.40 134705 4680.00 100.00
1.40
4680.0
AU
DDL_S7 crude1
1: Scan ES+
1036 2.00Da
2.37e6
DDL_S7 crude1
1: Scan ES+
TIC
2.08e8
Time

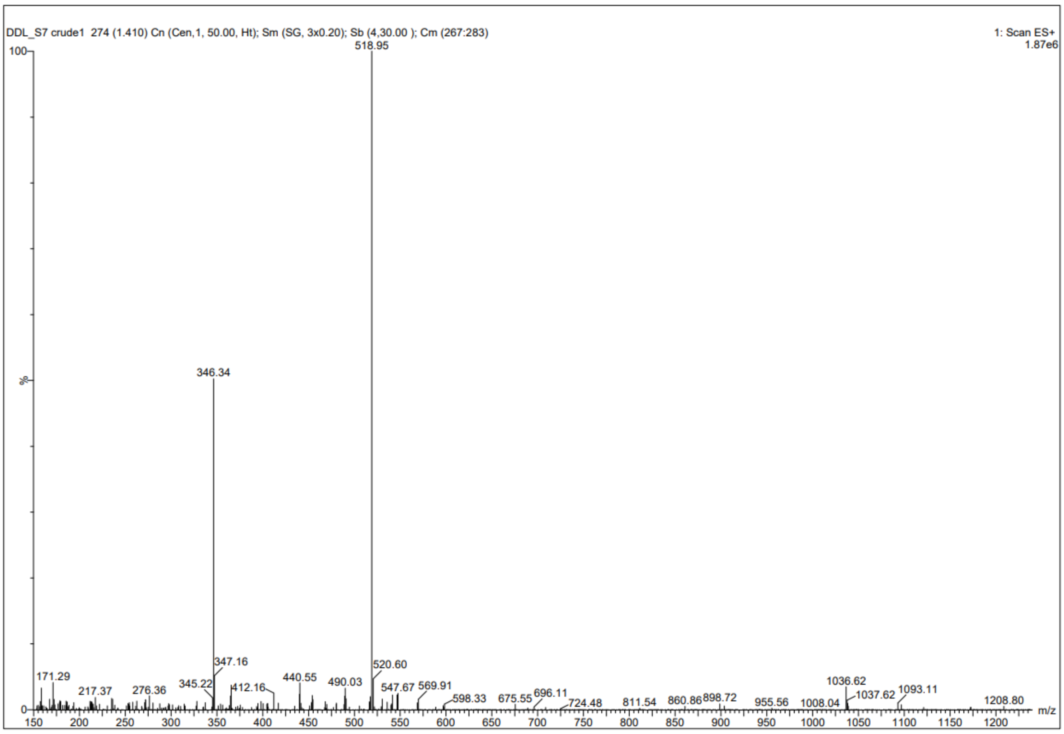
DDL_S7 crude1 274 (1.410) Cn (Cen,1, 50.00, Ht); Sm (SG, 3x0.20); Sb (4,30.00 ); Cm (267:283)
1: Scan ES+
1.87e6
518.95
346.34
m/z

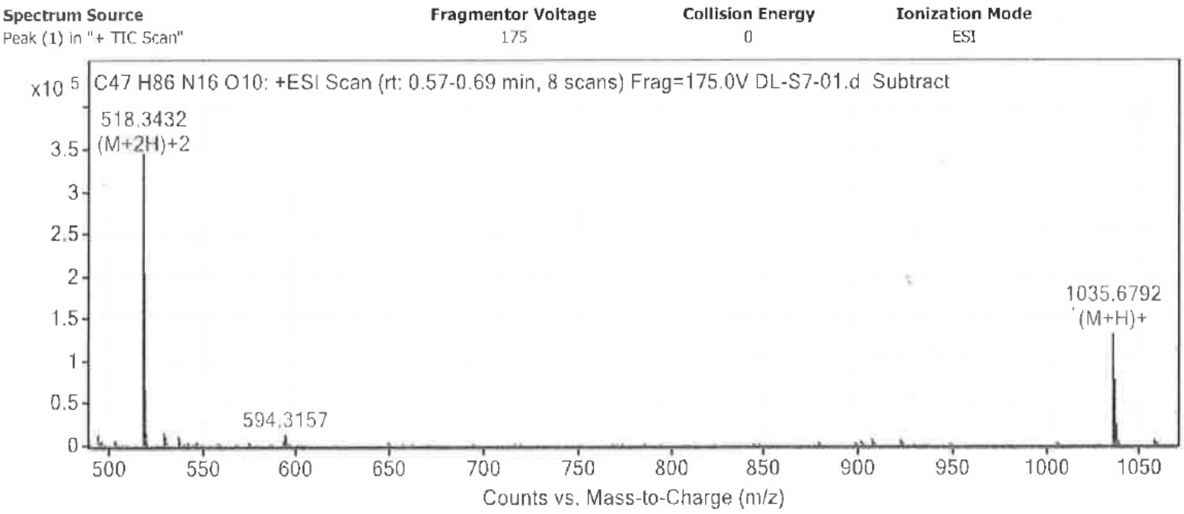


**Formula Calculator Results**

| Best | Generated Molecular Formula | Ion m/z | Generated Ion Formula | Calculated Molec Mass | Theorical Molec Mass | Rel Diff (ppm) | Diff (mDa) | Mass Match Probability |
|---|---|---|---|---|---|---|---|---|
| VRAI | C47 H86 N16 O10 | 518.3432 | C47 H88 N16 O10 | 1034.6715 | 1034.6713 | 0.22 | 0.23 | 99.99 |
| VRAI | C47 H86 N16 O10 | 1035.6792 | C47 H87 N16 O10 | 1034.6715 | 1034.6713 | 0.17 | 0.17 | 99.99 |

**Compound S4^short**

*(S)-2,6-diamino-N-((2S,3S,6S,9S,12S,15S)-6-((S)-sec-butyl)-3-carbamoyl-2-hydroxy-9,12-diisopropyl-16-methyl-5,8,11,14-tetraoxo-4,7,10,13-tetraazaheptadecan-15-yl)hexanamide*

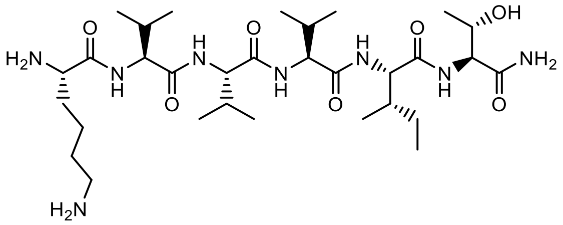


Peptide **S4^short** was synthesized accordingly to **General procedure C**.

The peptide was purified by semi-preparative RP-HPLC (Gradients of 5-100 % ACN in $H_2O$ containing 0.1% FA in 15 min, retention time = 7.09 min, yield isolation: 40%).

**Molecular weight**: 656.4585 g/mol

**HRMS**: Calcd. for $[C_{31}H_{60}N_8O_7 + H]^+$: m/z 657.4658 found 657.4675 $[M + H]^+$ Calcd. for $[C_{31}H_{60}N_8O_7 + Na]^+$: m/z 679.4486 found 679.4486 $[M + H]^+$

**HPLC purity**: Waters XBridge BEH300 (C18, 2.1 x 150mm-5µM); (Gradients of 5-100% ACN in $H_2O$ containing 0.1% TFA in 20 min); Rt = 10.65 min, 100%.

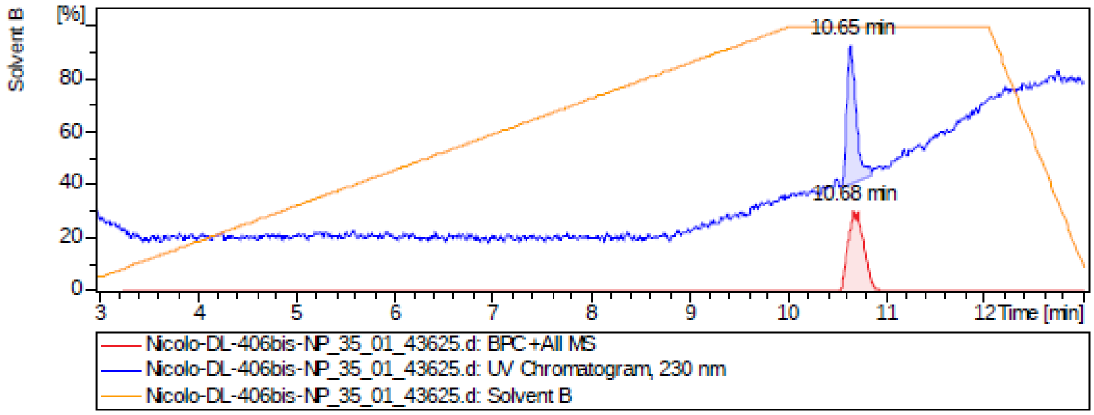


| # | RT [min] | Area | MW |
|---|---|---|---|
| Peak UV 230nm | 10.65 | 747.5 | 656.5 |

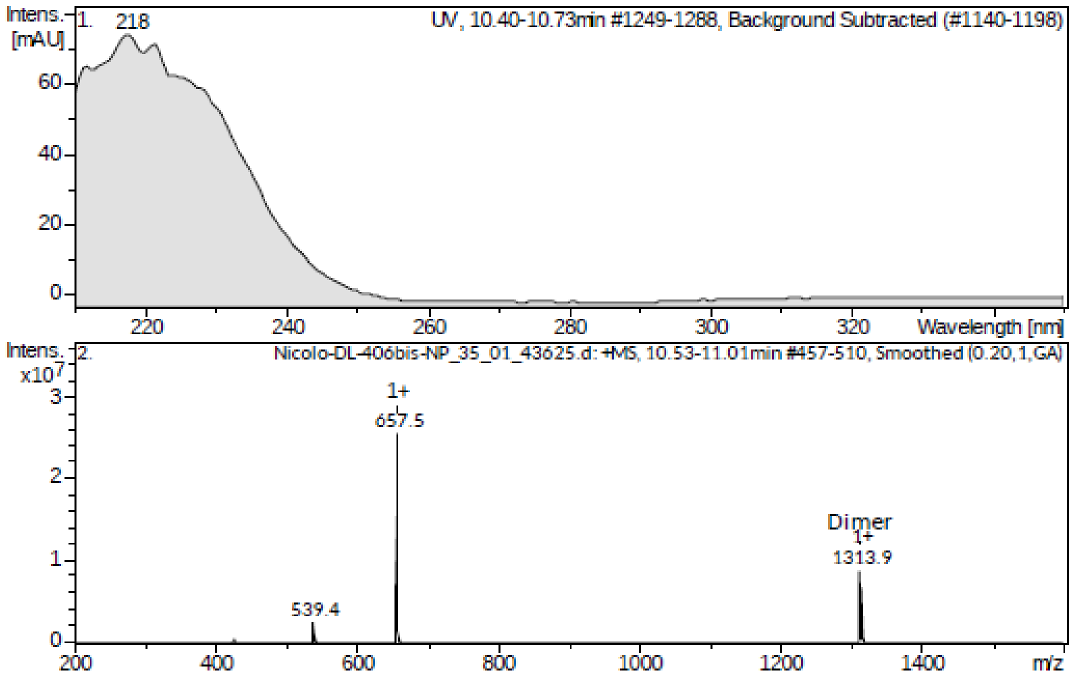

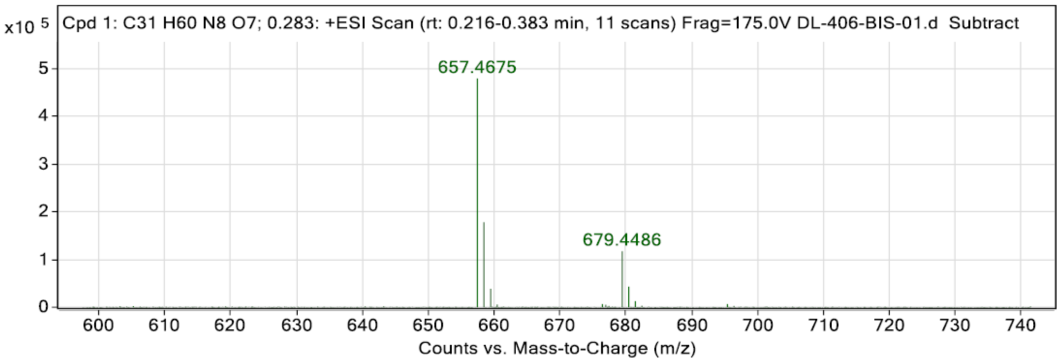


**Peak List**

| *m/z* | z | Abund | Target Molecular Formula | Target Ion Species | Calc. Mz | Rel. Diff. (ppm) |
|---|---|---|---|---|---|---|
| 657.4675 | 1 | 479228.94 | C31H60N8O7 | (M+H)+ | 657.4658 | 2.6 |
| 658.4700 | 1 | 177925.83 | C31H60N8O7 | (M+H)+ | 658.4687 | 2.0 |
| 659.4723 | 1 | 38352.61 | C31H60N8O7 | (M+H)+ | 659.4713 | 1.4 |
| 679.4486 | 1 | 117260.57 | C31H60N8O7 | (M+Na)+ | 679.4477 | 1.3 |
| 680.4510 | 1 | 42984.34 | C31H60N8O7 | (M+Na)+ | 680.4506 | 0.6 |

**Compound S7**$^{short}$

*(S)-6-amino-N-((S)-1-(((2S,3S)-1-(((S)-1-(((S)-1-amino-3-(1H-imidazol-5-yl)-1-oxopropan-2-yl)amino)-4-methyl-1-oxopentan-2-yl)amino)-3-methyl-1-oxopentan-2-yl)amino)-3-methyl-1-oxobutan-2-yl)-2-((2S,3S)-2-amino-3-hydroxybutanamido)hexanamide*

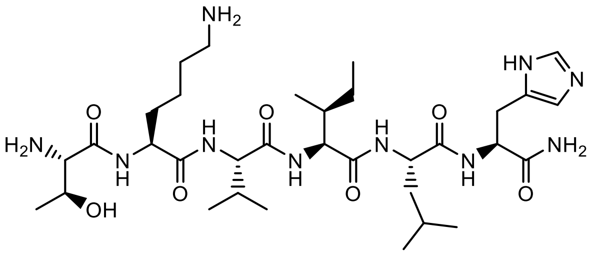

Peptide **S7**$^{short}$ was synthesized accordingly to **General procedure C**.

The peptide was purified by semi-preparative HPLC (Gradients of 5-100 % ACN in $H_2O$ containing 0.1% FA in 15 min, retention time = 6.10 min, yield isolation: 60%).

**Molecular weight**: 708.4646 g/mol

**HRMS**: Calcd. for [$C_{33}H_{60}N_{10}O_7$ + H]$^+$: m/z 709.4719 found 709.4721 [M + H]$^+$ Calcd. for [$C_{33}H_{60}N_{10}O_7$ + Na]$^+$: m/z 731.4569 found 731.4540 [M + Na]$^+$

**HPLC purity**: Waters XBridge BEH300 (C18, 2.1 x 150mm-5µM); (Gradients of 5-100% ACN in $H_2O$ containing 0.1% FA in 20 min); Rt = 6.69 min, 100%.

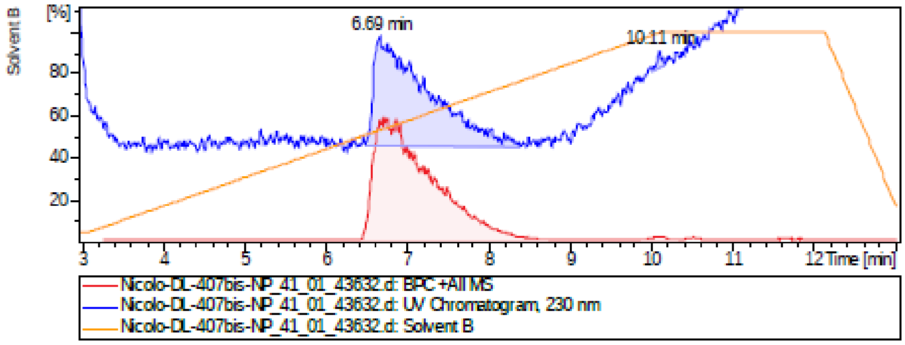


| # | RT [min] | Area | MW |
|---|---|---|---|
| Peak1 UV 230nm | 6.69 | 2101.7 | 708.5 |
| Peak2 UV 230nm | 10.11 | 26.4 | 709.5 |

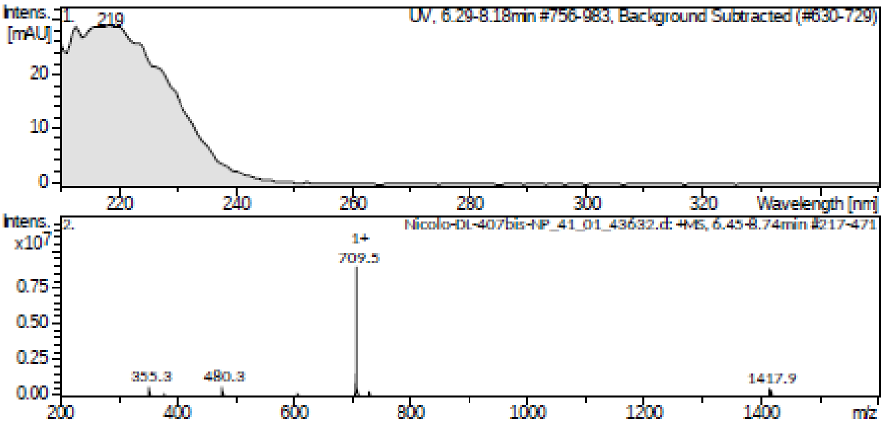


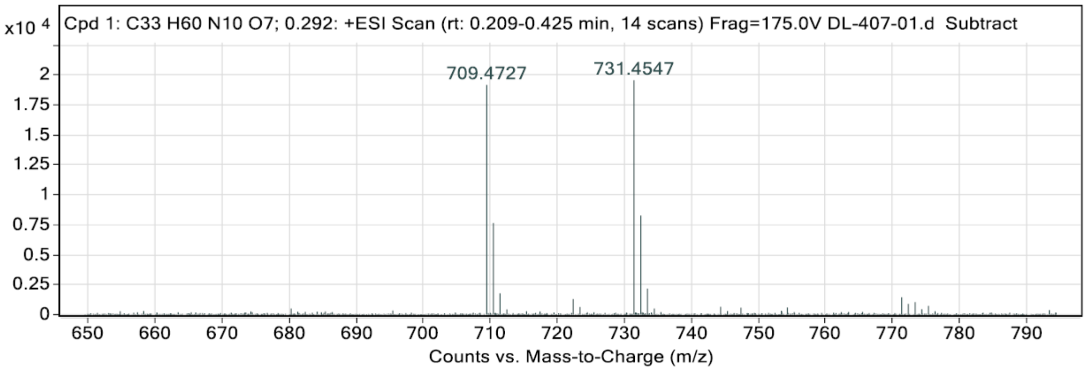


**Peak List**

| *m/z* | z | Abundance | Target Formula | Target Ion Species | Calculated m/z | Diff (mDa) | Diff (ppm) |
|---|---|---|---|---|---|---|---|
| 709,4727 | 1 | 19163,34 | C33H60N10O7 | (M+H)+ | 709,4719 | 0,74 | **1,04** |
| 710,4757 | 1 | 7644,27 | C33H60N10O7 | (M+H)+ | 710,4748 | 0,92 | **1,29** |
| 731,4547 | 1 | 19544,28 | C33H60N10O7 | (M+Na)+ | 731,4539 | 0,84 | **1,14** |
| 732,4583 | 1 | 8265,59 | C33H60N10O7 | (M+Na)+ | 732,4567 | 1,56 | **2,13** |
| 733,4595 | 1 | 2160,72 | C33H60N10O7 | (M+Na)+ | 733,4593 | 0,21 | **0,29** |

**Ac-PHF6*-NH$_2$**

Peptide **Ac-PHF6*-NH$_2$** was synthesized accordingly to **General procedure C**. Before the cleavage from the resin, the peptide was acetylated with acetic anhydride for 10 minutes.

The peptide was purified by semi-preparative HPLC (Gradients of 5-100 % ACN in $H_2O$ containing 0.1% FA in 20 min, retention time = 6.10 min, yield isolation: 78%).

**Molecular weight**:

**HRMS**: Calcd. for $[C_{34}H_{62}N_{10}O_9 + H]^+$: m/z 755.477 found 755.473 $[M + H]^+$ Calcd. for $[C_{34}H_{62}N_{10}O_9 + Na]^+$: m/z 777.459 found 777.460 $[M + Na]^+$

**HPLC purity**: XSELECT column (C18, 2.1 x 75mm-2.5µm); (Gradients of 5-100% ACN in $H_2O$ containing 0.1% FA in 15 min); Rt = 4.9 min, 100%.

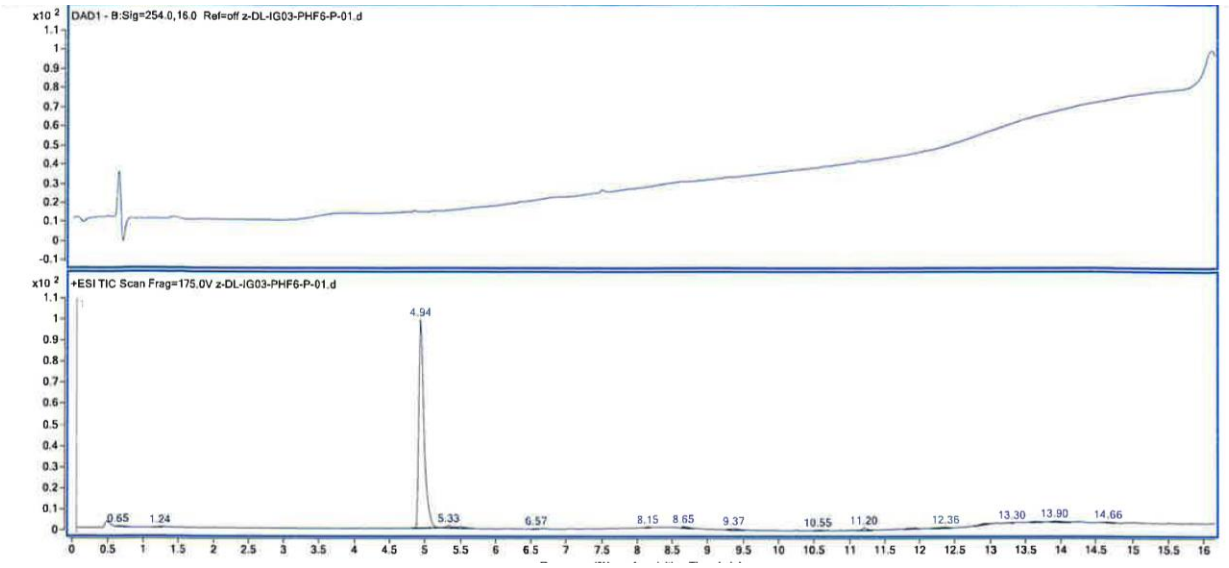

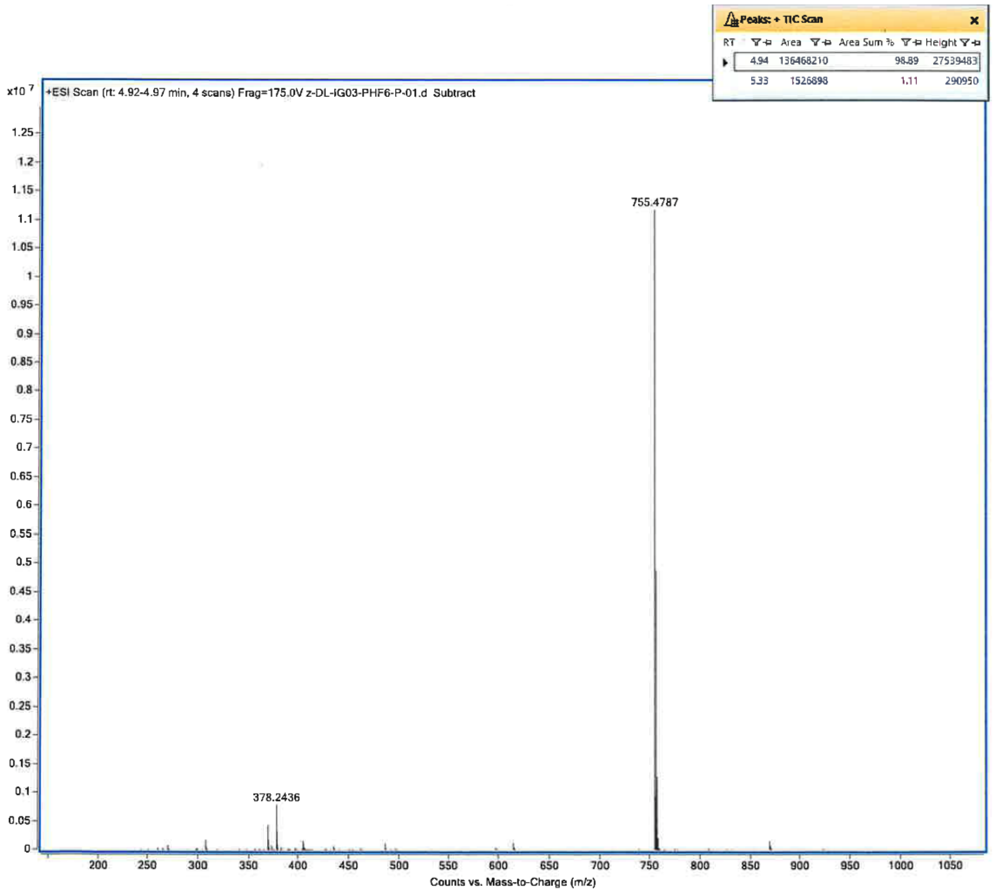
Peaks: + TIC Scan
RT Area Area Sum % Height
4.94 136468210 98.89 27539483
5.33 1526898 1.11 290950
x10 7
+ESI Scan (rt: 4.92-4.97 min, 4 scans) Frag=175.0V z-DL-IG03-PHF6-P-01.d Subtract
755.4787
378.2436
Counts vs. Mass-to-Charge (m/z)

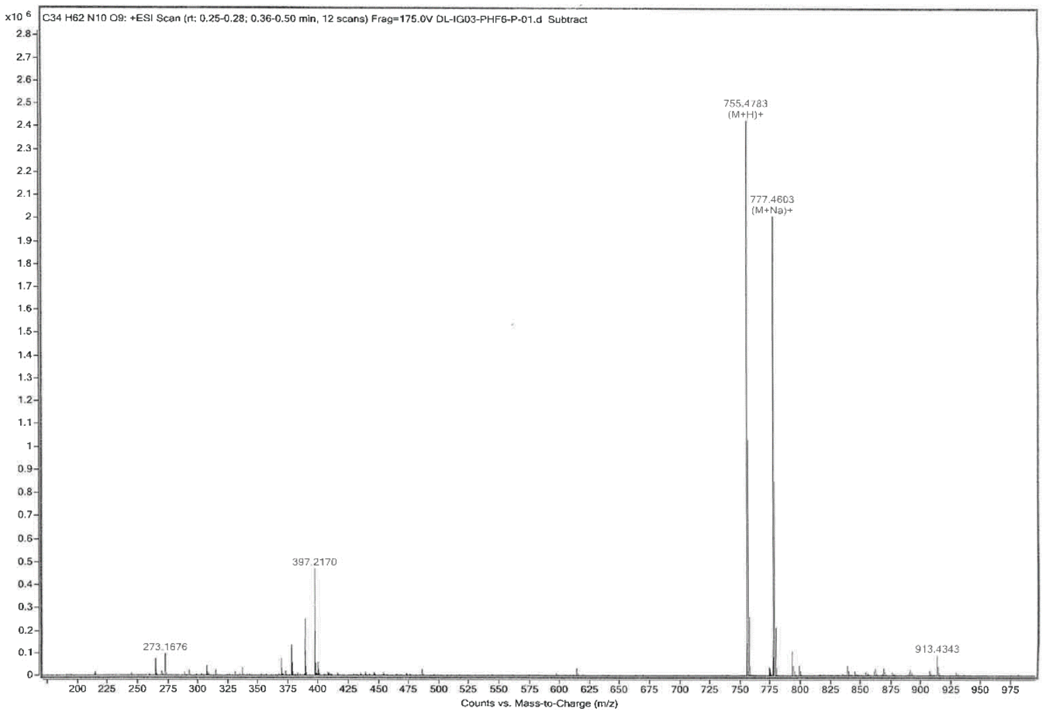
x10 6
C34 H62 N10 O9: +ESI Scan (rt: 0.25-0.28; 0.36-0.50 min, 12 scans) Frag=175.0V DL-IG03-PHF6-P-01.d Subtract
755.4783
(M+H)+
777.4603
(M+Na)+
397.2170
273.1676
913.4343
Counts vs. Mass-to-Charge (m/z)

| Spectrum Source | Fragmentor Voltage | Collision Energy | Ionization Mode |
|---|---|---|---|
| Peak (1) in "+ TIC Scan" | 175 | 0 | ESI |

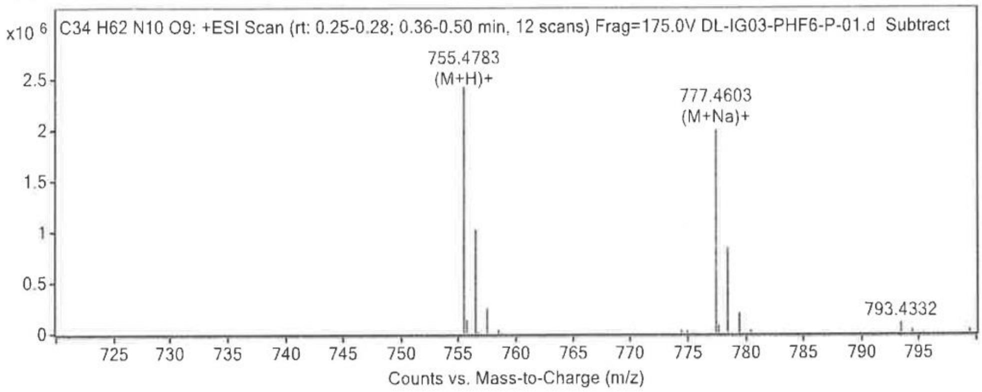


**Peak List**

| m/z | z | Abundance | Ion Species | Theorical Ion m/z | Rel Diff (ppm) | Diff (mAu) | Generated Molecular Formula |
|---|---|---|---|---|---|---|---|
| 389.2336 | 2 | 250915.89 | | | | | |
| 397.2170 | 2 | 469203.03 | | | | | |
| 397.7179 | 2 | 194761.16 | | | | | |
| 755.4783 | 1 | 2430046.00 | (M+H)+ | 755.4774 | 1.20 | 0.91 | C34 H62 N10 O9 |
| 755.7128 | 1 | 138197.22 | | 755.7110 | 2.42 | 1.83 | |
| 756.4816 | 1 | 1028225.75 | (M+H)+ | 756.4803 | 1.78 | 1.35 | C34 H62 N10 O9 |
| 757.4836 | 1 | 254129.64 | (M+H)+ | 757.4828 | 1.03 | 0.78 | C34 H62 N10 O9 |
| 777.4603 | 1 | 2009463.50 | (M+Na)+ | 777.4593 | 1.24 | 0.96 | C34 H62 N10 O9 |
| 778.4637 | 1 | 846120.50 | (M+Na)+ | 778.4622 | 1.88 | 1.47 | C34 H62 N10 O9 |
| 779.4653 | 1 | 208403.42 | (M+Na)+ | 779.4648 | 0.64 | 0.50 | C34 H62 N10 O9 |

**Formula Calculator Results**

| Best | Generated Molecular Formula | Ion m/z | Generated Ion Formula | Calculated Molec Mass | Theorical Molec Mass | Rel Diff (ppm) | Diff (mDa) | Mass Match Probability |
|---|---|---|---|---|---|---|---|---|
| VRAI | C34 H62 N10 O9 | 755.4783 | C34 H63 N10 O9 | 754.4711 | 754.4701 | 1.32 | 1.00 | 99.53 |
| VRAI | C34 H62 N10 O9 | 777.4603 | C34 H62 N10 Na O9 | 754.4712 | 754.4701 | 1.39 | 1.05 | 99.49 |

#### Circular Dichroism Spectroscopy

Peptidomimetics were dissolved in MQ water to a concentration of 500 µM as stock solutions. Before measurement, each compound was diluted to 125 µM concentration with 20 mM PB (pH 7.2) buffer or MeOH into a cuvette with a pathlength of 1 mm. The CD spectra were recorded on a J-815 spectropolarimeter (JASCO, Tokyo, Japan) from 190 to 260 nm at 20 and 37°C and a scan rate of 50 nm/min (accumulation n=3). Each CD spectrum was corrected by subtracting the corresponding baseline (PB buffer 20 mM or MeOH). Data processing was performed using Solver in Excel software (Microsoft). Deconvolution process was executed according to literature.[3]

**A**

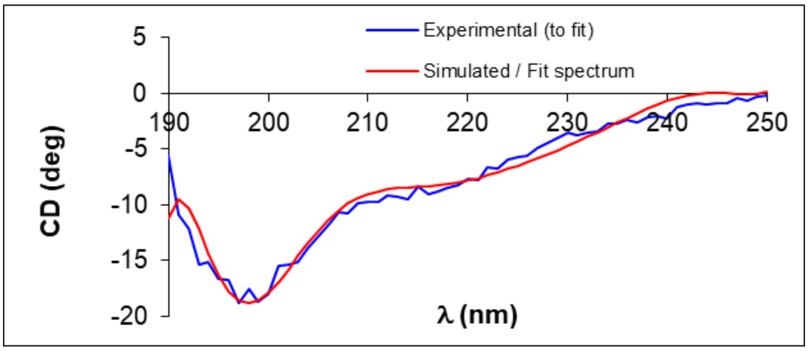


**B**

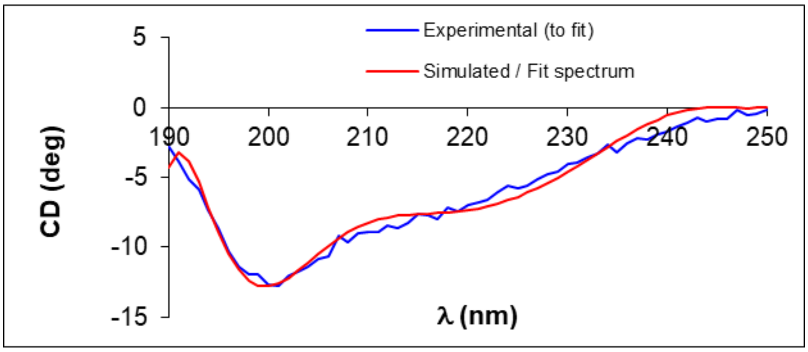


**Figure S1**: CD spectra of compounds **β-Tau (A)** and **β-Hsp90** (B) (125 µM) in PB (pH 7.4) at 20°C.

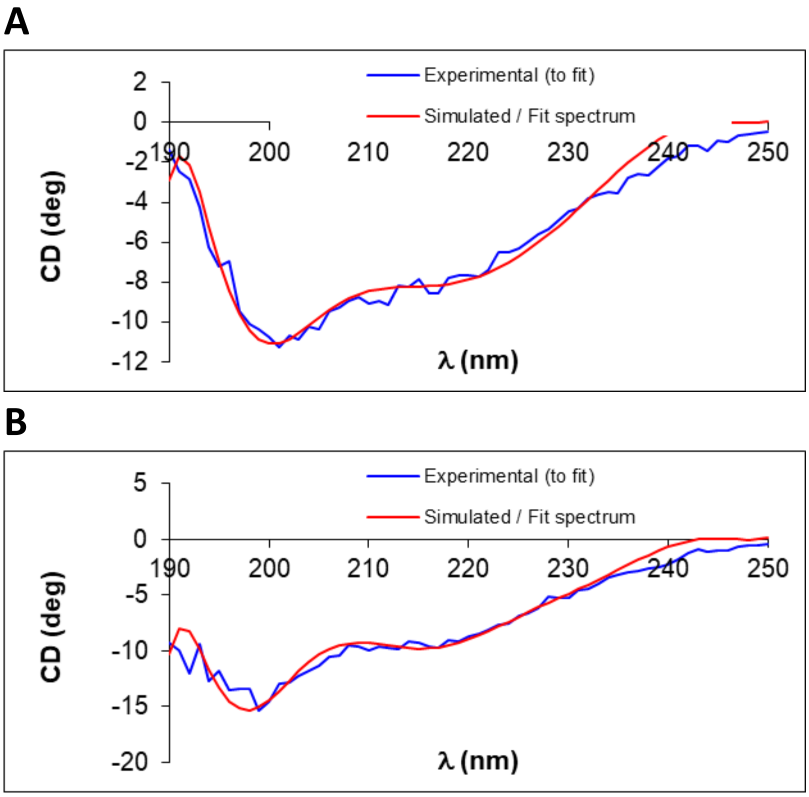


**Figure S2**: CD spectra of compounds **β-Tau (A)** and **β-Hsp90 (B)** (125 μM) in PB (pH 7.4) at 37°C.

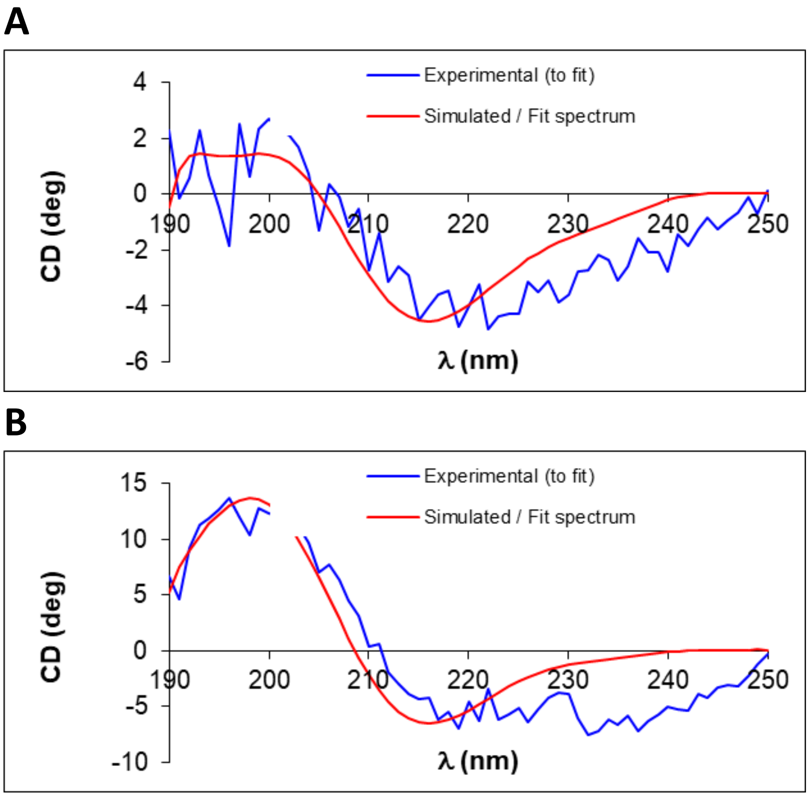


**Figure S3**: CD spectra of compounds **β-Tau (A)** and **β-Hsp90 (B)** (125 μM) in MeOH at 20°C

| Compound | Solvent | Temperature | Secondary Structure | | |
|---|---|---|---|---|---|
| | | | ***Helix (%)*** | **β-sheet (%)** | ***Others (%)*** ***(Random coil)*** |
| **β-Tau** | PB | 20°C | 7.2 | 32.4 | 60.4 |
| **β-Tau** | PB | 37°C | 4.6 | 39.5 | 55.9 |
| **β-Tau** | MeOH | 20°C | 0 | 63 | 37 |
| **β-Hsp90** | PB | 20°C | 12.1 | 29.5 | 58.5 |
| **β-Hsp90** | PB | 37°C | 11.7 | 32.3 | 56 |
| **β-Hsp90** | MeOH | 20°C | 0 | 83 | 17 |

**Table S1**: Summary of the result obtained from the deconvolution of the CD raw data
